# Understanding cryptic diversity within the honeypot ant species complex of *Myrmecocystus mendax*

**DOI:** 10.64898/2026.04.20.719579

**Authors:** Magnus Wolf, Nils Rensing, Hannah Neuhaus, Tobias van Elst, Ti H. Eriksson, Marek Borowiec, Philip S. Ward, Robert A. Johnson, Jürgen Gadau

## Abstract

Cryptic species diversity, overlooked due to extreme morphological similarity, is a common phenomenon among ants. The “honeypot ant” genus *Myrmecocystus* (Wesmael, 1838; Formicinae: Lasiini) likely features multiple cryptic species, as previously suggested by phylogenetic studies based on ultraconserved elements (UCEs). Here, this work is expanded upon by examining 140 specimens and 2,508 UCE loci, with a particular focus on the *M. mendax* species complex from the southwestern USA and northern Mexico. Phylogenomic and population genomic analyses revealed five distinct *M. mendax-*like lineages and identified two potential cases of cryptic species diversity, one within samples matching the morphology of *M. mendax* and another within samples conforming to *M. placodops*. Most specimens morphologically identified as *M. mendax* formed a well-supported monophyletic group sister to *M. melliger* assigned individuals, with evidence for ongoing hybridization between both species in the Madrean Sky Islands along the USA-Mexico border. Patterns in the main *M. mendax* clade also suggest adaptive divergence across ecological gradients, warranting further investigation. Overall, these findings highlight the power of UCE-based genomic data in phylogenetic reconstructions and population genetic analyses to better resolve cryptic species diversity, and clarify complex evolutionary histories shaped by introgression and incomplete lineage sorting.

## Introduction

The concept of cryptic species has been a part of the scientific discourse for almost 300 years (Bickford et al. 2007). The term generally describes a group of species that have been treated as a single nominal species due to their morphological indiscernibility and conflicts between species concepts (Bickford et al. 2007; Zúñiga-Reinoso and Benítez 2015). The development of modern sequencing techniques and high-resolution morphometry has allowed the delimitation of cryptic species with increased accuracy and confidence (Bickford et al. 2007; Zúñiga-Reinoso and Benítez 2015). While it is still uncertain whether specific taxa or ecosystems harbor more cryptic diversity than others (Bickford et al. 2007), Mayr (1963) predicted that cryptic species may be prevalent in organisms which rely mostly on olfaction to sense the environment and each other. As the human perception is predominantly visual, dissimilarities between such species are difficult to detect without the application of other objective methods (Mayr 1963; Seifert 2009).

Ants, eusocial insects which employ highly developed olfactory systems in collective social behavior (Ferguson et al. 2021) and nestmate recognition (Larsen et al. 2016), fit Mayr’s theoretical expectation for crypsis. Correspondingly, Seifert (2009) detected high levels of cryptic diversity in ants only indiscernible by particularly sensitive morphological analysis (NUMOBAT), e.g. in the genera *Lasius, Formica* and *Cardiocondyla*. He recognized intraspecific morphological as well as genetic polymorphisms and interspecific hybridization as biological drivers of crypsis, and suggested that some of the commonly used, single-locus genetic marker such as mitochondrial barcodes, are insufficient for species delimitation in ants.

*Myrmecocystus* Wesmael, 1838 (Formicidae: Formicinae: Lasiini) is a New World ant genus endemic to the western North America, including the United States, southern Canada and northern to central Mexico (Snelling 1976, 1982). These ants primarily inhabit arid and semi-arid habitats, and are popularly characterized by the production of replete workers (Snelling 1976), also called “honeypots” (Bodenheimer 1951) or “honey ants” (McCook 1882). Repletes are typically larger workers fed by nestmates to store fluids containing sugar, lipids and/or proteins in their greatly distended crop and abdomen, thus functioning as a communal storage units in times of reduced foraging activity (Burgett and Young 1974; Conway 1977). Repletism has been recorded in 20 of the 29 described *Myrmecocystus* species (Snelling 1976, 1982). However, it is not exclusive to the genus as at least seven other ant genera like *Camponotus, Melophorus, Leptomyrmex* and *Plagiolepis* produce repletes as well (Hölldobler and Wilson 1990; Nogueira et al. 2026). Species of *Myrmecocystus* also exhibit a diverse set of unusual social behaviors. The persistence of a queen foundress association referred to as primary polygyny was suggested for *M. mimicus* (Hölldobler et al. 2011) and demonstrated for *M. mendax* (Eriksson 2018). Moreover, non-violent strength assessments via ritualized display tournaments and subsequent brood raids resulting in dulosis (“slavery”) were recorded for *M. mimicus* (Hölldobler 1976) and *M. mendax* (Eriksson et al. 2019).

The taxonomic record of *Myrmecocystus* has been subject of debate and revisions since the 19th century (e.g. Cole 1936; Creighton 1950; Creighton et al. 1956; Emery 1894; Forel 1901; Gregg 1963; McCook 1882; Smith 1951; Wheeler 1908, 1913). Snelling (1976, 1982) provided the most recent revision of the genus, in which he included both extensive morphological descriptions as well as taxonomic keys. Doing so, he defined three subgenera, *Myrmecocystus s. str*., *Endiodioctes*, and *Eremnocystus*, containing 29 species (Snelling 1976, 1982). Snelling also recognized a strong similarity in the morphological patterns of *M. melliger, M. mendax* and *M. placodops* (*Endiodioctes*), placing them together in what he called the “*melliger group*” and describing complex variation in *M. mendax* which he hypothesized was due to character displacement (Snelling 1976). *M. mendax*, for example, was later also found to be polymorphic in terms of social structure and colony founding behavior, supporting the idea of cryptic diversity (Eriksson 2018; Eriksson et al. 2019).

More recently, van Elst et al. (2021) analyzed a set of genome-wide ultraconserved elements (UCEs) and published a comprehensive phylogenomic framework of *Myrmecocystus* which included 28 of the 29 described, as well as six undescribed species. Their findings both corroborated and conflicted with some aspects of Snelling’s observations. For example, *M. yuma*, originally placed in the *Eremnocystus* instead grouped with *Endiodioctes*, rendering both subgenera non-monophyletic. Furthermore, the phylogeny of van Elst et al. (2021) uncovered cryptic diversity for at least seven species, including *M. mendax* and *M. placodops*, and found two reciprocally monophyletic sister clades of *M. mendax* that could potentially represent separate species.

In the present study, the set of UCEs first established by van Elst et al. (2021) is examined to disentangle the phylogenetic and population genetic relationships within the cryptic diversity of the putatively cryptic *Myrmecocystus mendax* complex. In this work, the complex is defined as all samples keyed as *M. mendax* and those morphologically assigned to different species names but nested within the most expansive clade including all specimens described as *M. mendax* in van Elst et al. (2021). This corresponds to the *M. melliger* group of Snelling (1976), except that one species, *M. intonsus*, has not been sampled. The dataset presented here comprises a population-wide sampling of specimens distributed over the South-West USA and northern Mexico, totaling to 140 individuals and covering five previously recognized *Myrmecocystus* species closely related to *M. mendax*. An overview is provided about the existence of potentially undescribed species and population structures by assessing genetic isolation and differentiation. With this study, a guide to future taxonomic revisions supported by genomic data is provided.

## Results

### Sampling and UCE statistics

The initial dataset collected by van Elst et al. (2021) of 245 specimens and 2,519 UCEs, was filtered to a total of 140 specimens and 2,508 UCEs specific to the *M. mendax* species complex (see Supplementary Fig. S4-5 for filtering statistics and Fig. 1 for their geographic distribution). Apart from the limited number of individuals belonging to this species complex presented by van Elst et al. (2021), most of the here presented sequences have not previously been studied.

**Figure 1.**
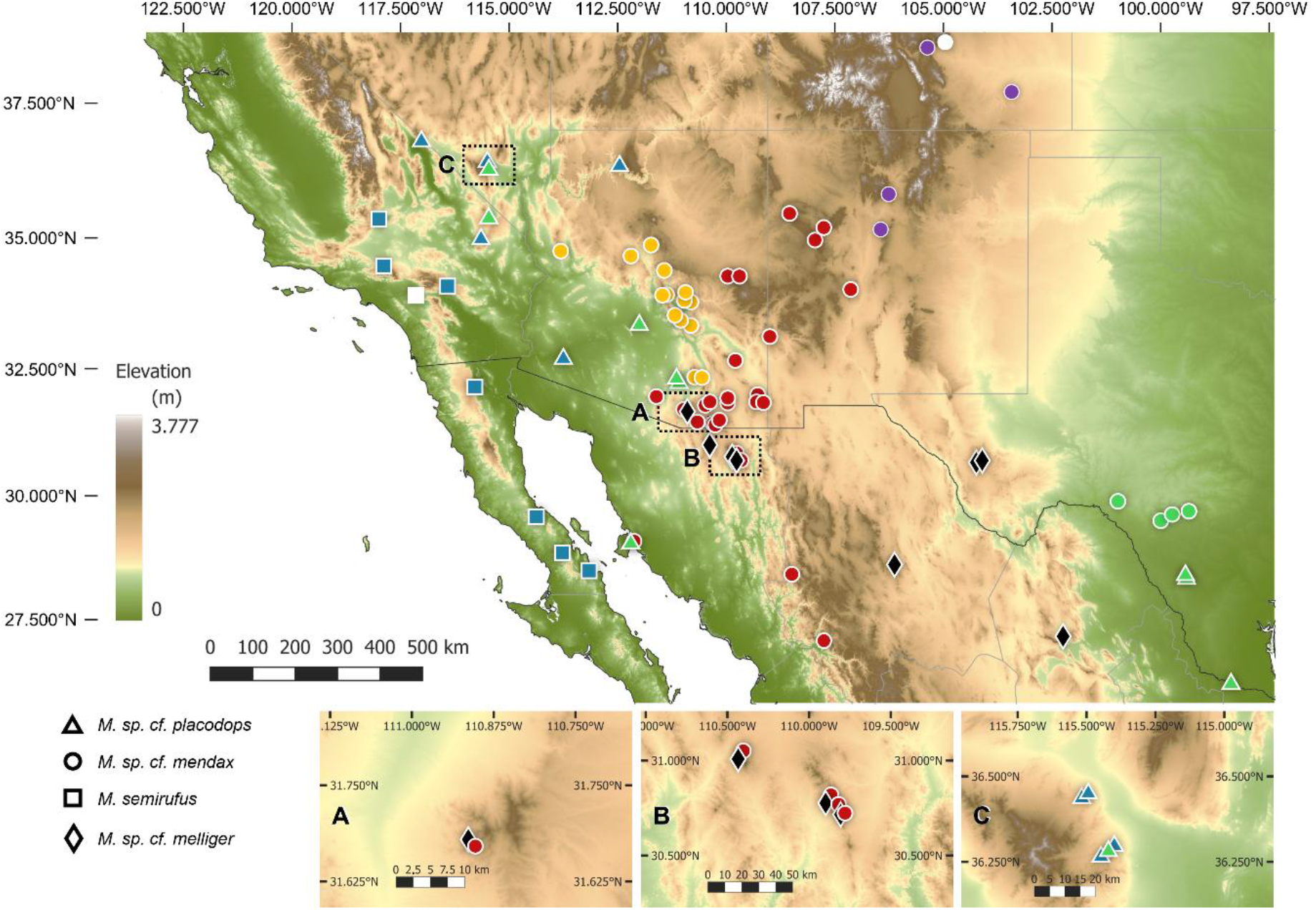
Geographic distribution of the analyzed samples of the *Myrmecocystus mendax* species complex. Coloration depicts cluster affiliation according to both the phylogenetic and population genetic analyses. Tag shape represents prior species assignments. White tags represent locations of type specimens for the respective species given in Creighton (1950). Map constructed in QGIS v.3.34 using the OpenTopology DEM Downloader v.3 add-on. Cases of close co-occurrence were further highlighted in magnified windows (A-C).

For phylogenetic inference, filtering and concatenating UCEs resulted in a total alignment length of 1,709,880 bp. The alignment incorporated sequences of 140 individuals and contained 159,334 informative sites from 2,092 UCE loci. The final partitioning scheme specified 2,023 subsets, which ranged in length from 146 to 3,197 bp and averaged at 845 bp. Accordingly, individual UCE alignments were between 146 bp and 2813 bp and contained an average of 76 bp informative sites.

For population genetics, raw reads of 139 samples were trimmed and mapped against all UCE loci extracted for the individual *Myrmecocystus_sp_cf_mexicanus*-02_RAJ_3061_ MEX_BC. The mean reference coverage depth per sample ranged from 4.06x to 140.01x, with a mean global average across all samples of 62.58x. After variant calling, the main VCF (Variant Call Format) file contained a set of 33,870 high-quality, biallelic SNPs.

### Phylogeny

The constructed phylogeny (Fig. 2) closely resembled the topology presented in van Elst et al. (2021) and exhibited high statistical support for most nodes. However, ultrafast bootstrap (UFB) values tended to decline towards the terminal nodes, particularly within the larger *M. mendax* clades. In total, 26 nodes showed UFB support below 95%. Quartet scores (QS), derived from comparisons between gene tree topologies and the final maximum likelihood species tree indicate a higher degree of topological conflicts across the tree, with a trend of higher amounts of conflicts at shorter branches (see pie charts in Fig. 2).

**Figure 2.**
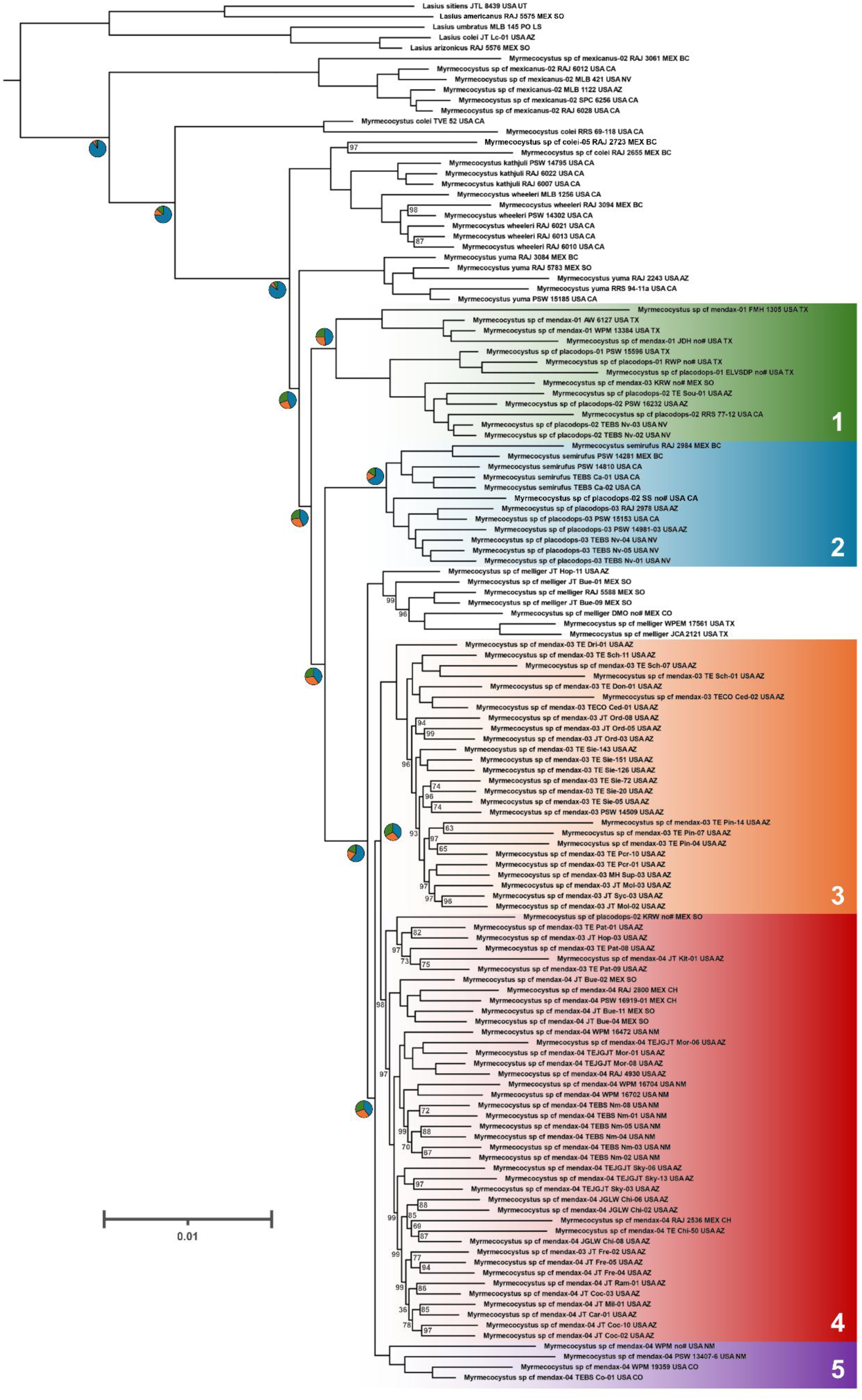
Maximum Likelihood phylogeny (cladogram) inferred from the concatenated and partitioned UCE sequence alignment. The node labels display support values generated from 1,000 ultrafast bootstraps. Clades labeled as clusters (1-5) represent the post-hoc population definitions that were input into population genetic analyses. Sample IDs were set according to van Elst et al. (2021). Quartet scores calculated with Astral-III are depicted as pie charts close to their respective nodes. The large number of alternative topologies (q2=green, q3=orange) suggest frequent conflicting signals in the individual UCE-trees compared to the overall species topology (q1=blue).

Within the samples morphologically assigned to the *M. mendax* species complex, six major monophyletic groups were identified, five of which included individuals identified as *M. mendax*. Colors used throughout the study reflect these clade assignments (cluster 1, green; cluster 2, blue; cluster 3, orange; cluster 4, red; cluster 5, violet; *M. melliger*, black). While the UFB values provided full (100%) support for the species complex clade (containing all *M. mendax-*assigned individuals and all nested relatives), the quartet scores were nearly evenly distributed among the three possible topologies, with a preference for the original species tree topology [q1=0.43, q2=0.3, q3=0.27].

The earliest diverging clade (cluster 1, green) comprised individuals identified as both *M. mendax* and *M. placodops*. There are two allopatric subclades, mostly comprising individuals identified as *M. mendax* and *M. placodops*, respectively, with the exception of one individual identified as *M. mendax* and nested within the *M. placodops* group. UFB support across this clade was consistently 100%, and the QS for the terminal branch indicated that nearly half of the gene trees supported the species tree topology [q1=0.48, q2=0.27, q3=0.25]. Another clade branching off subsequentially (cluster 2, blue) consisted primarily of individuals identified as *M. semirufus* and *M. placodops*, each forming distinct subclades. This clade also showed full UFB support and a high proportion of UCE trees consistent with the species tree topology [q1=0.67, q2=0.17, q3=0.16].

The remaining *M. mendax* individuals formed a monophyletic group comprising three distinct allopatric subclades (clusters 3+4+5; orange+red+violet), forming a sister lineage to a clade of individuals identified as *M. melliger*. While the branch at the split of *M. mendax* and *M. melliger* has a rather clear tendency towards the species tree topology q1 [q1=0.61, q2=0.2, q3=0.19], the quartet scores at the split of the remaining *M. mendax* subclades (clusters 3+4+5) reveal an almost equal frequencies for the species tree and alternative topologies [q1=0.39, q2=0.3, q3=0.3]. The smallest, more basal clade (cluster 5, violet) included five individuals and had full UFB support. The vast majority of *M. mendax* individuals were grouped into the two remaining clades (clusters 3+4, orange+red), which were also fully supported by UFB values, but again showed nearly equal frequencies across the three alternative topologies [q1=0.38, q2=0.29, q3=0.33]. Within these larger clades, UFB values tended to be lower at the terminal nodes.

### Phylogeography

In Texas, at the easternmost extent of the geographic distribution covered in this study, individuals from cluster 1 were recovered. Individuals identified as *M. mendax* were found at lower latitudes (Fig. 1) and in Edwards Plateau Riparian Forest, or - Limestone Savanna vegetation types (LANDFIRE 2022, EVT_CONUS, Fig. A1, dark gray). Some *M. placodops* individuals from cluster 1 were found in close geographic proximity to *M. mendax*, while others occurred much farther west, reaching as far as California. These specimens were found at similarly low latitudes, but in more scrub-like vegetation types (Fig. A1, Mojave-, Sonoran-, Tamaulipan Desert Scrub; light and dark brown, orange, dark blue). Additional individuals identified as *M. placodops* were recovered in the eastern cluster 2, with some individuals from clusters 1 and 2 sampled from the same mountain (Fig. 1C). Cluster 2 spans from northern Mexico into Baja California and is exclusively found in scrub deserts (Sonora-Mojave desert scrubs, Tab. A1). It includes a distinct species, *M. semirufus*, which forms a subclade sister to *M. placodops*-like individuals.

The large *M. mendax* clade (clusters 3, 4, and 5) is distributed across the transition zone between the Colorado Plateau (Colorado, Arizona and New Mexico), across the Madrean Sky Islands to the northern foothills of the Sierra Madre Occidental (Sonora and Chihuahua, Mexico) (Fig. 1). Although the ranges of these clusters are geographically close, they are separated by elevational and vegetation type gradients and the intervening desert habitat (Sonoran Paloverde-mixed cacti desert scrub, Fig. A1, orange). The northern distribution of cluster 4 is situated between clusters 3 and 5, effectively separating them. Cluster 4 is centered around the Madrean Sky Islands with a northward’s distribution into the Colorado Plateau. They occur mostly in Madrean Encinal, -Pinyon-Juniper woodlands as well as North American warm desert lower montane riparian woodlands vegetation types, but also reaching in the lower elevation shrub and scrub lands (Chihuanhuan Succulent desert scrub, Mogollon Chaparral, North American desert scrub, -woodland, Fig. A1, dark and light brown, Tab. A1). Cluster 3 is situated mostly around the Mogollon Rim north of Phoenix with similar vegetation types (Madrean Encinal, -Pinyon-Juniper woodlands). Cluster 5 is found in the northern areas of the Colorado Plateau within Southern Rocky Mountain pinyon-juniper woodland and closely situated inter-mountain Shrub-steppes. The southern portion of cluster 4 also partially overlaps geographically with several individuals of *M. melliger* (Fig. 1A, B) within the same Sky Island associated vegetation types.

### Population structure

Admixture analyses were performed on a subset of the population genetic dataset using the LEA R package (Gain and François 2021) in SambaR (Jong et al. 2021) with 2 to 20 ancestral *a priori* populations K, including clusters 1-5 and *M. melliger* (Fig. 3A, Tab. A2). Cross-entropy analysis across 200 simulations identified the lowest average cross-entropy at K = 12 (Fig. A4). However, since the cross-entropy curve flattened after K = 9 and no additional biologically meaningful structure emerged beyond this point, we considered K = 9 the most parsimonious model.

**Figure 3.**
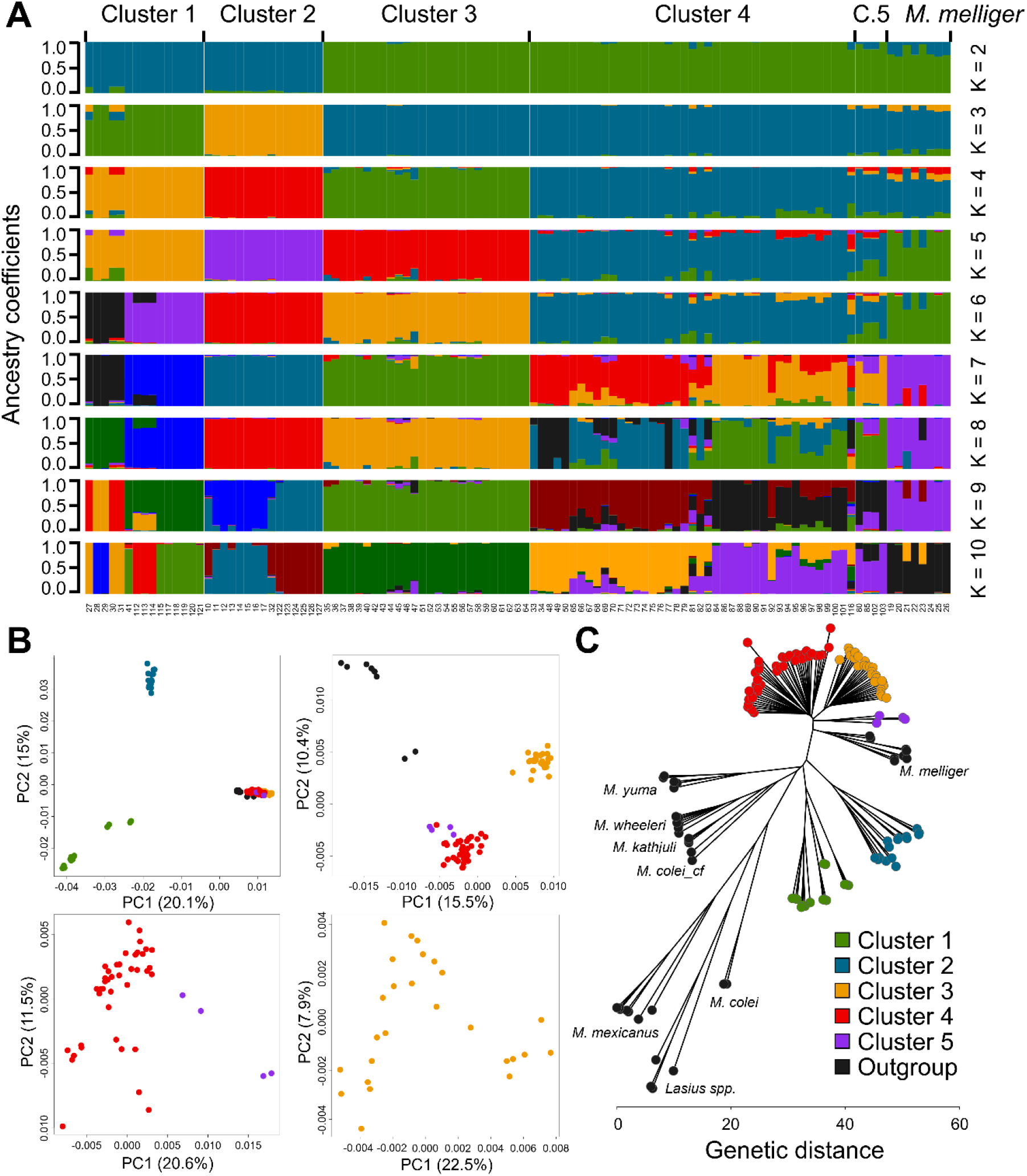
Population structure and genetic distance analysis of the *M. mendax* clusters defined in the phylogenetic analysis (Fig. 2), including *M. melliger* samples. *M. placodops* (cluster 1) and *M. semirufus* (cluster 2) are included in their respective cluster as well. **A** Admixture analysis across increasing numbers of *a priori* populations (K) performed using the LEA R package (Gain and François 2021). Colors indicate ancestry proportions of each individual (columns) and may not correspond to the colors used in other analyses. K=9 captured all biologically relevant clades identified in the phylogenetic analysis and was found to feature a low cross-entropy comparable to the global minimum of K=12. Numbers below the admixture plot represent individuals, their assignment can be found in Tab. A2. **B** PCoA performed on different subsets of *M. mendax* samples including *M. melliger* samples. Genetic distances were measured after Nei (1972). Upper left: Clusters 1 and 2 were distinctly separated from each other and the other groups. Upper right: *M. melliger* separates from clusters 3 to 5, except for two samples which had also shown shared ancestry with cluster 4 (see Fig. 3A). Clusters 4 and 5 are separated from cluster 3. Lower left: Cluster 4 and cluster 5 are not distinctly separated. Cluster 4 displays slight substructure. Lower right: Cluster 3 does not show any substructure. **C** Unrooted Euclidean distance-based phylogeny calculated from SNP data. The general topology was largely identical with the maximum likelihood phylogeny (Fig, 2). Some samples that showed clear signals of shared ancestry in the admixture analysis (Fig. 3A) possessed shortened or lengthened branches.

The most pronounced division in the admixture analysis separated clusters 1 and 2 from all other samples. These two clusters were subsequently distinguished from one another. Further partitioning separated cluster 3 from clusters 4, 5, and *M. melliger*. Within cluster 1, substructure separated individuals identified as *M. mendax* from individuals identified as *M. placodops*, while in cluster 4, two subgroups reflected the northern and southern portions of the geographic distribution. The northern subgroup of cluster 4 also showed affinity to the geographically adjacent cluster 5. Weaker substructure signals were observed within cluster 2, where *M. placodops* was genetically distinct from *M. semirufus*.

Several cases of shared ancestry were observed between groups. For example, two individuals from the *M. melliger* group (*M. melliger* JT Hop-11 USA AZ” and “*M. melliger* JT Bue-01 MEX SO”) share a large proportion of genetic components with cluster 4. These individuals coincide with cluster 4 individuals on a single Madrean sky island (Fig. 1A,B). Similarly, the northern subgroup of cluster 4 showed signs of shared ancestry with its southern counterpart. Geographically consistent patterns were also evident in cluster 1, where individuals identified as *M. mendax* shared genetic components with nearby individuals identified as *M. placodops*.

PCoAs (Principal Coordinates Analyses, Fig. 3B) were conducted on different sample subsets to resolve population structure on different scales using Nei’s genetic distance (Nei 1972). The largest PCoA (Fig. 3B, upper left) contained clusters 1 to 5 as well as *M. melliger* and showed great distances for clusters 1 and 2 to both each other and to all other groups. Removing clusters 1 and 2 from the PCoA (Fig. 3B, upper right) revealed a grouping of clusters 4 and 5, with cluster 3 separated. *M. melliger* was the most distant in this PCoA, except for two samples that had shown admixture with cluster 4 in the admixture analysis (Fig. 3A). When comparing just clusters 4 and 5 (Fig. 3B, lower left), cluster 4 was shown to separate into a larger group and a few smaller fractions. Cluster 3 displayed no further substructure (Fig. 3B, lower right).

A tree based on Euclidian genetic distance (Fig. 3C) was generated for the entire population genetic sample set. It showed the same general topology as the maximum likelihood tree (Fig. 2). Importantly, the Euclidian distance tree groups clusters 3 to 5 differently than the admixture analysis and PCoAs: the former infers clusters 3 and 4 as sister clades while the latter group clusters 4 and 5 together. However, the branch lengths in the distance-based phylogeny support the placement of clusters 1 and 2, which are clearly distant from all other clades in the *M. mendax* species complex. Patterns of shared ancestry in the admixture analysis were partially reflected in branch lengths of the distance-tree: both the samples of clusters 4 and 5 that appeared admixed with *M. melliger* in the aforementioned analyses differed noticeably in branch length from the rest of their respective group.

### Population differentiation

Genetic differentiation among the phylogenetic clusters 1 to 5 as well as the *M. melliger* group (Fig. 2) was quantified by calculating pairwise F_ST_ after Weir and Cockerham (1984), and significance was assessed through 1,000 permutations with a Wilcoxon rank-sum test (Fig. 4). To provide the context of clear species differentiation and clear panmictic populations, F_ST_ was also calculated for the more distant species *M. mexicanus* as well as for two random subsets of cluster 3. Furthermore, clusters 1 and 2 were split into two groups due to potential population substructure (Fig. 3).

**Figure 4.**
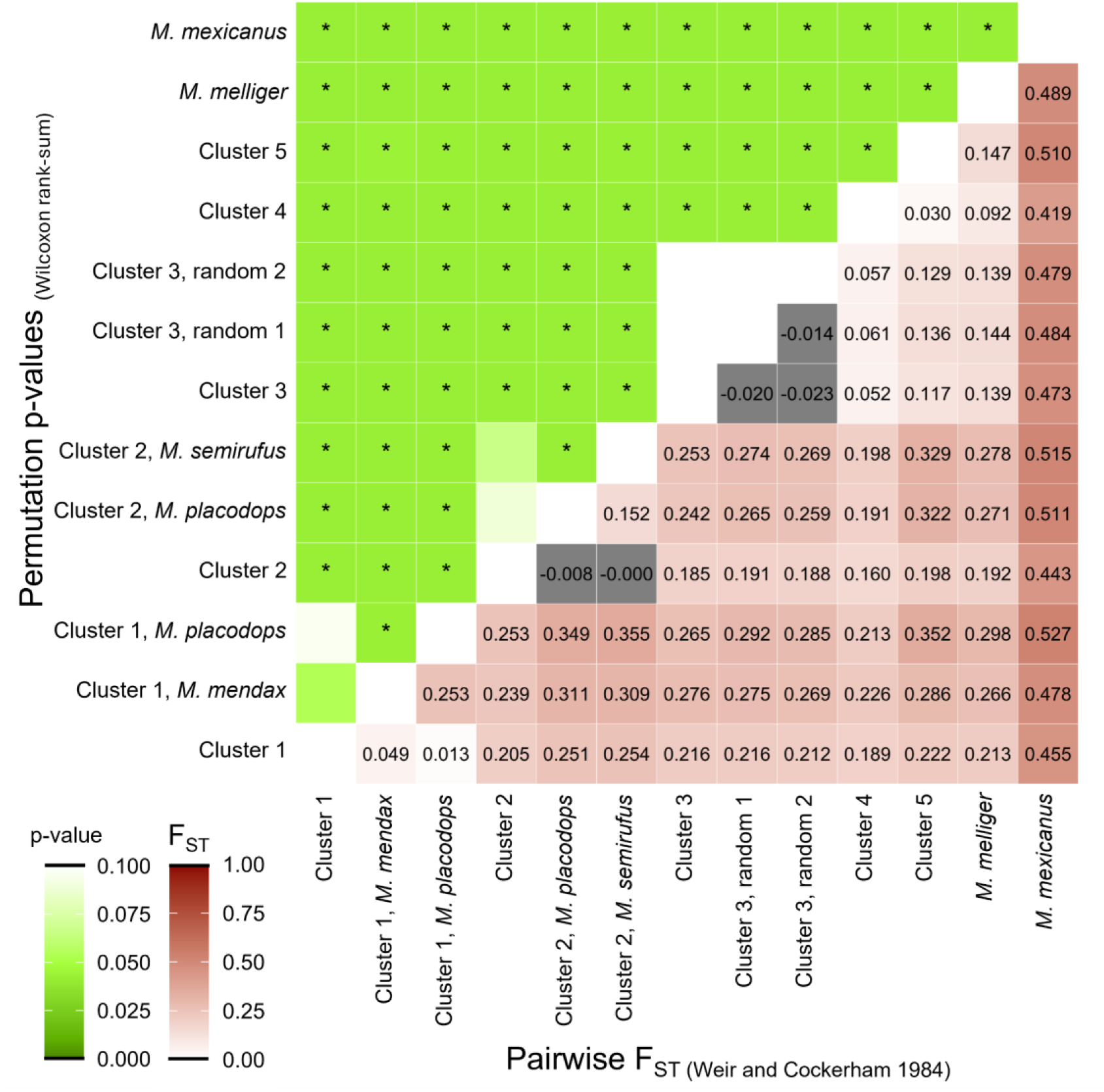
Pairwise F_ST_ estimates (Weir and Cockerham 1984) between phylogenetic clusters and the outgroup *M. mexicanus*. Clusters 1 and 2 were split into two groups respectively, due to potential population substructure. Within cluster 3, two random sub-populations were established to include a panmictic control group. P-values were calculated with a Wilcoxon pair-sum test following 1,000 pairwise permutations repetitions. Most F_ST_ values within the species complex were found to be between 0.16-0.3, including the sub-structures found in clusters 1 and 2. Lower values were only found in the comparisons of clusters 3 with 4 and 5 (0.05-0.14) or in the panmictic control which resulted in values around 0. Comparisons with the outgroup resulted in the highest values found, ranging from 0.42 to 0.53. Almost all F_ST_ values were found to be significant according to the permutation analysis with exceptions only found in the random control groups or in cases where substructures were compared to their own parent-cluster.

F_ST_ values were above 0.1 in most cases with notable exceptions found in the cross comparisons of clusters 3, 4 and 5 (F_ST_ = 0.05 for cluster 3 against 4; F_ST_ = 0.03 for cluster 4 against 5) as well as within the random control groups and the sub-structure groups of clusters 1 and 2. P-values were found to be significant in all cross-comparisons excluding the random control groups and the substructure versus parent cluster tests. The highest F_ST_ values were retrieved when testing against the outgroup *M. mexicanus* ranging from F_ST_ = 0.42 (cluster 4) to 0.53 (cluster 1, *M. placodops*).

Within the species complex, F_ST_ values also closely mirrored the approximated genetic distances depicted in the phylogenetic tree and the population genetic analyses (Fig. 2 and 3), with the *M. melliger* group for example being less differentiated from clusters 3, 4 and 5 (F_ST_ = 0.14 - 0.15) than from clusters 1 and 2 (F_ST_ = 0.19 - 0.21). Similarly, the sub-structures of clusters 1 and 2 returned rather high F_ST_ estimates when tested against each other (cluster 1: F_ST_ = 0.25 for individuals identified as *M. mendax* against *M. placodops;* cluster 2: F_ST_ = 0.15 for *M. semirufus* against individuals identified as *M. placodop*s) reflecting the higher genetic distances found in the phylogenetic and population genetic tests.

### Genetic diversity and hybridization

Genetic diversity, calculated as the UCE-wide rate of heterozygous sites in the called variants (HE, Fig. A2), returned the lowest average values for clusters 1 (0.125%) and 2 (0.119%) and the highest values for clusters 3 (0.176%) and 4 (0.179%). Cluster 5 (0.126%) and the *M. melliger* group (0.148%) fell in between those ranges. Notable outliers are one individual from cluster 1, “*M. mendax* 01 FMH 1305 USA TX”, with a UCE-wide HE of 0.449% as well as two individuals from the *M. melliger* group: “*M. melliger* JT Hop-11 USA AZ” with 0.302% and “*M. melliger* JT Bue-01 MEX SO” with 0.277%.

Since the two outliers from the *M. melliger* group had clear potential parent populations based on the admixture analysis and PCA (Fig. 3), a full-likelihood Bayesian hybrid inference approach (Chakraborty and Rannala 2023) was used to compare the likelihood of different hybridization scenarios: assignment as pure parent populations 1 or 2, as F1 or F2 hybrids, or as backcrosses to either of the parent populations (Fig. A3). Using linkage and recombination information based on 555 high-quality, phased SNPs retrieved from 32 UCE’s for both parent populations (*M. melliger* and *M. mendax* cluster 4) as well as the two hybrid individuals, individual “JT Bue01 MEX SO” showed the highest overall posterior probability (0.46) for the F1 hybrid scenario, while “JT Hop11 USA AZ” showed the highest probability (0.73) for a backcrossing scenario between an F1 hybrid and the *M. melliger* population.

## Discussion

The population-wide UCE dataset presented here for the *M. mendax* species complex supports earlier findings of cryptic diversity within the clade (Snelling 1976; van Elst et al. 2021). Evidence for cryptic speciation was identified in two formally described species, *M. mendax* and *M. placodops*, both of which appeared polyphyletic in the phylogenomic maximum-likelihood tree and separated into multiple distinct groups in the population-genomic analyses. Within individuals identified as *M. mendax*, the largest clade was further divided into two to three clearly distinct groups with distinct but adjacent geographic distributions. *M. melliger* emerged as the closest relative of this clade, with signals of ongoing admixture suggesting the presence of a potential hybridization zone along the U.S. - Mexico border in the Madrean sky islands.

### Cryptic speciation in *M. mendax*

The existence of multiple species in the taxon known as *M. mendax* was already presaged by Snelling (1976) based on variation in hair length. He identified a predominantly northern short-haired form, as well as an exclusively southern long-haired form, although he considered them as one species due to a geographic cline (Snelling 1976). Results of Eriksson (2018) partially conflicted with Snelling’s findings, as long-haired *M. mendax* were also collected in modern northern populations. Nevertheless, the overall extent of the polymorphism was affirmed by Eriksson (2018). Later, Eriksson et al. (2019) also found *M. mendax* to be polymorphic in terms of social structure and colony founding behavior, further supporting the notion of hidden biological diversity. The first comprehensive molecular phylogeny for the genus *Myrmecocystus* by van Elst et al. (2021) added more evidence to the presence of cryptic diversity in *M. mendax* with limited taxon sampling.

The present UCE-based phylogenomic and population genomic analysis confirmed the polyphyletic nature of specimens assigned to *M. mendax*. Most specimens cluster together as a monophyletic group (Fig. 1+2, clusters 3,4,5) while a couple of individuals identified as *M. mendax* grouped with *M. placodops* individuals (Fig. 1+2, cluster 1). This latter *M. mendax / M. placodops* clade showed high UCE-based quartet score support (Fig. 2), a very clear by-species separation in the population genetic analyses (Fig. 3) and considerably higher F_ST_ values towards all other clusters (F_ST_ = 0.189-0.254, Fig. 4). Specimens in cluster 1 identified as *M. mendax* are all located in southern Texas at lower latitudes (Fig. 1) and in “Edwards Plateau Riparian Forest, or - Limestone Savanna” vegetation types (LANDFIRE 2022, EVT_CONUS, Fig. A1, dark gray) while being grouped together with *M. placodops* specimens of wider distribution range, at similarly low latitudes, but more scrub-like vegetation types (Fig. A1, Mojave-, Sonoran, Tamaulipan Desert Scrub; light and dark brown, orange, dark blue) compared to its *M. mendax* identified relatives. There is furthermore a strong phylogenetic and population genetic support for an additional separation of the *M. mendax*-like and *M. placodops*-like specimens within this cluster, especially considering the high F_ST_ value between both subgroups (Fig. 4, F_ST_ = 0.253). Nevertheless, more extensive sampling will be required to draw definitive conclusions about the extent of their divergence.

However, the two major phylogenetic *M. mendax* groups (cluster 1 vs 3,4,5) and clusters 3 and 4 in particular (see below), were distributed West-to-East instead of North-to-South, conflicting with the hair length cline first described by Snelling (1976). Additional measurements by Eriksson (2018) showed that the hair-length cline may also not be as pronounced as originally thought. Consequently, this morphological trait may not be a sufficient indicator of species assignment or differentiation.

### Cryptic speciation in *M. placodops*

Apart from the individuals identified as *M. placodops* present in cluster 1, some specimens assigned to this species group together with specimens of *M. semirufus* in cluster 2. This polyphyletic and hence cryptic nature of *M. placodops* was also previously reported by van Elst et al. (2021) with limited taxon sampling. Intriguingly, these *M. placodops*-like and *M. semirufus* samples of cluster 2 display a clear allopatric geographic separation, but their general distribution is mostly limited to the western-most part of the sampling along the Mexican-Californian Baja Peninsular ranges and are exclusively found in desert scrub vegetation types (Sonora-Mojave, Fig. A1, Tab. A1), pointing towards a different habitat usage for cluster 2 compared to cluster 1 as well as the larger *M. mendax* (cluster 3,4,5) + *M. melliger* clade. Interestingly, Helms and Helms Cahan (2012) previously found genetic differentiation between two social forms of *Veromessor pergandei* in the same area which they hypothesized may be explained by a past isolation during glacial periods.

As already discussed for cluster 1, all analyses point to a strong support for cluster 2. The genetic distance between both clusters (1 and 2) containing *M. placodops*-like individuals is also comparable or even higher compared to their distances towards the other large cluster (*M. mendax* cluster 3,4,5 + *M. melliger*), as indicated by the PCoA analysis (Fig. 3B), the genetic distance-based NJ tree (Fig. 3C) and the high F_ST_ values between (Fig. 4) clusters 1 and 2.

Similar to what was found in cluster 1, specimens of cluster 2 can be separated into two distinct subclades composed mainly of *M. placodops*-like individuals and *M. semirufus*. The two individuals formerly described as *M. mendax* or *M. koso* in van Elst et al. (2021) grouping together with *M. placodops* were omitted from further evaluation since they showed high proportions of missing data and hence long branches in the phylogenetic tree (Fig. 2). Signals of genetic substructure in cluster 2 were clear but weaker compared to cluster 1 substructures, although internal branches were shorter in cluster 2 (Fig. 2.) and showed less pronounced separation in the PCoA analysis (Fig. 3B), indicating a more recent lineage divergence. Given the more restricted geographic range of cluster 2, a possible explanation for their more recent divergence might be a glacial desert scrub refugia within the Baja Peninsula range which led to a post-glacial distribution into continental scrub lands and isolation-by-distance as observed for other arthropod species like the wolf spider (*Pardosa sierra*) (González-Trujillo et al. 2016) and the associated vegetation types in general (Holmgren et al. 2011). However, the currently small sampling sizes prevents more conclusive interpretations of allopatry, warranting further investigation.

### Gene flow in the *M. mendax / M. melliger* clade

All *M. mendax* individuals in clusters 3, 4, and 5, together with the included *M. melliger* samples, form a monophyletic group strongly supported by both the phylogenomic quartet score analyses (Fig. 2) and the population genetic results, including admixture, PCA, and genetic distance analyses (Fig. 3). This finding is consistent with the results of van Elst et al. (2021), but contrasts with an earlier partial mtDNA based-phylogeny (Kronauer et al. 2004). Despite the strong support for this monophyletic group, the internal relationships among its members remain comparatively difficult to resolve, as indicated by ambiguous quartet scores, moderate signs of shared ancestry, and relatively even genetic distances. Together, these patterns suggest a strong influence of incomplete lineage sorting (ILS) and gene flow. Nonetheless, both the maximum-likelihood phylogenomic tree (Fig. 2) and the genetic distance–based NJ tree (Fig. 3C) support an initial split between *M. melliger* and the *M. mendax* clusters 3-5. Although all pairwise F_ST_ values are relatively low compared to others in the data (Fig.4), those between *M. melliger* and the *M. mendax* clusters tend to be slightly higher. This supports their divergence as the most ancient split within this monophyletic group (Fig. 4). Notably, cluster 4 shows the lowest F_ST_ values toward all other groups, likely reflecting its geographically central position.

While evidence of admixture (Fig. 3A), overlapping distributions in the Madrean Sky Islands (Fig. 1A, B), and low F_ST_ values (Fig. 4) may indicate ongoing hybridization and genetic exchange between *M. mendax* (clusters 3,4 and 5) and *M. melliger*, it remains difficult to distinguish these signals from incomplete lineage sorting (ILS) and post-divergence introgression close to their divergence. Nevertheless, two *M. melliger* individuals (“*M. melliger* JT Hop-11 USA AZ” and “*M. melliger* JT Bue-01 MEX SO”) show strong signals of admixture with cluster 4 and group more closely to cluster 4 and 5 than to conspecifics in the PCoA analysis. These trends may represent recent hybridization events. Both admixed *M. melliger* individuals were collected on the same Sky Islands inhabited by *M. mendax* cluster 4 as well (Fig. 1A, B), providing the necessary geographic proximity given the otherwise isolating nature of these mountain ranges for other ant species (Favé et al. 2015). This interpretation is further supported by exceptionally high levels of heterozygosity in both individuals (Fig. A2). A full-likelihood Bayesian inference suggested F1 hybrids and F1 / *M. melliger* backcrossing as the most likely hybridization scenarios for these two individuals, respectively (Fig. A3). Overall, these findings suggest ongoing hybridization between *M. mendax* cluster 4 and *M. melliger* in the Madrean Sky Islands along the U.S.-Mexico border. While syntopy has previously been reported for other montane arthropod species in the Madrean Sky Islands (e.g., *Aphonopelma*, Theraphosidae, tarantulas; Hamilton et al. 2024), this represents the first documented case of ongoing hybridization between two species of the same arthropod genus in this region.

Beyond the distinction between *M. melliger* and *M. mendax*, clusters 3 and 4, the two largest within this monophyletic group, also exhibit clear geographic and genetic separation (Fig. 1, Fig. 2, Fig. 3). Admixture signals between the clusters are subtle, yet frequent, and likely reflect a recent divergence of sister clades, potentially accompanied by incomplete lineage sorting (ILS) or ongoing gene flow. The relatively low F_ST_ value between the two clusters further indicates limited genetic differentiation (Fig. 4). Despite their distinct geographic distributions, the presence of a strong physical barrier appears unlikely given their close proximity in the northern portions of the Madrean Sky Islands (Fig. 1). Instead, ecological differentiation through niche partitioning may provide a more plausible explanation for their divergence. Cluster 3 primarily occurs within the “Madrean Pinyon–Juniper Woodland” vegetation type, whereas cluster 4 was mostly sampled in “Warm Desert Lower Montane Riparian Woodland” and “Madrean Encinal” habitats (Tab. A3). This pattern suggests a potential association with, or even preferences for different habitats along an elevational and vegetation gradient from low-elevation, desert-like areas through mid-elevation oak-dominated Encinal zones to higher-elevation pinyon–juniper woodlands (LANDFIRE 2022 EVT_CONUS, Moore et al. 2013). Notably, however, this hypothesis is challenged by the lack of distinct differences in sampling elevation (Tab. A1) and by the observation that individuals from cluster 4, though less common, also occur within “Madrean Pinyon–Juniper Woodlands” (Tab. A3). Further, more targeted sampling will be required to robustly test potential ecological differentiation between these clusters.

## Conclusion

A UCE-based phylogenomic and population genomic analysis of the genus *Myrmecocystus* from the southwestern USA and northern Mexico revealed five distinct lineages in the *M. mendax* species complex that included a larger monophyletic group sister to *M. melliger*, but also two likely cases of cryptic speciation within groups of individuals previously assigned to *M. mendax* and *M. placodops*. Beyond uncovering this cryptic species diversity, the results also indicate ongoing hybridization between the main *M. mendax* clade and *M. melliger* in the Madrean Sky Islands, as well as distinct population structure in *M. mendax* that may reflect adaptive divergence across environmental gradients. Altogether, these findings highlight the importance of incorporating molecular data to understand the evolutionary history of species complexes that exhibit frequent interbreeding and lack distinguishable morphological differentiation.

## Materials and Methods

### Sampling, DNA Extraction, Library preparation and Sequencing

Sampling was performed as outlined in van Elst et al. (2021), either by collecting from wild populations in the United States and Mexico or from various natural history collections (see Tab. A1). DNA extraction, library preparation, and target enrichment of UCEs were carried out as outlined in van Elst et al. (2021) following a modified version of the hybrid selection protocol of (Blumenstiel et al. 2010). Sequencing of the pooled libraries was conducted at the University of Utah High Throughput Genomics Core Facility (Salt Lake City, USA) on two full lanes of a 125-cycle paired-end Illumina HiSeq 2500 run.

As reported in van Elst et al. (2021), the original dataset comprised 231 *Myrmecocystus* specimens, representing 28 of the 29 described species according to Snelling’s revisions (Snelling 1976, 1982), as well as six undescribed taxa, three of which had previously been reported (Johnson and Ward 2002). In addition, 14 outgroup species from different genera were included. In this study, we used all samples for which contig assemblies had been generated previously (van Elst et al. 2021). Overall, this resulted in 214 *Myrmecocystus* samples and 12 outgroups. Tab. A1 provides sample identifiers which were reidentified in van Elst et al. (2021) using the keys from Snelling (1976, 1982), geographic coordinates and elevation data. There were three samples which were genetically placed outside their reidentified morphospecies but fell within the originally assigned morphospecies (see Figs. 2, 3). Given that these specific re-assignments were done with low confidence, the original species assignment was used instead. This includes (IDs listed as used in this study): Myrmecocystus_sp_cf_colei_RAJ_2723_MEX_BC, Myrmecocystus_sp_cf_mendax03_ KRW_no#_MEX_SO, Myrmecocystus_sp_cf_ placodops-02_KRW_no#_MEX_SO.

### Sequence data processing and UCE extraction

Illumina data were subsequently processed as outlined in van Elst et al. (2021) to generate UCE assemblies. In brief, trimmed reads were assembled using metaSPAdes v.3.13.1 (Nurk et al. 2017). UCE sequences were then identified, and paralogs removed, with the PHYLUCE pipeline (Faircloth 2016) using 9,446 probe sequences targeting 2,524 UCE loci (Branstetter et al. 2017). Probe matching resulted in a relational database containing the UCE loci recovered for each sample.

UCE statistics and UCE filter criteria were compiled based on all data reported in van Elst et al. (2021) or on a specific subset including only all 78 *M. mendax* samples. The final taxon set, used for phylogenetic analyses, was derived by including all samples of *M. mendax, M. placodops* and *M. melliger*, as well as representatives of all species that clustered with them in the maximum likelihood phylogeny constructed by van Elst et al. (2021). In addition, *M. yuma* and *M. mexicanus* were included as representatives of the subgenera *Eremnocystus* and *Myrmecocystus*, respectively, and *Lasius* samples were incorporated as outgroups. This resulted in an initial taxon set comprising a total of 142 samples, of which two were later removed for a revised phylogenetic inference (see Phylogenetic analysis with concatenated alignment). Taxon set-specific UCE were extracted using PHYLUCE v1.7.3 (Faircloth 2016) by running “phyluce_assembly_get_match_counts” (indexing UCE loci found for the specified samples), “phyluce_assembly_get_fastas_from_match_counts” (extracting UCE sequences into monolithic FASTA file), and “phyluce_assembly_explode_get_fastas_file (sorting into by-locus FASTA files)”.

### UCE statistics and filtering

To assess whether the extracted UCE sequences were sufficient to infer phylogenies at an intraspecific level, their informative site content and taxon completeness were examined with a focus on *Myrmecocystus mendax* specimens (Fig. A4, A5). To infer per-UCE informative site content, we first generated multiple sequence alignments using the FFT-NS-1 algorithm implemented in MAFFT v7.505 (Katoh and Standley 2013). Alignments were subsequently trimmed with trimAl v.1.5.1 (Capella-Gutiérrez et al. 2009) (option -automated1), and the number of sequences per locus was recorded. Locus alignments with exceptionally low taxon completeness (i.e., fewer than 30 out of 78 *M. mendax* sequences), were excluded from the calculation of informative sites. Informative sites were quantified using PhyKIT v1.11.7 (Steenwyk et al. 2021) (option -parsimony_informative_sites). A detailed taxon completeness statistic was derived from the previously generated sequence counts. Eventually, all locus alignments that showed either less than 90 % taxon completeness or contained less than three parsimony informative sites were excluded for phylogenetic inference (Fig. A4 A, B). Because mitochondrial loci can convey different phylogenetic signals due to their maternal mode of inheritance (Hurst and Jiggins 2005; Moritz et al. 1987), the corresponding UCE sequences were identified and excluded from subsequent analyses. Identification was done by querying a mitochondrial *M. mendax* genome (pers. comm. Liliya Doronina) against a local database of UCE sequences with BLAST+ v.2.15.0 (Camacho et al. 2009). Additionally, the GC content of each locus alignment was assessed with PHYKIT v1.11.7 (Steenwyk et al. 2021) (option - gc_content) (Fig. A4 C). Eventually, all samples were evaluated for locus completeness, defined as the proportion of UCE loci successfully recovered. This was determined by counting the number of locus alignments present for each sample, resulting in a final cutoff of 35% completeness (Fig. A4 D).

### Phylogenetic analysis

Filtered locus alignments were concatenated using AMAS (Borowiec 2016), which simultaneously generated a by-locus partitioning scheme. This scheme was then provided as input to PartitionFinder v2.1.1 (Lanfear et al. 2017) to estimate substitution models for each partition using the AICc model-selection criterion. PartitionFinder was set to optimize the initial partitioning scheme with the *rclusterf* algorithm (Lanfear et al. 2014; Lanfear et al. 2017) and then assign substitution models to the resulting optimized partitions.

Maximum-likelihood phylogenetic inference of the concatenated alignment was performed with IQ-TREE v2.1.4 (Minh et al. 2020), applying the optimized partitioning scheme and substitution models identified by PartitionFinder. Node support was assessed using 1,000 iterations of ultrafast bootstrapping (Hoang et al. 2018).

To minimize potential artifacts from long-branch attraction, all samples with exceptionally long branches were excluded from the final tree, including: Lasius_fuliginosus_MLB_2009, Myrmecocystus_kathjuli_JPDKED_no#_USA_CA, Myrmecocystus_sp._cf._mexicanus-01_ RRS_69-298_USA_AZ, Myrmecocystus_wheeleri_RRS_no#_USA_CA, Myrmecocystus_ sp._cf._mendax-01_FMH_no#_USA_TX, Myrmecocystus_sp._cf._placodops-02_GCS_no# _USA_AZ, Myrmecocystus_semirufus_RAJ_BC1310_MEX_BC, Myrmecocystus_semirufus _CG_no#_USA_CA, Myrmecocystus_koso_RRS_67-274_USA_CA, Myrmecocystus_sp._cf. _melliger_WPM_5148_MEX_CH.

IQ-TREE was further used to construct maximum-likelihood trees for each UCE alignment independently, again with 1,000 UFB replicates. ASTRAL-III v5.7.3 (Zhang et al. 2018) was then applied to assign quartet scores to the final maximum-likelihood tree using the generated UCE trees as input and the “-t 8” argument. The resulting tree was visualized with iTOL v7 (Letunic and Bork 2024).

### Phylogeography

Specimen coordinates were recorded during initial sampling or retrieved from museum specimens collected by van Elst et al. (2021). The digital elevation model (DEM) “SRTM90m” (NASA Shuttle Radar Topographic Mission, 90 m resolution) was downloaded for the southwestern United States and Mexico using the OpenTopography DEM Downloader v3 add-on in QGIS v3.34 (Dawson et al. 2026). Existing vegetation type (EVT) data were obtained for the continental United States from the LANDFIRE program (v2022). Maps were constructed in QGIS (Fig. 1, Fig. A1).

### Read trimming and variant calling

Raw sequencing reads generated by van Elst et al. (2021) were retrieved from the NCBI Sequence Read Archive (BioProject ID PRJNA657464), with the exception of *Lasius sitiens*, for which no FASTQ files were available. Reads were trimmed for quality with FASTP v.1.0.1 (Chen et al. 2018) using the parameter “-g” (forced polyG tail trimming), “-3” (starting sliding windows from 3’), “-l 40” (Minimum length of reads after trimming), “-Y 30” (low sequence complexity filter), “-q 15” (Minimum base phred score), “-u 40” (percentage of unqualified bases), “-c” (correction in overlapped regions). Trimmed reads were then mapped against a FASTA file containing all UCE sequences from the sample Myrmecocystus*_*sp*_*cf*_*mexicanus-02_RAJ_3061_MEX_BC, which was selected from the *M. mexicanus* specimens to serve as a suitable but not overly distant outgroup. The individual was specifically chosen because it showed high locus completeness values (95.42%). Mapping was performed with BWA-MEM v2.2.1 (Vasimuddin et al. 2019). Potential duplicate reads were subsequently removed using the Picard toolkit v2.27.5 (https://github.com/broadinstitute/picard). Mapping quality was assessed with Qualimap v2.2.2a (Okonechnikov et al. 2016). Genotype calling was performed collectively for all mapping files using BCFtools v.1.15.1 (Danecek et al. 2021) functions “mpileup” and “call”. Minimum thresholds for mapping quality (≥ 20) and base quality (≥ 13) were specified. The resulting raw VCF file was subsequently filtered as follows:

Insertions and deletions (InDels) were removed using the “view” function in BCFtools. Sites were filtered for a minimum total read depth (the sum of read depths across all samples) of 0.3 times the average total read depth. A missing data cutoff of 10% was applied while all genotype calls with a read depth below 3 were masked as missing data prior to filtering using BCFtools v1.15.1 “filter”. The 10% threshold was determined by extracting per-site missing data with VCFtools v0.1.16 (Danecek et al. 2011) (option --missing-site) and inspecting the resulting distribution (Fig. A5). Biallelic sites were extracted from the filtered VCF file with BCFtools “view”. Finally, a minimum minor allele frequency (MAF) cutoff was applied in VCFtools to exclude technical artifacts that might be misinterpreted as rare alleles. Since minor alleles were expected to occur at least twice among a total of 278 alleles, the minimum MAF was set to 2/278 ∼ 0.007.

### Population genetic analysis

Population structure was inferred using the SambaR R package v.1.10 (Jong et al. 2021), which first conducts an additional filtering step to retain only SNPs without missing data. This ensured equal levels of missing data across individuals (by setting them to zero) and thereby prevented potential distortions of ordination methods such as PCoA (Jong et al. 2021). The subsequently employed findstructure() wrapper function performs several analyses, including admixture analysis with the LEA R package v.3.16.0 (Gain and François 2021), Principal Coordinates Analysis (PCoA) with the APE package v.5.8 (Paradis and Schliep 2019), estimation of Nei’s genetic distance (Nei 1972), as well as Euclidean distance-based phylogenetic inference with the POPPR v.2.9.8 package (Kamvar et al. 2014).

The first findstructure() run included all 139 samples from the initial taxon set selected for population genetics. Input populations were defined either by strict affiliation with a morphospecies (e.g., the *M. melliger* clade) or by the phylogenetic clusters (1–5) identified in the maximum-likelihood phylogenetic analysis (Fig. 2). Subsets for subsequent runs were generated with the excludepop() function of SambaR to investigate different levels of genetic structure. For the full dataset, the LEAce() function was used to conduct a total of 18 admixture analyses with increasing numbers of presupposed lineages (K=1 to 18) to determine the optimal number of genetic clusters based on the cross-entropy criterion.

Population differentiation was assessed using pairwise F_ST_ index calculations. F_ST_ values were estimated following (Weir and Cockerham 1984) with VCFtools v0.1.16 (Danecek et al. 2011) using the “--weir-fst-pop” function. In addition to the previously defined clusters, random subsets of the phylogenetic cluster 3 were included into these cross-comparisons, which served as control F_ST_ values for total panmixia. To evaluate the reliability of these estimates, a permutation test was performed by randomly shuffeling, and reassigning individuals of each population pair into two new populations of the same sizes. After 1,000 F_ST_ values had been calculated from these permutations, a Wilcoxon signed-rank test was carried out in R (stats package v3.6.2) using the “wilcox.test” function to determine whether the observed F_ST_ value fell within the permutation distribution. Finally, a heatmap was generated in R with the ggplot2 package v4.0.0.

Genetic diversity was evaluated by counting heterozygous sites in the VCF file still containing monomorphic sites and comparing them to the total number, using the python script countHE.py of Wolf et al. (2025).

To test whether two *M. melliger* individuals (“M_melliger_JT_Hop-11_USA_AZ” and “M_ melliger*_*JT_Bue-01_MEX_SO”) represent recent hybrids between *M. mendax* cluster 4 and *M. melliger*, as suggested by the admixture analysis and PCA (Fig. 3), we applied a full-likelihood Bayesian hybrid inference approach implemented in Mongrail v1.0.2 (Chakraborty and Rannala 2023). Mongrail uses linkage and recombination information derived from high-quality, phased SNP data to estimate posterior probabilities for different hybridization scenarios, including parental lineages (P1, P2), F_1_ and F_2_ hybrids, and backcrosses to either parental population. Starting from the filtered SNP dataset used in the population genetic analyses described above, we further removed all sites containing missing genotypes using VCFtools (option --missing-site). The resulting VCF file was then split into three corresponding to individuals from the *M. melliger* population, *M. mendax* cluster 4, and the putative hybrid individuals, using BCFtools “view”. The two VCF files representing the parental populations were phased with the linkage disequilibrium-based approach in Beagle v5.5 (Browning et al. 2021). Phasing was not applied to the hybrid individuals, following the example test files from the Mongrail github repository (https://github.com/mongrail/mongrail).

Finally, scaffolds without variant information were removed from all three datasets using BCFtools “view”, and the resulting files were provided as input for Mongrail.

## Acknowledgments

We are grateful to Jay E. Taylor (Arizona State University), Roy R. Snelling^†^ (Los Angeles County Museum), Weiping Xie (Natural History Museum of Los Angeless County), William Mackay (University of Texas El Paso Biodiversity Collections), Breda Zimkus, Stefan Cover (Museum of Comparative Zoology), Alex Wild (University of Texas Biodiversity Collections: Entomology) and Steven Messer (Arizona State University) for providing samples.

## Data Accessibility

The sequences used in this study were generated in van Elst et al. (2021) and are available at the NCBI Sequence Read Archive (BioProject ID PRJNA657464). Information about the used specimens can be found in Tab. A4, A5 including determination history, voucher institution and accession numbers. A documentation of the bash and R code used in this study is stored at the GitLab repository: https://zivgitlab.uni-muenster.de/nrensing/Myrmecocystus_phylo. Files generated in this study, such as alignments or SNP data, are uploaded to a DRYAD repository: DOI: 10.5061/dryad.59zw3r2q5.

## Author contributions

J.G. and M.W. conceived and designed the study; M.W. and N.R. performed the analyses; H.N. conducted the quartet score analysis, T.v.E., T.H.E, M.B., P.W. and R.J. advised in the interpretation; all authors wrote the manuscript.

