## Supplementary Materials for "Understanding cryptic diversity within the honeypot ant species complex of *Myrmecocystus mendax*"

**This PDF file includes:**

Fig. A1 to A6

- **Fig. A1 Vegetation types across the distribution of the *Myrmecocystus mendax* species complex.**
- **Fig. A2 UCE-wide heterozygosity calculated for all phylogenetic clusters, including potentially hybrid outliers of cluster 1 and *M. melliger*.**
- **Fig. A3 Posterior probabilities retrieved from a Mongrail hybrid class test using 555 high quality SNPs derived from two potential hybrid samples.**
- **Fig. A4 Statistics on extracted UCE sequences for different sample sets and parameters.**
- **Fig. A5 Missing data per site and minor allele frequency for variant calls generated in this study.**
- **Fig. A6 Cross-entropy analysis of the optimal number of ancestral populations (K) to be defined in admixture analysis.**

Tab. A1 to A5

- **Tab. A1 Summary information for all samples used in this study.**
- **Tab. A2 Reference numbers for each sample used in the admixture analysis.**
- **Tab. A3 Vegetation types and their respective frequency associated with samples within the *M. mendax* cluster 3 to 5 defined in the phylogenomic and population genomic analyses.**
- **Tab. A4 Sampling and determination history of the samples used in this study.**
- **Tab. A5 Sample and data accession information.**

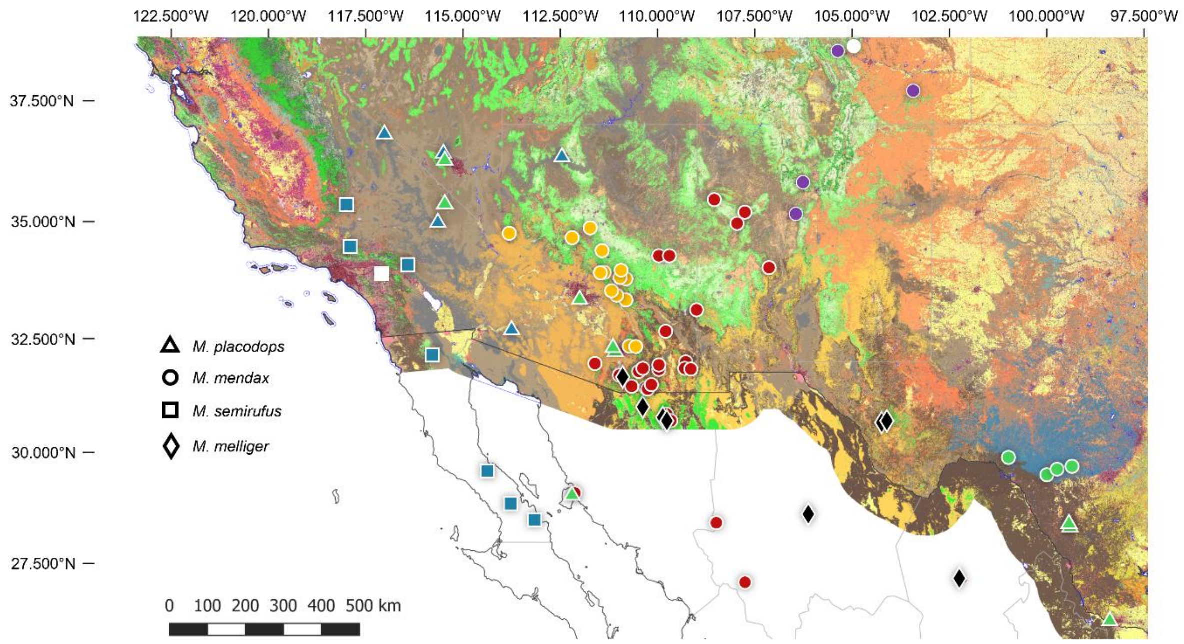


**Fig. A1 Vegetation types across the distribution of the *Myrmecocystus mendax* species complex.** Coloration of samples depicts clustering according to both the phylogenetic and population genetic analyses. Tag shape represents prior species assignments. White tags represent locations of type-specimens for the respective species given in Creighton (1950). Map constructed in QGIS v.3.34 using the LANDFIRE (2022, EVT_CONUS) vegetation type data. Vegetation data for Mexico are not represented in the LANDFIRE database. Detailed vegetation types can be retrieved from Tab. A1.

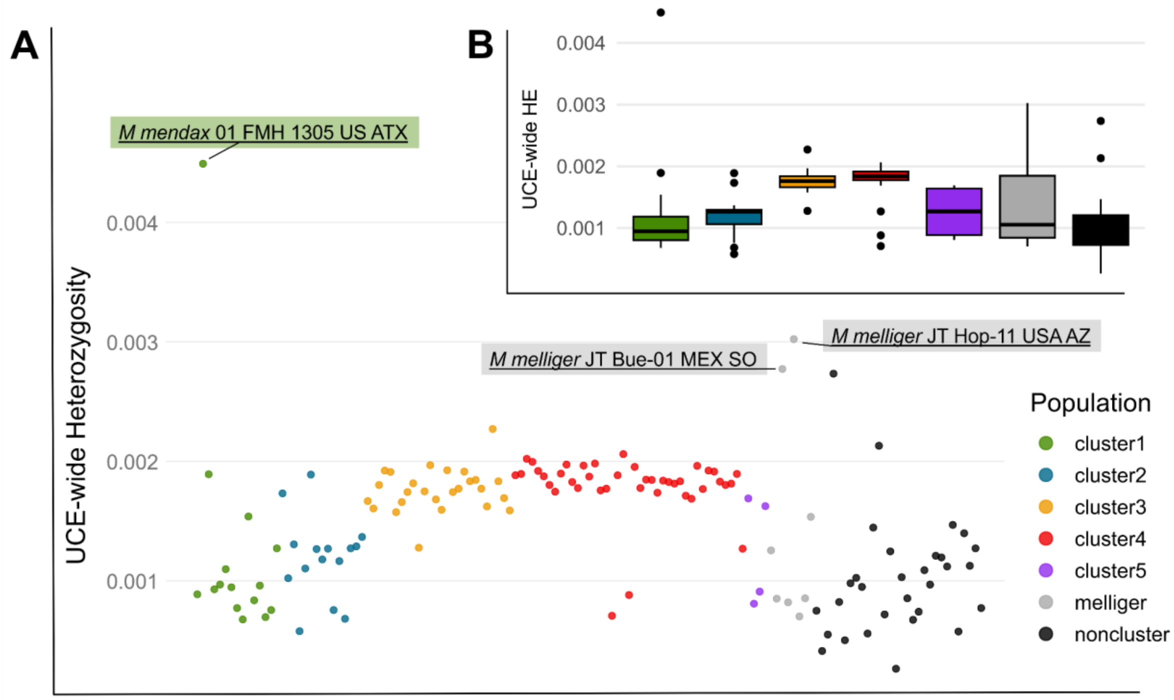


**Fig. A2 UCE-wide heterozygosity calculated for all phylogenetic clusters, including potential hybrid outliers of cluster 1 and *M. melliger*. A** Dotplot, **B** Boxplots depicting all measured values. Values have been calculated using an inhouse script (countHE.py, Wolf et al. 2025) by calculating the rate of heterozygous sites in the total VCF file containing monomorphic sites.

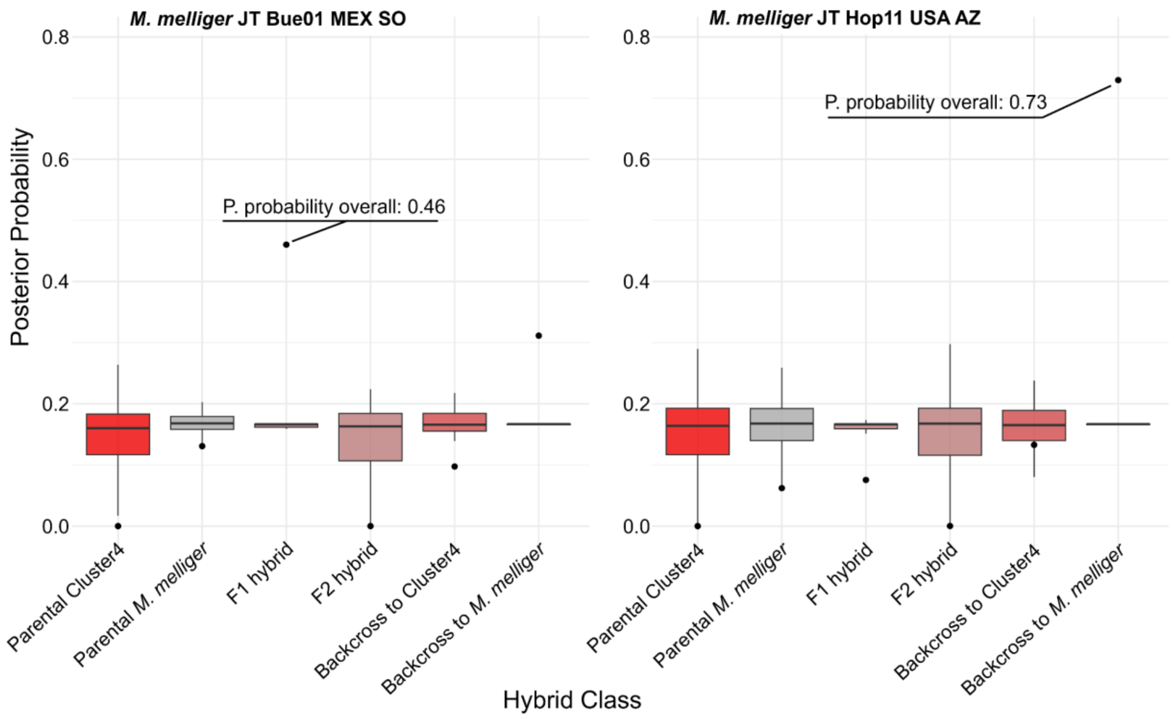


**Fig. A3 Posterior probabilities retrieved from a Mongrail hybrid class test using 555 high quality SNPs derived from two potential hybrid samples.** Boxplots depict probability distributions over all 32 included UCE. Overall probabilities, obtained by adding up the likelihood values per class for every UCE, are highlighted as dots. The highest overall posterior probability is marked additionally, indicating an F1 hybrid as the best class for JT Bue01 MEX SO and a backcross between an F1 hybrid and the *M. melliger* population as best class for JT Hop11 USA AZ.

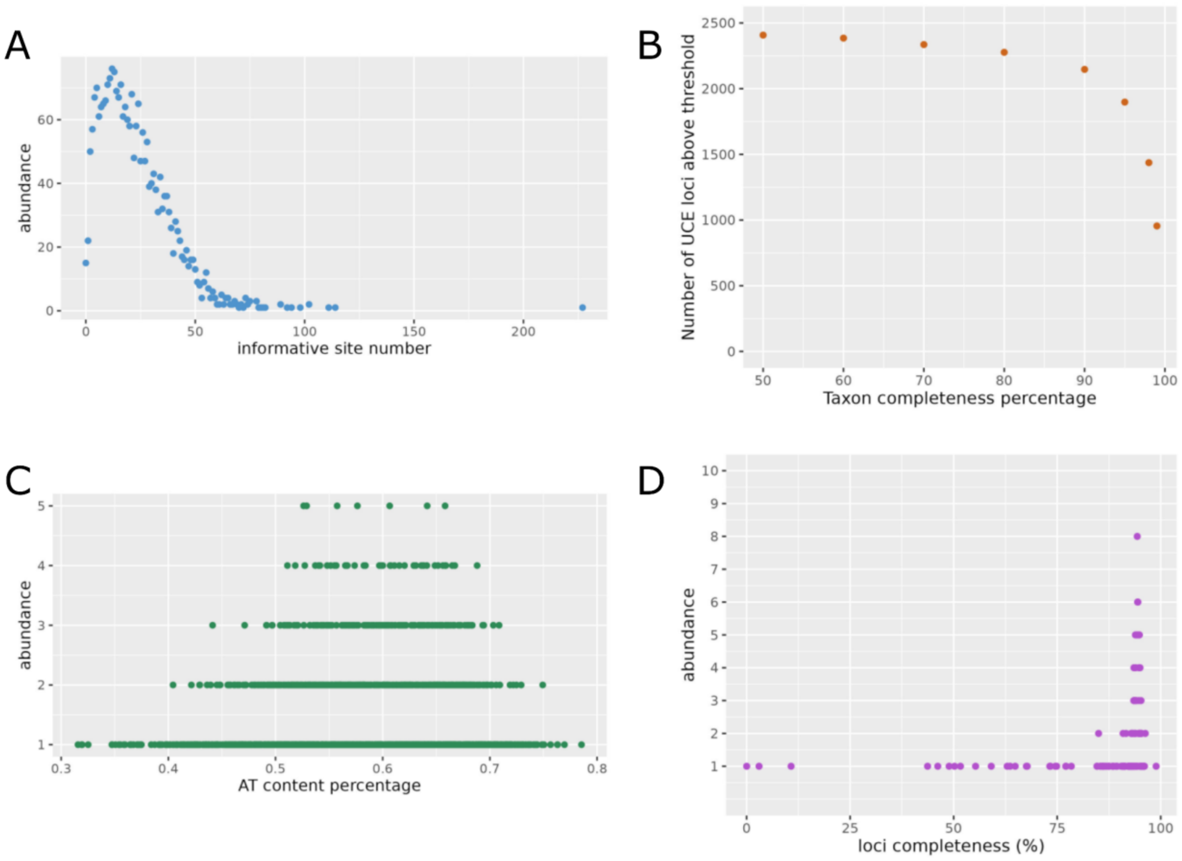


**Fig. A4 Statistics on extracted UCE sequences for different sample sets and parameters. A** Distribution of informative site content for UCE locus alignments of exclusively *Myrmecocystus mendax* samples. Only alignments with at least 30 sequences were included in the statistic. 54,796 sites of the total 2,013,645 sites (~2.7 %) were informative, and the mean informative site content was 22.43. **B** Distribution of taxon completeness for UCE locus alignments of exclusively *M. mendax* samples. Most UCE loci showed above 90% taxon completeness. **C** Distribution of AT-content for UCE loci of all samples. There were no distinct peaks in the distribution that may have allowed to distinguish mitochondrial and nuclear sequences. **D** Distribution of UCE loci completeness over all samples. In most samples, more than 75% of UCE loci were present, and the mean loci completeness was 89.77 %. There were three clear outlier samples with very low loci completeness, confirming sequencing failures as already assessed by van Elst et al. (2021).

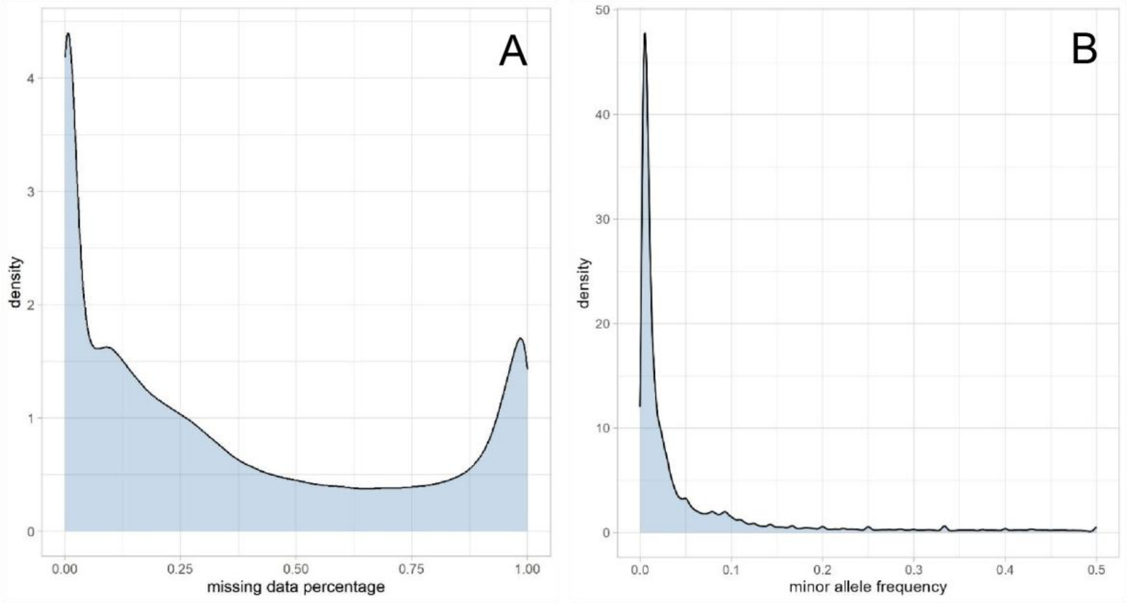


**Fig. A5 Missing data per site and minor allele frequency for variant calls generated in this study. A** Variant calls based on a read depth of three were masked as missing data. The resulting distribution showed two distinct peaks: Most sites displayed almost no missing data, and a considerable number of sites approached a missing data percentage of 100%. **B** Calculating minor allele frequencies for the 33,870 biallelic SNPs showed a high abundance of rare alleles.

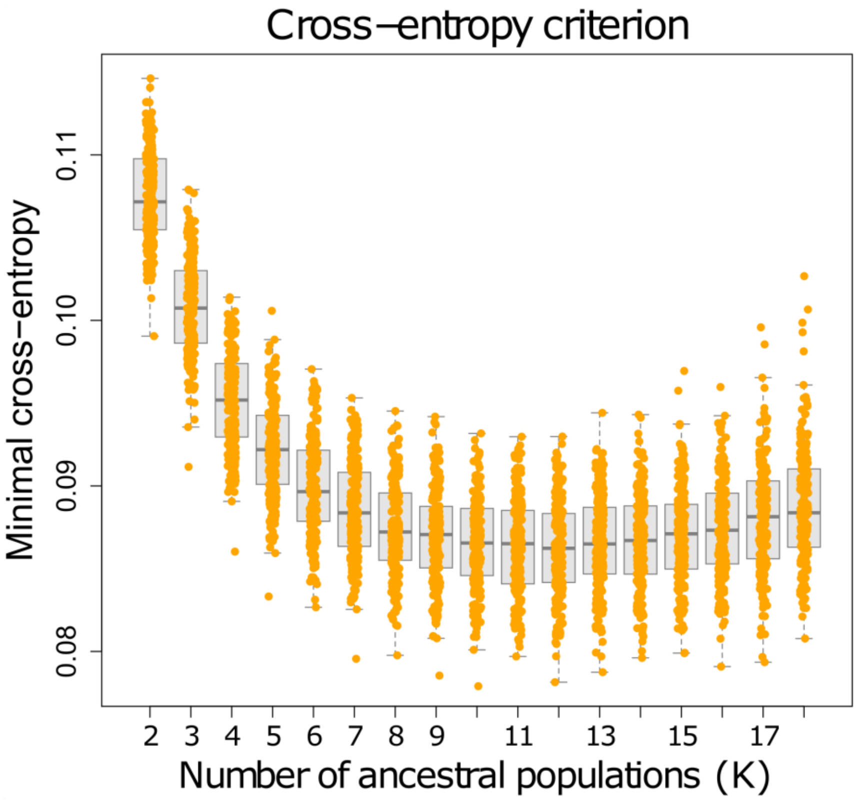


**Fig. A6 Cross-entropy analysis of the optimal number of ancestral populations (K) to be defined in admixture analysis.** In each of the 200 simulated runs of admixture analysis, the minimum cross-entropy was measured for values of K ranging from 2 to 18. Each value is represented by a data point in the graph, and the dark grey lines in the boxplots represent averages. Boxplots depict everything between the 25th and 75th percentile. Lower cross-entropy indicates a closer fit of the model to the input genetic data. The lowest average minimum cross-entropy was determined at K=12. However, no distinct and biologically relevant genetic structure was introduced after K=9, and since the minimum cross-entropy dropped only marginally above this value, it was deemed the most parsimonious assumption.

**Tab. A1 Summary information for all samples used in this study.** Specimen IDs correspond to van Elst et al. (2021). Vegetation types retrieved from LANDFIRE (2022, EVT_CONUS)**.** Colors indicate cluster-assignment based on the phylogenomic and population genetic analyses.

| SpecimenID | Latitude | Longitude | Elevation(m) | Vegetation Type |
|---|---|---|---|---|
| Formica_moki_MLB_1000_USA_CA | 38.9 | -121.02 | 450 | Mediterranean California Lower Montane Conifer Forest and Woodland |
| Gnamptogenys_simulans_Wa-D-04-2-38_NI | 13.1049 | -85.8673 | | |
| Lasius_alienus_RAJ_5575_MEX_SO | 30.9933 | -110.385 | 2260 | Madrean Lower Montane Pine-Oak Forest and Woodland |
| Lasius_arizonicus_RAJ_5576_MEX_SO | 30.9933 | -110.385 | 2260 | Madrean Lower Montane Pine-Oak Forest and Woodland |
| Lasius_colei_JT_Lc-01_USA_AZ | 32.655 | -109.805 | 1795 | Madrean Encinal |
| Lasius_fuliginosus_MLB_2009 | | | | |
| Lasius_sitiens_JTL_8439_USA_UT | 40.7898 | -111.8477 | | Inter-Mountain Basins Big Sagebrush Shrubland |
| Lasius_umbratus_MLB_145_PO_LS | 50.4342 | 19.7239 | 115 | |
| Myrmecocystus_arenarius_PSW_14630_USA_NV | 39.6086 | -119.3229 | 1271 | Great Basin & Intermountain Ruderal Shrubland |
| Myrmecocystus_arenarius_RCBRLB_no#_USA_NV | 39.1554 | -118.6252 | 1402 | Inter-Mountain Basins Cliff and Canyon |
| Myrmecocystus_christineae_RAJ_6029_USA_CA | 35.3383 | -115.56 | 1460 | Mojave Mid-Elevation Mixed Desert Scrub |
| Myrmecocystus_christineae_RAJ_6035_USA_NV | 35.4667 | -115.1883 | 1260 | Mojave Mid-Elevation Mixed Desert Scrub |
| Myrmecocystus_christineae_RRS_77-10,1_USA_CA | 35.371 | -115.473 | 1280-1298 | Mojave Mid-Elevation Mixed Desert Scrub |
| Myrmecocystus_colei_RRS_69-118_USA_CA | 34.2653 | -117.4288 | 1219 | Southern California Oak Woodland and Savanna |
| Myrmecocystus_colei_TVE_52_USA_CA | 34.2109 | -117.4007 | 605 | Southern California Coast Ranges Cliff and Canyon |
| Myrmecocystus_sp_cf_colei_RAJ_2723_MEX_BC | 28.4983 | -113.165 | 80 | |
| Myrmecocystus_creightoni_GCWJNW_NEV862_USA_NV | 36.2764 | -115.5042 | 1585 | Mojave Mid-Elevation Mixed Desert Scrub |
| Myrmecocystus_creightoni_PSW_17805_USA_CA | 34.9945 | -119.5522 | 965 | Developed-Roads |
| Myrmecocystus_creightoni_RAJ_961_USA_CA | 33.2133 | -116.4633 | 990 | Sonora-Mojave Semi-Desert Chaparral |
| Myrmecocystus_depilis_CAK_1_USA_NM | 32.3291 | -106.7464 | | Developed-Roads |
| Myrmecocystus_depilis_PSW_15480_USA_NM | 31.9433 | -109.1367 | 1391 | Chihuahuan Mixed Desert and Thornscrub |
| Myrmecocystus_depilis_SPC_6242_USA_NM | 32.7063 | -109.742 | 1080 | North American Warm Desert Ruderal & Planted Scrub |
| Myrmecocystus_ewarti_RRS_no#_USA_CA | 33.619 | -115.9161 | | Developed-Roads |
| Myrmecocystus_flaviceps_PSW_15180_USA_CA | 35.3441 | -115.4277 | 1002 | Mojave Mid-Elevation Mixed Desert Scrub |
| Myrmecocystus_flaviceps_PSW_15424_MEX_BC | 29.7317 | -114.5474 | 500 | |
| Myrmecocystus_flaviceps_RAJ_2727_MEX_BC | 31.2967 | -115.3983 | 580 | |
| Myrmecocystus_flaviceps_RAJ_2958_USA_AZ | 32.6333 | -113.7417 | 140 | North American Warm Desert Active and Stabilized Dune |
| Myrmecocystus_flaviceps_RAJ_2983_MEX_BC | 29.7283 | -114.4067 | 110 | |
| Myrmecocystus_flaviceps_RRS_94-116_USA_CA | 33.995 | -114.8883 | | North American Warm Desert Pavement |
| Myrmecocystus_flaviceps_TVE_49_USA_CA | 33.8822 | -116.3039 | 325 | North American Warm Desert Pavement |
| Myrmecocystus_hammettensis_GCWJNW_NEV1338_USA_NV | 39.4401 | -114.8307 | 1829 | Western Cool Temperate Pasture and Hayland |
| Myrmecocystus_intonsus_RRS_69-71_MEX_BCS | 23.9713 | -110.2851 | 30 | |
| Myrmecocystus_kathjuli_JPDKED_no#_USA_CA | 34.6114 | -117.92 | 844 | Sonora-Mojave Creosotebush-White Bursage Desert Scrub |
| Myrmecocystus_kathjuli_PSW_14795_USA_CA | 35.3636 | -117.9728 | 738 | Sonora-Mojave Creosotebush-White Bursage Desert Scrub |
| Myrmecocystus_kathjuli_RAJ_6007_USA_CA | 34.535 | -117.825 | 900 | Developed-Roads |
| Myrmecocystus_kathjuli_RAJ_6022_USA_CA | 34.5117 | -117.8417 | 940 | Mojave Mid-Elevation Mixed Desert Scrub |
| Myrmecocystus_kennedyi_GCSJH_98-065_USA_CA | 34.5333 | -116.8 | | Sonora-Mojave Creosotebush-White Bursage Desert Scrub |
| Myrmecocystus_kennedyi_RAJ_1249_MEX_BC | 29.6033 | -114.3917 | 300 | |
| Myrmecocystus_kennedyi_RAJ_2234_USA_AZ | 32.6967 | -113.7883 | 140 | North American Warm Desert Pavement |
| Myrmecocystus_kennedyi_RAJ_2725_MEX_BC | 28.4983 | -113.165 | 80 | |
| Myrmecocystus_kennedyi_RAJ_5778_MEX_SO | 31.2917 | -113.485 | 20 | Sonora-Mojave Creosotebush-White Bursage Desert Scrub |
| Myrmecocystus_kennedyi_RAJ_910-1_USA_AZ | 34.075 | -114.265 | 150 | Sonora-Mojave Creosotebush-White Bursage Desert Scrub |
| Myrmecocystus_koso_RRS_67-274_USA_CA | 36.1193 | -117.0951 | 2012 | Inter-Mountain Basins Big Sagebrush Shrubland |
| Myrmecocystus_placodops_SS_no#_USA_CA | 36.7882 | -117.0287 | | Sonora-Mojave Creosotebush-White Bursage Desert Scrub |
| Myrmecocystus_lugubris_GHA_no#_USA_CA | 37.1833 | -117.71 | | Sonora-Mojave Mixed Salt Desert Scrub |
| Myrmecocystus_melanoticus_LBC_no#_MEX_PU | 19.4303 | -98.4854 | 2621 | |
| Myrmecocystus_navajo_GCWJNW_NEV892_USA_NV | 37.6838 | -114.4858 | 1463 | Inter-Mountain Basins Cliff and Canyon |
| Myrmecocystus_navajo_RAJ_2914_USA_AZ | 31.6817 | -109.32 | 1620 | Apacherian-Chihuahuan Semi-Desert Shrub-Steppe |
| Myrmecocystus_navajo_RAJ_2986_MEX_BC | 29.3867 | -114.335 | 620 | |
| Myrmecocystus_navajo_RAJ_3101_MEX_BC | 31.5017 | -115.575 | 970 | |
| Myrmecocystus_navajo_RAJ_3139-2_USA_AZ | 31.93 | -109.135 | 1410 | Chihuahuan Mixed Desert and Thornscrub |
| Myrmecocystus_navajo_SPC_7476_USA_AZ | 34.9505 | -112.4227 | 1463 | Developed-Roads |
| Myrmecocystus_nequazcatl_EMF_no#_MEX_SO | 29.5031 | -112.3793 | | |
| Myrmecocystus_nequazcatl_PSW_13493_MEX_SO | 28.9399 | -112.2188 | 0 | |

| | | | | |
|---|---|---|---|---|
| Myrmecocystus_nequazcatl_RAJ_962_MEX_SO | 29.3283 | -112.26 | 10 | |
| Myrmecocystus_nequazcatl_RAJ_SON95-26_MEX_SO | 29.895 | -112.6617 | 10 | |
| Myrmecocystus_perimeces_EMFb_no#_MEX_BC | 30.4738 | -115.9477 | | |
| Myrmecocystus_pyramicus_GCWJNW_NEV1304_USA_NV | 39.063 | -115.0984 | 1829 | Inter-Mountain Basins Big Sagebrush Shrubland |
| Myrmecocystus_pyramicus_PSW_16114_USA_CA | 40.3623 | -120.2216 | 1245 | Inter-Mountain Basins Greasewood Flat |
| Myrmecocystus_romainei_MAJO_71_USA_UT | 38.6397 | -110.6317 | 1591 | Inter-Mountain Basins Active and Stabilized Dune |
| Myrmecocystus_romainei_MLB_1543_USA_AZ | 32.6939 | -113.8598 | 110 | Developed-Roads |
| Myrmecocystus_romainei_PSW_15174_USA_CA | 35.2391 | -115.3007 | 1710 | Mojave Mid-Elevation Mixed Desert Scrub |
| Myrmecocystus_romainei_PSW_16759_USA_NV | 39.2897 | -119.274 | 1288 | Developed-Roads |
| Myrmecocystus_romainei_RAJ_2349_USA_AZ | 32.705 | -109.7433 | 1079 | North American Warm Desert Ruderal & Planted Scrub |
| Myrmecocystus_romainei_RAJ_6039_USA_CA | 34.7383 | -114.6033 | 270 | Sonora-Mojave Creosotebush-White Bursage Desert Scrub |
| Myrmecocystus_romainei_RRS_73-47_USA_TX | 31.653 | -106.3026 | 1097 | Developed-Roads |
| Myrmecocystus_romainei_SPC_6239_USA_AZ | 32.7063 | -109.742 | 1200 | North American Warm Desert Ruderal & Planted Scrub |
| Myrmecocystus_semirufus_CG_no#_USA_CA | 34.4807 | -117.9106 | 1067 | Western Warm Temperate Urban Shrubland |
| Myrmecocystus_semirufus_PSW_14281_MEX_BC | 32.144 | -115.8045 | 600 | Sonora-Mojave Creosotebush-White Bursage Desert Scrub |
| Myrmecocystus_semirufus_PSW_14810_USA_CA | 35.3747 | -117.9958 | 806 | Sonora-Mojave Creosotebush-White Bursage Desert Scrub |
| Myrmecocystus_semirufus_RAJ_2984_MEX_BC | 29.5933 | -114.3917 | 320 | |
| Myrmecocystus_semirufus_RAJ_BC1310_MEX_BC | 28.8583 | -113.7917 | 600 | |
| Myrmecocystus_semirufus_TEBS_Ca-01_USA_CA | 34.0899 | -116.426 | 1190 | Developed-Roads |
| Myrmecocystus_semirufus_TEBS_Ca-02_USA_CA | 34.0905 | -116.4224 | 1190 | Mojave Mid-Elevation Mixed Desert Scrub |
| Myrmecocystus_sp_cf_colei_RAJ_2655_MEX_BC | 30.4867 | -116.0467 | 10 | |
| Myrmecocystus_sp_cf_kennedyi_RAJ_2810_USA_AZ | 33.55 | -111.5667 | 430 | Sonoran Paloverde-Mixed Cacti Desert Scrub |
| Myrmecocystus_sp_cf_kennedyi-romainei_MAJO_15_USA_NV | 38.425 | -118.1306 | 1378 | Developed-Roads |
| Myrmecocystus_sp_cf_kennedyi-romainei_PSW_15850_USA_CA | 36.2485 | -117.0778 | 2113 | Inter-Mountain Basins Big Sagebrush Shrubland |
| Myrmecocystus_sp_cf_kennedyi-romainei_RAJ_6009_USA_CA | 34.4733 | -117.9083 | 1100 | Developed-Roads |
| Myrmecocystus_sp_cf_melliger_DMO_no#_MEX_CO | 27.1667 | -102.25 | 900 | |
| Myrmecocystus_sp_cf_melliger_JCA_2121_USA_TX | 30.699 | -104.117 | | Western Warm Temperate Urban Herbaceous |
| Myrmecocystus_sp_cf_melliger_JT_Bue-01_MEX_SO | 30.7728 | -109.8665 | 1670 | Madrean Lower Montane Pine-Oak Forest and Woodland |
| Myrmecocystus_sp_cf_melliger_JT_Bue-09_MEX_SO | 30.7334 | -109.8208 | 1690 | Madrean Lower Montane Pine-Oak Forest and Woodland |
| Myrmecocystus_sp_cf_melliger_JT_Hop-11_USA_AZ | 31.6725 | -110.9053 | 1750 | Madrean Encinal |
| Myrmecocystus_sp_cf_melliger_RAJ_5588_MEX_SO | 31.0117 | -110.39 | 1940 | Madrean Lower Montane Pine-Oak Forest and Woodland |
| Myrmecocystus_sp_cf_melliger_RRS_no#_MEX_PU | 18.6476 | -97.3788 | 2042 | |
| Myrmecocystus_sp_cf_melliger_WPEM_17561_USA_TX | 30.6642 | -104.2464 | 1777 | Western Warm Temperate Urban Shrubland |
| Myrmecocystus_sp_cf_melliger_WPM_5148_MEX_CH | 28.6276 | -106.1368 | | |
| Myrmecocystus_sp_cf_mendax-01_AW_6127_USA_TX | 29.6974 | -99.3621 | 500 | Edwards Plateau Limestone Shrubland |
| Myrmecocystus_sp_cf_mendax-01_FMH_1305_USA_TX | 29.5 | -100 | | Edwards Plateau Limestone Shrubland |
| Myrmecocystus_sp_cf_mendax-01_FMH_no#_USA_TX | 29.5 | -100 | | Edwards Plateau Limestone Shrubland |
| Myrmecocystus_sp_cf_mendax-01_JDH_no#_USA_TX | 29.8845 | -100.9935 | | Edwards Plateau Floodplain Terrace Shrubland |
| Myrmecocystus_sp_cf_mendax-01_WPM_13384_USA_TX | 29.6282 | -99.7444 | | Edwards Plateau Riparian Herbaceous |
| Myrmecocystus_sp_cf_mendax-03_JT_Fre-02_USA_AZ | 31.8086 | -110.3889 | 1560 | North American Warm Desert Lower Montane Riparian Woodland |
| Myrmecocystus_sp_cf_mendax-03_JT_Hop-03_USA_AZ | 31.6711 | -110.9031 | 1760 | Mogollon Chaparral |
| Myrmecocystus_sp_cf_mendax-03_JT_Mol-02_USA_AZ | 32.3361 | -110.6925 | 1320 | Chihuahuan Succulent Desert Scrub |
| Myrmecocystus_sp_cf_mendax-03_JT_Mol-03_USA_AZ | 32.3367 | -110.6922 | 1320 | North American Warm Desert Lower Montane Riparian Woodland |
| Myrmecocystus_sp_cf_mendax-03_JT_Ord-03_USA_AZ | 33.9192 | -111.4475 | 1440 | Madrean Pinyon-Juniper Woodland |
| Myrmecocystus_sp_cf_mendax-03_JT_Ord-05_USA_AZ | 33.9181 | -111.4358 | 1430 | Madrean Pinyon-Juniper Woodland |
| Myrmecocystus_sp_cf_mendax-03_JT_Ord-08_USA_AZ | 33.9192 | -111.4175 | 1750 | Madrean Encinal |
| Myrmecocystus_sp_cf_mendax-03_JT_Syc-03_USA_AZ | 32.3375 | -110.7269 | 1470 | Mogollon Chaparral |
| Myrmecocystus_sp_cf_mendax-03_KRW_no#_MEX_SO | 29.0667 | -112.1333 | 50 | |
| Myrmecocystus_sp_cf_mendax-03_MH_Sup-03_USA_AZ | 33.5334 | -111.1932 | 1430 | Madrean Pinyon-Juniper Woodland |
| Myrmecocystus_sp_cf_mendax-03_PSW_14509_USA_AZ | 33.7817 | -110.9685 | 1432 | Developed-Roads |
| Myrmecocystus_sp_cf_mendax-03_TE_Don-01_USA_AZ | 34.6576 | -112.196 | 1720 | Madrean Pinyon-Juniper Woodland |
| Myrmecocystus_sp_cf_mendax-03_TE_Dri-01_USA_AZ | 34.3859 | -111.4364 | 1800 | Madrean Encinal |
| Myrmecocystus_sp_cf_mendax-03_TE_Pat-01_USA_AZ | 31.4565 | -110.6667 | 1730 | Madrean Encinal |
| Myrmecocystus_sp_cf_mendax-03_TE_Pat-08_USA_AZ | 31.4565 | -110.6661 | 1480 | Madrean Encinal |
| Myrmecocystus_sp_cf_mendax-03_TE_Pat-09_USA_AZ | 31.4566 | -110.6669 | 1480 | Madrean Encinal |
| Myrmecocystus_sp_cf_mendax-03_TE_Pcr-01_USA_AZ | 33.4336 | -111.0619 | 1090 | Madrean Pinyon-Juniper Woodland |
| Myrmecocystus_sp_cf_mendax-03_TE_Pcr-10_USA_AZ | 33.4309 | -111.0567 | 1190 | North American Warm Desert Bedrock Cliff and Outcrop |
| Myrmecocystus_sp_cf_mendax-03_TE_Pin-04_USA_AZ | 33.3394 | -110.8159 | 1290 | Southern Rocky Mountain Ponderosa Pine Woodland |
| Myrmecocystus_sp_cf_mendax-03_TE_Pin-07_USA_AZ | 33.3406 | -110.8158 | 1280 | Madrean Pinyon-Juniper Woodland |
| Myrmecocystus_sp_cf_mendax-03_TE_Pin-14_USA_AZ | 33.3406 | -110.8195 | 1310 | Madrean Pinyon-Juniper Woodland |
| Myrmecocystus_sp_cf_mendax-03_TE_Sch-01_USA_AZ | 34.8671 | -111.7461 | 1380 | Developed-Roads |

| | | | | |
|---|---|---|---|---|
| Myrmecocystus_sp_cf_mendax-03_TE_Sch-07_USA_AZ | 34.8668 | -111.741 | 1380 | Madrean Pinyon-Juniper Woodland |
| Myrmecocystus_sp_cf_mendax-03_TE_Sch-11_USA_AZ | 34.8714 | -111.7332 | 1440 | Madrean Pinyon-Juniper Woodland |
| Myrmecocystus_sp_cf_mendax-03_TE_Sie-05_USA_AZ | 33.7981 | -110.9707 | 1570 | Madrean Encinal |
| Myrmecocystus_sp_cf_mendax-03_TE_Sie-126_USA_AZ | 33.9697 | -110.9485 | 1800 | Developed-Roads |
| Myrmecocystus_sp_cf_mendax-03_TE_Sie-143_USA_AZ | 33.9724 | -110.9473 | 1760 | Developed-Roads |
| Myrmecocystus_sp_cf_mendax-03_TE_Sie-151_USA_AZ | 33.9716 | -110.9462 | 1695 | Madrean Encinal |
| Myrmecocystus_sp_cf_mendax-03_TE_Sie-20_USA_AZ | 33.7974 | -110.9831 | 1680 | Developed-Roads |
| Myrmecocystus_sp_cf_mendax-03_TE_Sie-72_USA_AZ | 33.8082 | -110.981 | 1758 | Developed-Roads |
| Myrmecocystus_sp_cf_mendax-03_TECO_Ced-01_USA_AZ | 34.7521 | -113.8049 | 1300 | Mogollon Chaparral |
| Myrmecocystus_sp_cf_mendax-03_TECO_Ced-02_USA_AZ | 34.752 | -113.8055 | 1300 | Mogollon Chaparral |
| Myrmecocystus_sp_cf_mendax-04_JGLW_Chi-02_USA_AZ | 31.8877 | -109.2084 | | North American Warm Desert Lower Montane Riparian Woodland |
| Myrmecocystus_sp_cf_mendax-04_JGLW_Chi-06_USA_AZ | 31.8877 | -109.2084 | | North American Warm Desert Lower Montane Riparian Woodland |
| Myrmecocystus_sp_cf_mendax-04_JGLW_Chi-08_USA_AZ | 31.8877 | -109.2084 | | North American Warm Desert Lower Montane Riparian Woodland |
| Myrmecocystus_sp_cf_mendax-04_JT_Bue-02_MEX_SO | 30.7564 | -109.8334 | 1520 | Madrean Encinal |
| Myrmecocystus_sp_cf_mendax-04_JT_Bue-04_MEX_SO | 30.7384 | -109.8185 | 1600 | Madrean Encinal |
| Myrmecocystus_sp_cf_mendax-04_JT_Bue-11_MEX_SO | 30.7728 | -109.8665 | 1670 | Madrean Lower Montane Pine-Oak Forest and Woodland |
| Myrmecocystus_sp_cf_mendax-04_JT_Car-01_USA_AZ | 31.4447 | -110.2578 | 1630 | North American Warm Desert Ruderal & Planted Scrub |
| Myrmecocystus_sp_cf_mendax-04_JT_Coc-02_USA_AZ | 31.9228 | -109.9669 | 1500 | North American Warm Desert Lower Montane Riparian Woodland |
| Myrmecocystus_sp_cf_mendax-04_JT_Coc-03_USA_AZ | 31.9225 | -109.9669 | 1500 | North American Warm Desert Lower Montane Riparian Woodland |
| Myrmecocystus_sp_cf_mendax-04_JT_Coc-10_USA_AZ | 31.9075 | -109.9675 | 1640 | North American Warm Desert Lower Montane Riparian Woodland |
| Myrmecocystus_sp_cf_mendax-04_JT_Fre-04_USA_AZ | 31.8092 | -110.3914 | 1560 | North American Warm Desert Lower Montane Riparian Woodland |
| Myrmecocystus_sp_cf_mendax-04_JT_Fre-05_USA_AZ | 31.815 | -110.4031 | 1560 | North American Warm Desert Lower Montane Riparian Woodland |
| Myrmecocystus_sp_cf_mendax-04_JT_Kit-01_USA_AZ | 31.9654 | -111.621 | 1600 | Madrean Encinal |
| Myrmecocystus_sp_cf_mendax-04_JT_Mil-01_USA_AZ | 31.4267 | -110.2578 | 1570 | Developed-Roads |
| Myrmecocystus_sp_cf_mendax-04_JT_Ram-01_USA_AZ | 31.4403 | -110.3156 | 1920 | Madrean Pinyon-Juniper Woodland |
| Myrmecocystus_sp_cf_mendax-04_PSW_13407-6_USA_NM | 35.2 | -106.5 | 1860 | Inter-Mountain Basins Semi-Desert Shrub-Steppe |
| Myrmecocystus_sp_cf_mendax-04_PSW_16919-01_MEX_CH | 27.0688 | -107.7599 | 1110 | |
| Myrmecocystus_sp_cf_mendax-04_RAJ_2536_MEX_CH | 31.8767 | -109.2317 | 1785 | Madrean Encinal |
| Myrmecocystus_sp_cf_mendax-04_RAJ_2800_MEX_CH | 28.4333 | -108.5 | 1611 | |
| Myrmecocystus_sp_cf_mendax-04_RAJ_4930_USA_AZ | 34.2733 | -109.6933 | 2100 | Colorado Plateau Pinyon-Juniper Woodland |
| Myrmecocystus_sp_cf_mendax-04_TE_Chi-50_USA_AZ | 31.9122 | -109.2424 | 1870 | Developed-Roads |
| Myrmecocystus_sp_cf_mendax-04_TEBS_Co-01_USA_CO | 37.7447 | -103.4845 | 1290 | Inter-Mountain Basins Mixed Salt Desert Scrub |
| Myrmecocystus_sp_cf_mendax-04_TEBS_Nm-01_USA_NM | 35.4625 | -108.5438 | 2180 | Colorado Plateau Pinyon-Juniper Woodland |
| Myrmecocystus_sp_cf_mendax-04_TEBS_Nm-02_USA_NM | 34.956 | -107.9495 | 2120 | Inter-Mountain Basins Semi-Desert Grassland |
| Myrmecocystus_sp_cf_mendax-04_TEBS_Nm-03_USA_NM | 34.9556 | -107.9496 | 2120 | Developed-Roads |
| Myrmecocystus_sp_cf_mendax-04_TEBS_Nm-04_USA_NM | 34.9575 | -107.9542 | 2130 | Inter-Mountain Basins Semi-Desert Shrub-Steppe |
| Myrmecocystus_sp_cf_mendax-04_TEBS_Nm-05_USA_NM | 34.9619 | -107.9557 | 2150 | Colorado Plateau Pinyon-Juniper Woodland |
| Myrmecocystus_sp_cf_mendax-04_TEBS_Nm-08_USA_NM | 35.1999 | -107.7557 | 2170 | Inter-Mountain Basins Mixed Salt Desert Scrub |
| Myrmecocystus_sp_cf_mendax-04_TEJGJT_Mor-01_USA_AZ | 34.2671 | -109.9721 | 1960 | Developed-Roads |
| Myrmecocystus_sp_cf_mendax-04_TEJGJT_Mor-06_USA_AZ | 34.2806 | -109.9671 | 1950 | Interior West Ruderal Riparian Forest |
| Myrmecocystus_sp_cf_mendax-04_TEJGJT_Mor-08_USA_AZ | 34.2753 | -109.693 | 2100 | Developed-Roads |
| Myrmecocystus_sp_cf_mendax-04_TEJGJT_Sky-03_USA_AZ | 32.664 | -109.797 | 1600 | Madrean Encinal |
| Myrmecocystus_sp_cf_mendax-04_TEJGJT_Sky-06_USA_AZ | 32.6627 | -109.7939 | 1580 | Madrean Pinyon-Juniper Woodland |
| Myrmecocystus_sp_cf_mendax-04_TEJGJT_Sky-13_USA_AZ | 32.6649 | -109.797 | 1600 | Mogollon Chaparral |
| Myrmecocystus_sp_cf_mendax-04_WPM_16472_USA_NM | 33.1227 | -109.0038 | 1018 | Madrean Pinyon-Juniper Woodland |
| Myrmecocystus_sp_cf_mendax-04_WPM_16702_USA_NM | 34.0234 | -107.1325 | 1930 | Madrean Pinyon-Juniper Woodland |
| Myrmecocystus_sp_cf_mendax-04_WPM_16704_USA_NM | 34.0234 | -107.1325 | 1930 | Madrean Pinyon-Juniper Woodland |
| Myrmecocystus_sp_cf_mendax-04_WPM_19359_USA_CO | 38.5501 | -105.4239 | | Inter-Mountain Basins Semi-Desert Shrub-Steppe |
| Myrmecocystus_sp_cf_mendax-04_WPM_no#_USA_NM | 35.8522 | -106.3149 | 1770 | Southern Rocky Mountain Pinyon-Juniper Woodland |
| Myrmecocystus_sp_cf_mexicanus-01_RRS_69-298_USA_AZ | 31.3514 | -110.2859 | 2096 | Developed-Roads |
| Myrmecocystus_sp_cf_mexicanus-02_MLB_1122_USA_AZ | 33.6349 | -115.3973 | 670 | Sonoran Paloverde-Mixed Cacti Desert Scrub |
| Myrmecocystus_sp_cf_mexicanus-02_MLB_421_USA_NV | 39.2898 | -119.274 | 1290 | Developed-Roads |
| Myrmecocystus_sp_cf_mexicanus-02_RAJ_3061_MEX_BC | 31.275 | -115.3217 | 520 | |
| Myrmecocystus_sp_cf_mexicanus-02_RAJ_6012_USA_CA | 34.4633 | -117.9083 | 1160 | Mojave Mid-Elevation Mixed Desert Scrub |
| Myrmecocystus_sp_cf_mexicanus-02_RAJ_6028_USA_CA | 35.42 | -115.6517 | 1170 | Developed-Roads |
| Myrmecocystus_sp_cf_mexicanus-02_SPC_6256_USA_CA | 34.2087 | -112.1417 | 1245 | Sonoran Paloverde-Mixed Cacti Desert Scrub |
| Myrmecocystus_sp_cf_mimicus-flaviceps-01_PSW_10552_MEX_BC | 26.8667 | -113.1333 | 9 | |
| Myrmecocystus_sp_cf_mimicus-flaviceps-01_RAJ_2990_MEX_BC | 29.3867 | -114.38 | 630 | |
| Myrmecocystus_sp_cf_mimicus-flaviceps-01_RAJ_3029_MEX_BC | 27.7767 | -113.5733 | 70 | |
| Myrmecocystus_sp_cf_mimicus-flaviceps-01_RAJ_3076_MEX_BC | 31.2133 | -115.6117 | 1200 | |
| Myrmecocystus_sp_cf_mimicus-flaviceps-01_RAJ_6025_USA_CA | 35.145 | -118.4483 | 1200 | Northern and Central California Dry-Mesic Chaparral |

| | | | | |
|---|---|---|---|---|
| Myrmecocystus_sp_cf_mimicus-flaviceps-01_RRS_77-22_USA_CA | 34.478 | -117.9112 | | Mojave Mid-Elevation Mixed Desert Scrub |
| Myrmecocystus_sp_cf_mimicus-flaviceps-02_RAJ_2665_MEX_BCS | 27.6733 | -113.4017 | 90 | |
| Myrmecocystus_sp_cf_mimicus-flaviceps-02_RAJ_3030_MEX_BCS | 27.7767 | -113.5733 | 70 | |
| Myrmecocystus_sp_cf_mimicus-flaviceps-03_JT_Fre-01_USA_AZ | 31.8086 | -110.3881 | 1556 | Madrean Encinal |
| Myrmecocystus_sp_cf_mimicus-flaviceps-03_MLB_1544_USA_AZ | 34.5912 | -112.4235 | 1590 | Madrean Encinal |
| Myrmecocystus_sp_cf_mimicus-flaviceps-03_RAJ_5784_USA_AZ | 32.9317 | -111.7083 | 430 | Sonora-Mojave Creosotebush-White Bursage Desert Scrub |
| Myrmecocystus_sp_cf_mimicus-flaviceps-03_SPC_4846_USA_AZ | 33.5486 | -113.1832 | 384 | Sonoran Paloverde-Mixed Cacti Desert Scrub |
| Myrmecocystus_sp_cf_navajo_RAJ_3077_MEX_BC | 31.2133 | -115.6117 | 1200 | |
| Myrmecocystus_sp_cf_placodops-01_ELVSDP_no#_USA_TX | 28.3292 | -99.4107 | | Tamaulipan Calcareous Thornscrub |
| Myrmecocystus_sp_cf_placodops-01_PSW_15596_USA_TX | 26.1789 | -98.379 | 30 | Tamaulipan Floodplain Herbaceous |
| Myrmecocystus_sp_cf_placodops-01_RWP_no#_USA_TX | 28.34 | -99.43 | | Tamaulipan Mixed Deciduous Thornscrub |
| Myrmecocystus_sp_cf_placodops-02_GCS_no#_USA_AZ | 32.2354 | -111.0851 | | Sonoran Mid-Elevation Desert Scrub |
| Myrmecocystus_sp_cf_placodops-02_KRW_no#_MEX_SO | 29.0607 | -112.1656 | 10 | |
| Myrmecocystus_sp_cf_placodops-02_PSW_16232_USA_AZ | 32.2244 | -111.1006 | 960 | Sonora-Mojave Creosotebush-White Bursage Desert Scrub |
| Myrmecocystus_sp_cf_placodops-02_RRS_77-12_USA_CA | 35.371 | -115.473 | 1280-1298 | Mojave Mid-Elevation Mixed Desert Scrub |
| Myrmecocystus_sp_cf_placodops-02_TE_Sou-01_USA_AZ | 33.3484 | -112.005 | 450 | Sonoran Paloverde-Mixed Cacti Desert Scrub |
| Myrmecocystus_sp_cf_placodops-02_TEBS_Nv-02_USA_NV | 36.2803 | -115.4367 | 1320 | Mojave Mid-Elevation Mixed Desert Scrub |
| Myrmecocystus_sp_cf_placodops-02_TEBS_Nv-03_USA_NV | 36.2821 | -115.4344 | 1320 | Mojave Mid-Elevation Mixed Desert Scrub |
| Myrmecocystus_sp_cf_placodops-03_PSW_14981-03_USA_AZ | 36.348 | -112.455 | 610 | Colorado Plateau Mixed Bedrock Canyon and Tableland |
| Myrmecocystus_sp_cf_placodops-03_PSW_15153_USA_CA | 34.9795 | -115.6476 | 239 | Sonora-Mojave Creosotebush-White Bursage Desert Scrub |
| Myrmecocystus_sp_cf_placodops-03_RAJ_2978_USA_AZ | 32.7033 | -113.745 | 180 | Sonoran Paloverde-Mixed Cacti Desert Scrub |
| Myrmecocystus_sp_cf_placodops-03_TEBS_Nv-01_USA_NV | 36.2813 | -115.435 | 1320 | Mojave Mid-Elevation Mixed Desert Scrub |
| Myrmecocystus_sp_cf_placodops-03_TEBS_Nv-04_USA_NV | 36.4476 | -115.5078 | 1250 | Mojave Mid-Elevation Mixed Desert Scrub |
| Myrmecocystus_sp_cf_placodops-03_TEBS_Nv-05_USA_NV | 36.4485 | -115.5067 | 1240 | Mojave Mid-Elevation Mixed Desert Scrub |
| Myrmecocystus_sp_SON-1_RAJ_5781_MEX_SO | 31.295 | -113.51 | 15 | Sonora-Mojave Creosotebush-White Bursage Desert Scrub |
| Myrmecocystus_tenuinodis_PSW_15178_USA_CA | 34.9795 | -115.6476 | 665 | Sonora-Mojave Creosotebush-White Bursage Desert Scrub |
| Myrmecocystus_tenuinodis_RRS_no#_USA_CA | 33.8278 | -116.4874 | 58 | Developed-Low Intensity |
| Myrmecocystus_tenuinodis_SPC_4826_USA_AZ | 34.0936 | -114.2209 | 140 | North American Warm Desert Pavement |
| Myrmecocystus_testaceus_MLB_1258_USA_CA | 39.3512 | -122.749 | 1680 | Developed-Roads |
| Myrmecocystus_testaceus_MLB_397_USA_NV | 41.6362 | -119.842 | 1695 | Inter-Mountain Basins Big Sagebrush Shrubland |
| Myrmecocystus_testaceus_MLB_446_USA_CA | 39.774 | -120.0732 | 1470 | Inter-Mountain Basins Big Sagebrush Shrubland |
| Myrmecocystus_testaceus_RAJ_2259_MEX_BC | 30.5 | -116.0033 | 10 | |
| Myrmecocystus_testaceus_RAJ_6018_USA_CA | 34.4183 | -117.79 | 1540 | Developed-Roads |
| Myrmecocystus_testaceus_RRSCDG_77-20_USA_CA | 35.3395 | -115.4795 | 1200-1300 | Mojave Mid-Elevation Mixed Desert Scrub |
| Myrmecocystus_wheeleri_MLB_1256_USA_CA | 36.539 | -120.5669 | 500 | California Ruderal Scrub |
| Myrmecocystus_wheeleri_PSW_14302_USA_CA | 33.0888 | -116.4975 | 805 | Western Warm Temperate Urban Shrubland |
| Myrmecocystus_wheeleri_RAJ_3094_MEX_BC | 31.5917 | -115.87 | 1030 | |
| Myrmecocystus_wheeleri_RAJ_6010_USA_CA | 34.4733 | -117.9083 | 1100 | Developed-Roads |
| Myrmecocystus_wheeleri_RAJ_6013_USA_CA | 34.4633 | -117.9083 | 1160 | Mojave Mid-Elevation Mixed Desert Scrub |
| Myrmecocystus_wheeleri_RAJ_6021_USA_CA | 34.5 | -118.2283 | 930 | Western Warm Temperate Urban Herbaceous |
| Myrmecocystus_wheeleri_RRS_no#_USA_CA | 34.4386 | -117.8395 | 1067 | Mojave Mid-Elevation Mixed Desert Scrub |
| Myrmecocystus_yuma_PSW_15185_USA_CA | 35.3441 | -115.4277 | 1000 | Mojave Mid-Elevation Mixed Desert Scrub |
| Myrmecocystus_yuma_RAJ_2243_USA_AZ | 32.6333 | -113.7417 | 150 | North American Warm Desert Active and Stabilized Dune |
| Myrmecocystus_yuma_RAJ_3084_MEX_BC | 31.32 | -115.5133 | 930 | |
| Myrmecocystus_yuma_RAJ_5783_MEX_SO | 31.3 | -113.52 | 20 | Sonora-Mojave Creosotebush-White Bursage Desert Scrub |
| Myrmecocystus_yuma_RRS_94-11a_USA_CA | 33.995 | -114.8883 | | North American Warm Desert Pavement |
| Myrmelachista_joycei_JTL_8492_CR | 10.3134 | -84.7121 | | |
| Nylanderia_terricola_JTL_14763_USA_TX | 30.09 | -99.51 | 610 | Edwards Plateau Limestone Shrubland |
| Paratrechina_longicornis_LRD_270806-14_USA_FL | 24.6515 | -81.3058 | | Eastern Warm Temperate Urban Herbaceous |
| Polyergus_mexicanus_MLB_389-01_USA_CA | 39.4159 | -120.317 | 2630 | Mediterranean California Subalpine Woodland |

**Tab. A2 Reference numbers for each sample used in the admixture analysis.** The numbers in the “Ref.” columns refer to the sample numbers shown below each column of the admixture plot (Fig. 3).

| Ref. | Sample ID | Ref. | Sample ID |
|---|---|---|---|
| 10 | M_placodopsRRS67274USACASRR12600378 | 66 | M_mendax-04JGLWChi06USAAZSRR12600328 |
| 11 | M_semirufusCGnoUSACASRR12600483 | 67 | M_mendax-04JGLWChi08USAAZSRR12600327 |
| 12 | M_semirufusPSW14281MEXBCSRR12600482 | 68 | M_mendax-04JTBue02MEXSOSRR12600326 |
| 13 | M_semirufusPSW14810USACASRR12600481 | 69 | M_mendax-04JTBue04MEXSOSRR12600325 |
| 14 | M_semirufusRAJ2984MEXBCSRR12600480 | 70 | M_mendax-04JTBue11MEXSOSRR12600324 |
| 15 | M_semirufusRAJBC1310MEXBCSRR12600479 | 71 | M_mendax-04JTCar01USAAZSRR12600323 |
| 16 | M_semirufusTEBSCa01USACASRR12600478 | 72 | M_mendax-04JTCoc02USAAZSRR12600322 |
| 17 | M_semirufusTEBSCa02USACASRR12600477 | 73 | M_mendax-04JTCoc03USAAZSRR12600321 |
| 19 | M_melligerDMOnoMEXCOSRR12600470 | 74 | M_mendax-04JTCoc10USAAZSRR12600319 |
| 20 | M_melligerJCA2121USATXSRR12600469 | 75 | M_mendax-04JTFre04USAAZSRR12600318 |
| 21 | M_melligerJTBue01MEXSOSRR12600468 | 76 | M_mendax-04JTFre05USAAZSRR12600317 |
| 22 | M_melligerJTBue09MEXSOSRR12600467 | 77 | M_mendax-04_JTKit01USAAZSRR12600316 |
| 23 | M_melliger_JTHop11USAAZSRR12600466 | 78 | M_mendax-04_JTMil01USAAZSRR12600315 |
| 24 | M_melliger_RAJ5588MEXSOSRR12600465 | 79 | M_mendax-04_JTRam01USAAZSRR12600314 |
| 25 | M_melliger_WPEM17561USATXSRR12600462 | 80 | M_mendax-04_PSW134076USANMSRR12600313 |
| 26 | M_melliger_WPM5148MEXCHSRR12600461 | 81 | M_mendax-04_PSW1691901MEXCHSRR12600312 |
| 27 | M_mendax-01_AW6127USATXSRR12600460 | 82 | M_mendax-04_RAJ2536MEXCHSRR12600311 |
| 28 | M_mendax-01_FMH1305USATXSRR12600459 | 83 | M_mendax-04_RAJ2800MEXCHSRR12600310 |
| 29 | M_mendax-01_FMHnoUSATXSRR12600458 | 84 | M_mendax-04_RAJ4930USAAZSRR12600308 |
| 30 | M_mendax-01_JDHnoUSATXSRR12600457 | 85 | M_mendax-04_TEBSCo01USACOSRR12600306 |
| 31 | M_mendax-01_WPM13384USATXSRR12600456 | 86 | M_mendax-04_TEBSNm01USANMSRR12600305 |
| 32 | M_mendax-02_SSnoUSACASRR12600455 | 87 | M_mendax-04_TEBSNm02USANMSRR12600304 |
| 33 | M_mendax-03_JTFre02USAAZSRR12600454 | 88 | M_mendax-04_TEBSNm03USANMSRR12600303 |
| 34 | M_mendax-03_JTHop03USAAZSRR12600452 | 89 | M_mendax-04_TEBSNm04USANMSRR12600302 |
| 35 | M_mendax-03_JTMol02USAAZSRR12600451 | 90 | M_mendax-04_TEBSNm05USANMSRR12600301 |
| 36 | M_mendax-03_JTMol03USAAZSRR12600450 | 91 | M_mendax-04_TEBSNm08USANMSRR12600300 |
| 37 | M_mendax-03_JTOrd03USAAZSRR12600449 | 92 | M_mendax-04_TEChi50USAAZSRR12600307 |
| 38 | M_mendax-03_JTOrd05USAAZSRR12600448 | 93 | M_mendax-04_TEJGJTMor01USAAZSRR12600299 |
| 39 | M_mendax-03_JTOrd08USAAZSRR12600447 | 94 | M_mendax-04_TEJGJTMor06USAAZSRR12600297 |
| 40 | M_mendax-03_JTSyc03USAAZSRR12600446 | 95 | M_mendax-04_TEJGJTMor08USAAZSRR12600296 |
| 41 | M_mendax-03_KRWnoMEXSOSRR12600445 | 96 | M_mendax-04_TEJGJTSky03USAAZSRR12600295 |
| 42 | M_mendax-03_MHSup03USAAZSRR12600444 | 97 | M_mendax-04_TEJGJTSky06USAAZSRR12600294 |
| 43 | M_mendax-03_PSW14509USAAZSRR12600443 | 98 | M_mendax-04_TEJGJTSky13USAAZSRR12600293 |
| 44 | M_mendax-03_TECOCed01USAAZSRR12600332 | 99 | M_mendax-04_WPM16472USANMSRR12600292 |
| 45 | M_mendax-03_TECOCed02USAAZSRR12600330 | 100 | M_mendax-04_WPM16702USANMSRR12600291 |
| 46 | M_mendax-03_TEDon01USAAZSRR12600352 | 101 | M_mendax-04_WPM16704USANMSRR12600290 |
| 47 | M_mendax-03_TEDri01USAAZSRR12600351 | 102 | M_mendax-04_WPM19359USACOSRR12600289 |
| 48 | M_mendax-03_TEPat01USAAZSRR12600350 | 103 | M_mendax-04_WPMnoUSANMSRR12600288 |
| 49 | M_mendax-03_TEPat08USAAZSRR12600349 | 112 | M_placodops-01_ELVSDPnoUSATXSRR12600437 |
| 50 | M_mendax-03_TEPat09USAAZSRR12600348 | 113 | M_placodops-01_PSW15596USATXSRR12600436 |
| 51 | M_mendax-03_TEPcr01USAAZSRR12600347 | 114 | M_placodops-01_RWPnoUSATXSRR12600435 |
| 52 | M_mendax-03_TEPcr10USAAZSRR12600346 | 115 | M_placodops-02_GCSnoUSAAZSRR12600434 |
| 53 | M_mendax-03_TEPin04USAAZSRR12600345 | 116 | M_placodops-02_KRWnoMEXSOSRR12600433 |
| 54 | M_mendax-03_TEPin07USAAZSRR12600344 | 117 | M_placodops-02_PSW16232USAAZSRR12600432 |

| | | | |
|---|---|---|---|
| 55 | M_mendax-03_TEPin14USAAZSRR12600343 | 118 | M_placodops-02_RRS7712USACASRR12600431 |
| 56 | M_mendax-03_TESch01USAAZSRR12600341 | 119 | M_placodops-02_TEBSNv02USANVSRR12600429 |
| 57 | M_mendax-03_TESch07USAAZSRR12600340 | 120 | M_placodops-02_TEBSNv03USANVSRR12600427 |
| 58 | M_mendax-03_TESch11USAAZSRR12600339 | 121 | M_placodops-02_TESou01USAAZSRR12600430 |
| 59 | M_mendax-03_TESie05USAAZSRR12600338 | 122 | M_placodops-03_PSW1498103USAAZSRR12600426 |
| 60 | M_mendax-03_TESie126USAAZSRR12600337 | 123 | M_placodops-03_PSW15153USACASRR12600425 |
| 61 | M_mendax-03_TESie143USAAZSRR12600336 | 124 | M_placodops-03_RAJ2978USAAZSRR12600424 |
| 62 | M_mendax-03_TESie151USAAZSRR12600335 | 125 | M_placodops-03_TEBSNv01USANVSRR12600423 |
| 63 | M_mendax-03_TESie20USAAZSRR12600334 | 126 | M_placodops-03_TEBSNv04USANVSRR12600422 |
| 64 | M_mendax-03_TESie72USAAZSRR12600333 | 127 | M_placodops-03_TEBSNv05USANVSRR12600421 |
| 65 | M_mendax-04JGLWChi02USAAZSRR12600329 | 66 | M_mendax-04JGLWChi06USAAZSRR12600328 |

**Tab. A3 Vegetation types and their respective frequency associated with samples within the *M. mendax* cluster 3 to 5 defined in the phylogenomic and population genomic analyses.** Vegetation types retrieved from LANDFIRE (2022, EVT_CONUS).

| Habitat_Type | Cluster3 | Cluster4 | Cluster5 |
|---|---|---|---|
| Chihuahuan Succulent Desert Scrub | 1 | 0 | 0 |
| Colorado Plateau Pinyon-Juniper Woodland | 0 | 3 | 0 |
| Developed-Roads | 6 | 5 | 0 |
| Fill-NoData | 0 | 3 | 0 |
| Inter-Mountain Basins Mixed Salt Desert Scrub | 0 | 1 | 1 |
| Inter-Mountain Basins Semi-Desert Grassland | 0 | 1 | 0 |
| Inter-Mountain Basins Semi-Desert Shrub-Steppe | 0 | 1 | 2 |
| Interior West Ruderal Riparian Forest | 0 | 1 | 0 |
| Madrean Encinal | 4 | 8 | 0 |
| Madrean Lower Montane Pine-Oak Forest and Woodland | 0 | 1 | 0 |
| Madrean Pinyon-Juniper Woodland | 9 | 5 | 0 |
| Mogollon Chaparral | 3 | 2 | 0 |
| North American Warm Desert Bedrock Cliff and Outcrop | 1 | 0 | 0 |
| North American Warm Desert Lower Montane Riparian Woodland | 1 | 9 | 0 |
| North American Warm Desert Ruderal & Planted Scrub | 0 | 1 | 0 |
| Southern Rocky Mountain Pinyon-Juniper Woodland | 0 | 0 | 1 |
| Southern Rocky Mountain Ponderosa Pine Woodland | 1 | 0 | 0 |

**Tab. A4 Sampling and determination history of the samples used in this study.** Colors indicate cluster-assignment based on the phylogenomic and population genetic analyses. The genus name *Myrmecocystus* was shortened to M..

| Sample-ID used in this study and van Elst et al. 2021 | Species according to van Elst et al. 2021 | Species assigned prior to van Elst et al. 2021 | Collection_ID | Collection date | Collector |
|---|---|---|---|---|---|
| Formica_moki_MLB_1000_USA_CA | *Formica moki* | *Formica_moki* | MLB_1000 | May 19 2013 | ML Borowiec |
| Gnamptogenys_simulans_Wa-D-04-2-38_NI | *Gnamptogenys simulans* | *Gnamptogenys_simulans* | Wa-D-04-2-38 | May 18 2011 | Project Llama |
| Lasius_alienus_RAJ_5575_MEX_SO | *Lasius alienus* | *Lasius_alienus* | RAJ_5575 | May 1 2016 | RA Johnson |
| Lasius_arizonicus_RAJ_5576_MEX_SO | *Lasius arizonicus* | *Lasius_arizonicus* | RAJ_5576 | May 1 2016 | RA Johnson |
| Lasius_colei_JT_Lc-01_USA_AZ | *Lasius colei* | *Lasius_colei* | Lc-01 | May 17 2016 | JE Taylor |
| Lasius_fuliginosus_MLB_2009 | *Lasius fuliginosus* | *Lasius_fuliginosus* | MLB_2009 | NA | ML Borowiec |
| Lasius_sitiens_JTL_8439_USA_UT | *Lasius sitiens* | *Lasius_sitiens* | JTL_8439 | Nov 10 2013 | JT Longino |
| Lasius_umbratus_MLB_145_PO_LS | *Lasius umbratus* | *Lasius_umbratus* | MLB_145 | Aug 15 2010 | ML Borowiec |
| M_arenarius_PSW_14630_USA_NV | *M. arenarius* | *M_arenarius* | PSW_14630 | Jun 30 2002 | PS Ward |
| M_arenarius_RCBRLB_no#_USA_NV | *M. arenarius* | *M_arenarius* | NA | Apr 4 1979 | RC Bechtel & RL Bradlay |
| M_christineae_RAJ_6029_USA_CA | *M. christineae* | *M_christineae* | RAJ_6029 | May 2 2018 | RA Johnson |
| M_christineae_RAJ_6035_USA_NV | *M. christineae* | *M_christineae* | RAJ_6035 | May 2 2018 | RA Johnson |
| M_christineae_RRS_77-10,1_USA_CA | *M. christineae* | *M_christineae* | RRS_77-10.1 | Apr 14 1977 | RR Snelling |
| M_colei_RRS_69-118_USA_CA | *M. colei* | *M_colei* | RRS_69-118 | Apr 21 1969 | RR Snelling |
| M_colei_TVE_52_USA_CA | *M. colei* | *M_colei* | TVE_52 | May 31 2018 | T van Elst |
| M_creightoni_GCWJNW_NEV862_USA_NV | *M. creightoni* | *M_creightoni* | NEV_862 | May 10 1970 | GC & JN Wheeler |
| M_creightoni_PSW_17805_USA_CA | *M. creightoni* | *M_creightoni* | PSW_17805 | Apr 9 2017 | PS Ward |
| M_creightoni_RAJ_961_USA_CA | *M. creightoni* | *M_creightoni* | RAJ_961 | Apr 1 1997 | RA Johnson |
| M_depilis_CAK_1_USA_NM | *M. depilis* | *M_depilis* | NA | May 17 1972 | CA Kay |
| M_depilis_PSW_15480_USA_NM | *M. depilis* | *M_depilis* | PSW_15480 | Aug 8 2005 | PS Ward |
| M_depilis_SPC_6242_USA_NM | *M. depilis* | *M_depilis* | SPC_6242 | Apr 9 2001 | SP Cover |
| M_ewarti_RRS_no#_USA_CA | *M. ewarti* | *M_ewarti* | NA | Mar 01 1964 | RR Snelling |
| M_flaviceps_PSW_15180_USA_CA | *M. flaviceps* | *M_flaviceps* | PSW_15180 | Mar 29 2004 | PS Ward |
| M_flaviceps_PSW_15424_MEX_BC | *M. flaviceps* | *M_flaviceps* | PSW_15424 | Mar 25 2005 | PS Ward |
| M_flaviceps_RAJ_2727_MEX_BC | *M. flaviceps* | *M_flaviceps* | RAJ_2727 | Mar 20 2002 | RA Johnson |
| M_flaviceps_RAJ_2983_MEX_BC | *M. flaviceps* | *M_flaviceps* | RAJ_2983 | Mar 10 2003 | RA Johnson |
| M_flaviceps_RRS_94-116_USA_CA | *M. flaviceps* | *M_flaviceps* | RRS_94-116 | Apr 8 1994 | RR Snelling |
| M_flaviceps_TVE_49_USA_CA | *M. flaviceps* | *M_flaviceps* | TVE_49 | May 30 2018 | T van Elst |
| M_romainei_PSW_16759_USA_NV | *M. romainei* | *M_flaviceps* | PSW_16759 | Jul 3 2012 | PS Ward |
| M_sp_cf_mimicus-flaviceps-01_PSW_10552_MEX_BC | *M.* sp. cf. *mimicus-flaviceps*-01 | *M_flaviceps* | PSW_10552 | Jan 4 1990 | PS Ward |
| M_sp_cf_mimicus-flaviceps-01_RAJ_6025_USA_CA | *M.* sp. cf. *mimicus-flaviceps*-01 | *M_flaviceps* | RAJ_6025 | Apr 30 2018 | RA Johnson |
| M_sp_cf_mimicus-flaviceps-03_MLB_1544_USA_AZ | *M.* sp. cf. *mimicus-flaviceps*-03 | *M_flaviceps* | MLB_1544 | Apr 15 2018 | ML Borowiec |
| M_sp_cf_mimicus-flaviceps-03_SPC_4846_USA_AZ | *M.* sp. cf. *mimicus-flaviceps*-03 | *M_flaviceps* | SPC_4846 | Apr 4 1997 | SP Cover |
| M_hammettensis_GCWJNW_NEV1338_USA_NV | *M. hammettensis* | *M_hammettensis* | NEV_1338 | Jul 15 1970 | GC & JN Wheeler |
| M_intonsus_RRS_69-71_MEX_BCS | *M. intonsus* | *M_intonsus* | RRS_69-71 | Mar 2 1969 | RR Snelling |
| M_kathjuli_JPDKED_no#_USA_CA | *M. kathjuli* | *M_kathjuli* | NA | Mar 31 1972 | JP & KE Do-hue |
| M_kathjuli_PSW_14795_USA_CA | *M. kathjuli* | *M_kathjuli* | PSW_14795 | Mar 16 2003 | PS Ward |
| M_kathjuli_RAJ_6007_USA_CA | *M. kathjuli* | *M_kathjuli* | RAJ_6007 | Apr 26 2018 | RA Johnson |
| M_kathjuli_RAJ_6022_USA_CA | *M. kathjuli* | *M_kathjuli* | RAJ_6022 | Apr 30 2018 | RA Johnson |
| M_kennedyi_GCSJH_98-065_USA_CA | *M. kennedyi* | *M_kennedyi* | 98-065 | Jun 6 1998 | GC Snelling & J Hogue |
| M_kennedyi_RAJ_1249_MEX_BC | *M. kennedyi* | *M_kennedyi* | RAJ_1249 | Mar 5 1998 | RA Johnson |
| M_kennedyi_RAJ_2725_MEX_BC | *M. kennedyi* | *M_kennedyi* | RAJ_2725 | Mar 19 2002 | RA Johnson |
| M_kennedyi_RAJ_5778_MEX_SO | *M. kennedyi* | *M_kennedyi* | RAJ_5778 | Oct 29 2016 | RA Johnson |
| M_kennedyi_RAJ_910-1_USA_AZ | *M. kennedyi* | *M_kennedyi* | RAJ_910-1 | Apr 4 1997 | RA Johnson |
| M_sp_cf_kennedyi-romainei_MAJO_15_USA_NV | *M.* sp. cf. *kennedyi-romainei* | *M_kennedyi* | O_15 | May 26 2013 | MA Jansen & Obrien |
| M_koso_RRS_67-274_USA_CA | *M. koso* | *M_koso* | RRS_67-274 | Nov 3 1967 | RR Snelling |
| M_sp_cf_kennedyi-romainei_PSW_15850_USA_CA | *M.* sp. cf. *kennedyi-romainei* | *M_koso* | PSW_15850 | Mar 23 2007 | PS Ward |
| M_lugubris_GHA_no#_USA_CA | *M. lugubris* | *M_lugubris* | NA | Apr 01 1978 | Giulani, Hardy & Andrews |
| M_melanoticus_LBC_no#_MEX_PU | *M. melanoticus* | *M_melanoticus* | NA | Jun 29 1961 | LB Carney |
| M_sp_cf_melliger_DMO_no#_MEX_CO | *M.* sp. cf. *melliger* | *M_melliger* | NA | Jun 2 1998 | DM Olson |
| M_sp_cf_melliger_JCA_2121_USA_TX | *M.* sp. cf. *melliger* | *M_melliger* | JCA_2121 | May 15-19 2005 | JC Abott |
| M_sp_cf_melliger_RRS_no#_MEX_PU | *M.* sp. cf. *melliger* | *M_melliger* | NA | Jul 16 1965 | RR Snelling |

| | | | | | |
|---|---|---|---|---|---|
| M_sp_cf_melliger_WPEM_17561_USA_TX | *M.* sp. cf. *melliger* | *M_melliger* | WM_17561 | Aug 3 1777 | WP & E MacKay |
| M_sp_cf_melliger_WPM_5148_MEX_CH | *M.* sp. cf. *melliger* | *M_melliger* | WM_5148 | Jun 27 1981 | WP MacKay |
| M_sp_cf_mendax-01_JDH_no#_USA_TX | *M.* sp. cf. *mendax*-01 | *M_melliger* | NA | Sep 15 2005 | JD Harrison |
| M_placodops_SS_no#_USA_CA | *M.* sp. cf. *mendax*-02 | *M_mendax* | NA | Apr 15 1990 | S Schoening |
| M_semirufus_TEBS_Ca-01_USA_CA | *M. semirufus* | *M_mendax* | Ca-01 | May 13 2017 | TH Eriksson & B Siqueiros |
| M_semirufus_TEBS_Ca-02_USA_CA | *M. semirufus* | *M_mendax* | Ca-02 | May 13 2017 | TH Eriksson & B Siqueiros |
| M_sp_cf_melliger_JT_Bue-01_MEX_SO | *M.* sp. cf. *melliger* | *M_mendax* | Bue-01 | Aug 16 2016 | JE Taylor |
| M_sp_cf_melliger_JT_Bue-09_MEX_SO | *M.* sp. cf. *melliger* | *M_mendax* | Bue-09 | Aug 14 2016 | JE Taylor |
| M_sp_cf_melliger_JT_Hop-11_USA_AZ | *M.* sp. cf. *melliger* | *M_mendax* | Hop-11 | Apr 11 2015 | JE Taylor |
| M_sp_cf_melliger_RAJ_5588_MEX_SO | *M.* sp. cf. *melliger* | *M_mendax* | RAJ_5588 | May 1 2016 | RA Johnson |
| M_sp_cf_mendax-01_FMH_1305_USA_TX | *M.* sp. cf. *mendax*-01 | *M_mendax* | FMH_1305 | Mar 23 1978 | OF Francke, JV Moody & JB Hall |
| M_sp_cf_mendax-01_FMH_no#_USA_TX | *M.* sp. cf. *mendax*-01 | *M_mendax* | NA | Mar 23 1978 | OF Francke, JV Moody & JB Hall |
| M_sp_cf_mendax-01_WPM_13384_USA_TX | *M.* sp. cf. *mendax*-01 | *M_mendax* | WM_13384 | Mar 15 1990 | WP MacKay |
| M_sp_cf_mendax-03_JT_Fre-02_USA_AZ | *M.* sp. cf. *mendax*-03 | *M_mendax* | Fre-02 | Jun 6 2015 | JE Taylor |
| M_sp_cf_mendax-03_JT_Hop-03_USA_AZ | *M.* sp. cf. *mendax-03* | *M_mendax* | Hop-03 | Apr 11 2015 | JE Taylor |
| M_sp_cf_mendax-03_JT_Mol-02_USA_AZ | *M.* sp. cf. *mendax*-03 | *M_mendax* | Mol-02 | May 2 2015 | JE Taylor |
| M_sp_cf_mendax-03_JT_Mol-03_USA_AZ | *M.* sp. cf. *mendax*-03 | *M_mendax* | Mol-03 | May 2 2015 | JE Taylor |
| M_sp_cf_mendax-03_JT_Ord-03_USA_AZ | *M.* sp. cf. *mendax*-03 | *M_mendax* | Ord-03 | Jul 25 2015 | JE Taylor |
| M_sp_cf_mendax-03_JT_Ord-05_USA_AZ | *M.* sp. cf. *mendax*-03 | *M_mendax* | Ord-05 | Jul 25 2015 | JE Taylor |
| M_sp_cf_mendax-03_JT_Ord-08_USA_AZ | *M.* sp. cf. *mendax*-03 | *M_mendax* | Ord-08 | Jul 25 2015 | JE Taylor |
| M_sp_cf_mendax-03_JT_Syc-03_USA_AZ | *M.* sp. cf. *mendax*-03 | *M_mendax* | Syc-03 | Mar 14 2015 | JE Taylor |
| M_sp_cf_mendax-03_KRW_no#_MEX_SO | *M.* sp. cf. *mendax*-03 | *M_mendax* | NA | Mar 15 1998 | KR Walker |
| M_sp_cf_mendax-03_MH_Sup-03_USA_AZ | *M.* sp. cf. *mendax*-03 | *M_mendax* | Sup-03 | Mar 2012 | M Helmkampf |
| M_sp_cf_mendax-03_PSW_14509_USA_AZ | *M.* sp. cf. *mendax*-03 | *M_mendax* | PSW_14509 | Oct 13 2001 | PS Ward |
| M_sp_cf_mendax-03_TE_Don-01_USA_AZ | *M.* sp. cf. *mendax*-03 | *M_mendax* | Don-01 | Jun 5 2016 | TH Eriksson |
| M_sp_cf_mendax-03_TE_Dri-01_USA_AZ | *M.* sp. cf. *mendax*-03 | *M_mendax* | Dri-01 | May 16 2016 | TH Eriksson |
| M_sp_cf_mendax-03_TE_Pat-01_USA_AZ | *M.* sp. cf. *mendax*-03 | *M_mendax* | Pat-01 | Apr 26 2013 | TH Eriksson |
| M_sp_cf_mendax-03_TE_Pat-08_USA_AZ | *M.* sp. cf. *mendax*-03 | *M_mendax* | Pat-08 | Jul 16 2016 | TH Eriksson |
| M_sp_cf_mendax-03_TE_Pat-09_USA_AZ | *M.* sp. cf. *mendax*-03 | *M_mendax* | Pat-09 | Jul 16 2016 | TH Eriksson |
| M_sp_cf_mendax-03_TE_Pcr-01_USA_AZ | *M.* sp. cf. *mendax*-03 | *M_mendax* | Pcr-01 | Aug 12 2017 | TH Eriksson |
| M_sp_cf_mendax-03_TE_Pcr-10_USA_AZ | *M.* sp. cf. *mendax*-03 | *M_mendax* | Pcr-10 | Aug 12 2017 | TH Eriksson |
| M_sp_cf_mendax-03_TE_Pin-04_USA_AZ | *M.* sp. cf. *mendax*-03 | *M_mendax* | Pin-04 | Mar 11 2015 | TH Eriksson |
| M_sp_cf_mendax-03_TE_Pin-07_USA_AZ | *M.* sp. cf. *mendax*-03 | *M_mendax* | Pin-07 | Mar 11 2015 | TH Eriksson |
| M_sp_cf_mendax-03_TE_Pin-14_USA_AZ | *M.* sp. cf. *mendax*-03 | *M_mendax* | Pin-14 | Mar 11 2015 | TH Eriksson |
| M_sp_cf_mendax-03_TE_Sch-01_USA_AZ | *M.* sp. cf. *mendax*-03 | *M_mendax* | Sch-01 | Jun 4 2016 | TH Eriksson |
| M_sp_cf_mendax-03_TE_Sch-07_USA_AZ | *M.* sp. cf. *mendax*-03 | *M_mendax* | Sch-07 | Jun 4 2016 | TH Eriksson |
| M_sp_cf_mendax-03_TE_Sch-11_USA_AZ | *M.* sp. cf. *mendax*-03 | *M_mendax* | Sch-11 | Jun 4 2016 | TH Eriksson |
| M_sp_cf_mendax-03_TE_Sie-05_USA_AZ | *M.* sp. cf. *mendax*-03 | *M_mendax* | Sie-05 | Jan 07 2013 | TH Eriksson |
| M_sp_cf_mendax-03_TE_Sie-126_USA_AZ | *M.* sp. cf. *mendax*-03 | *M_mendax* | Sie-126 | Jun 4 2016 | TH Eriksson |
| M_sp_cf_mendax-03_TE_Sie-143_USA_AZ | *M.* sp. cf. *mendax*-03 | *M_mendax* | Sie-143 | Jun 4 2016 | TH Eriksson |
| M_sp_cf_mendax-03_TE_Sie-151_USA_AZ | *M.* sp. cf. *mendax*-03 | *M_mendax* | Sie-151 | Jun 4 2016 | TH Eriksson |
| M_sp_cf_mendax-03_TE_Sie-20_USA_AZ | *M.* sp. cf. *mendax*-03 | *M_mendax* | Sie-20 | Jan 07 2013 | TH Eriksson |
| M_sp_cf_mendax-03_TE_Sie-72_USA_AZ | *M.* sp. cf. *mendax*-03 | *M_mendax* | Sie-72 | Jun 24 2013 | TH Eriksson |
| M_sp_cf_mendax-03_TECO_Ced-01_USA_AZ | *M.* sp. cf. *mendax*-03 | *M_mendax* | Ced-01 | Jul 18 2016 | TH Eriksson & C d'Orgiex |
| M_sp_cf_mendax-03_TECO_Ced-02_USA_AZ | *M.* sp. cf. *mendax*-03 | *M_mendax* | Ced-02 | Jul 18 2016 | TH Eriksson & C d'Orgiex |
| M_sp_cf_mendax-04_JGLW_Chi-02_USA_AZ | *M.* sp. cf. *mendax*-04 | *M_mendax* | Chi-02 | Aug 6 2011 | J Gadau & L Wissler |
| M_sp_cf_mendax-04_JGLW_Chi-06_USA_AZ | *M.* sp. cf. *mendax*-04 | *M_mendax* | Chi-06 | Aug 6 2011 | J Gadau & L Wissler |
| M_sp_cf_mendax-04_JGLW_Chi-08_USA_AZ | *M.* sp. cf. *mendax*-04 | *M_mendax* | Chi-08 | Aug 6 2011 | J Gadau & L Wissler |
| M_sp_cf_mendax-04_JT_Bue-02_MEX_SO | *M.* sp. cf. *mendax*-04 | *M_mendax* | Bue-02 | Mar 7 2016 | JE Taylor |
| M_sp_cf_mendax-04_JT_Bue-04_MEX_SO | *M.* sp. cf. *mendax*-04 | *M_mendax* | Bue-04 | Mar 7 2016 | JE Taylor |
| M_sp_cf_mendax-04_JT_Bue-11_MEX_SO | *M.* sp. cf. *mendax*-04 | *M_mendax* | Bue-11 | Aug 16 2016 | JE Taylor |
| M_sp_cf_mendax-04_JT_Car-01_USA_AZ | *M.* sp. cf. *mendax*-04 | *M_mendax* | Car-01 | May 16 2015 | JE Taylor |
| M_sp_cf_mendax-04_JT_Coc-02_USA_AZ | *M.* sp. cf. *mendax*-04 | *M_mendax* | Coc-02 | June 13 2015 | JE Taylor |
| M_sp_cf_mendax-04_JT_Coc-03_USA_AZ | *M.* sp. cf. *mendax*-04 | *M_mendax* | Coc-03 | June 13 2015 | JE Taylor |
| M_sp_cf_mendax-04_JT_Coc-10_USA_AZ | *M.* sp. cf. *mendax*-04 | *M_mendax* | Coc-10 | June 13 2015 | JE Taylor |
| M_sp_cf_mendax-04_JT_Fre-04_USA_AZ | *M.* sp. cf. *mendax*-04 | *M_mendax* | Fre-04 | Jun 6 2015 | JE Taylor |
| M_sp_cf_mendax-04_JT_Fre-05_USA_AZ | *M.* sp. cf. *mendax*-04 | *M_mendax* | Fre-05 | Jun 6 2015 | JE Taylor |
| M_sp_cf_mendax-04_JT_Kit-01_USA_AZ | *M.* sp. cf. *mendax*-04 | *M_mendax* | Kit-01 | March 12 2016 | JE Taylor |

| | | | | | |
|---|---|---|---|---|---|
| M_sp_cf_mendax-04_JT_Mil-01_USA_AZ | *M.* sp. cf. *mendax*-04 | *M_mendax* | Mil-01 | May 16 2015 | JE Taylor |
| M_sp_cf_mendax-04_JT_Ram-01_USA_AZ | *M.* sp. cf. *mendax*-04 | *M_mendax* | Ram-01 | May 16 2015 | JE Taylor |
| M_sp_cf_mendax-04_PSW_16919-01_MEX_CH | *M.* sp. cf. *mendax*-04 | *M_mendax* | PSW_16919-01 | Mar 29 2013 | PS Ward |
| M_sp_cf_mendax-04_RAJ_2536_MEX_CH | *M.* sp. cf. *mendax*-04 | *M_mendax* | RAJ_2536 | Aug 4 2001 | RA Johnson |
| M_sp_cf_mendax-04_RAJ_2800_MEX_CH | *M.* sp. cf. *mendax*-04 | *M_mendax* | RAJ_2800 | Sep 18 1993 | RA Johnson |
| M_sp_cf_mendax-04_RAJ_4930_USA_AZ | *M.* sp. cf. *mendax*-04 | *M_mendax* | RAJ_4930 | Aug 5 2012 | RA Johnson |
| M_sp_cf_mendax-04_TE_Chi-50_USA_AZ | *M.* sp. cf. *mendax*-04 | *M_mendax* | Chi-50 | Jul 16 2014 | TH Eriksson |
| M_sp_cf_mendax-04_TEBS_Co-01_USA_CO | *M.* sp. cf. *mendax*-04 | *M_mendax* | Co-01 | May 8 2017 | TH Eriksson & B Siqueiros |
| M_sp_cf_mendax-04_TEBS_Nm-01_USA_NM | *M.* sp. cf. *mendax*-04 | *M_mendax* | Nm-01 | May 7 2017 | TH Eriksson & B Siqueiros |
| M_sp_cf_mendax-04_TEBS_Nm-02_USA_NM | *M.* sp. cf. *mendax*-04 | *M_mendax* | Nm-02 | May 7 2017 | TH Eriksson & B Siqueiros |
| M_sp_cf_mendax-04_TEBS_Nm-03_USA_NM | *M.* sp. cf. *mendax*-04 | *M_mendax* | Nm-03 | May 7 2017 | TH Eriksson & B Siqueiros |
| M_sp_cf_mendax-04_TEBS_Nm-04_USA_NM | *M.* sp. cf. *mendax*-04 | *M_mendax* | Nm-04 | May 7 2017 | TH Eriksson & B Siqueiros |
| M_sp_cf_mendax-04_TEBS_Nm-05_USA_NM | *M.* sp. cf. *mendax*-04 | *M_mendax* | Nm-05 | May 7 2017 | TH Eriksson & B Siqueiros |
| M_sp_cf_mendax-04_TEBS_Nm-08_USA_NM | *M.* sp. cf. *mendax*-04 | *M_mendax* | Nm-08 | May 7 2017 | TH Eriksson & B Siqueiros |
| M_sp_cf_mendax-04_TEJGJT_Mor-01_USA_AZ | *M.* sp. cf. *mendax*-04 | *M_mendax* | Mor-01 | May 30 2016 | TH Eriksson, J Gadau & JE Taylor |
| M_sp_cf_mendax-04_TEJGJT_Mor-06_USA_AZ | *M.* sp. cf. *mendax*-04 | *M_mendax* | Mor-06 | May 30 2016 | TH Eriksson, J Gadau & JE Taylor |
| M_sp_cf_mendax-04_TEJGJT_Mor-08_USA_AZ | *M.* sp. cf. *mendax*-04 | *M_mendax* | Mor-08 | May 30 2016 | TH Eriksson, J Gadau & JE Taylor |
| M_sp_cf_mendax-04_TEJGJT_Sky-03_USA_AZ | *M.* sp. cf. *mendax*-04 | *M_mendax* | Sky-03 | May 17 2016 | TH Eriksson, J Gadau & JE Taylor |
| M_sp_cf_mendax-04_TEJGJT_Sky-06_USA_AZ | *M.* sp. cf. *mendax*-04 | *M_mendax* | Sky-06 | May 17 2016 | TH Eriksson, J Gadau & JE Taylor |
| M_sp_cf_mendax-04_TEJGJT_Sky-13_USA_AZ | *M.* sp. cf. *mendax*-04 | *M_mendax* | Sky-13 | May 17 2016 | TH Eriksson, J Gadau & JE Taylor |
| M_sp_cf_mendax-04_WPM_16472_USA_NM | *M.* sp. cf. *mendax*-04 | *M_mendax* | WM_16472 | Mar 22 1994 | WP MacKay |
| M_sp_cf_mendax-04_WPM_16704_USA_NM | *M.* sp. cf. *mendax*-04 | *M_mendax* | WM_16704 | Aug 1 1994 | WP MacKay |
| M_sp_cf_mendax-04_WPM_no#_USA_NM | *M.* sp. cf. *mendax*-04 | *M_mendax* | NA | 1986 | WP MacKay |
| M_sp_cf_placodops-02_RRS_77-12_USA_CA | *M.* sp. cf. *placodops*-02 | *M_mendax* | RRS_77-12 | Apr 14 1977 | RR Snelling |
| M_sp_cf_placodops-02_TEBS_Nv-02_USA_NV | *M.* sp. cf. *placodops*-02 | *M_mendax* | Nv-02 | May 12 2017 | TH Eriksson & B Siqueiros |
| M_sp_cf_placodops-02_TEBS_Nv-03_USA_NV | *M.* sp. cf. *placodops*-02 | *M_mendax* | Nv-03 | May 12 2017 | TH Eriksson & B Siqueiros |
| M_sp_cf_placodops-03_TEBS_Nv-01_USA_NV | *M.* sp. cf. *placodops*-03 | *M_mendax* | Nv-01 | May 11 2017 | TH Eriksson & B Siqueiros |
| M_sp_cf_placodops-03_TEBS_Nv-04_USA_NV | *M.* sp. cf. *placodops*-03 | *M_mendax* | Nv-04 | May 12 2017 | TH Eriksson & B Siqueiros |
| M_sp_cf_placodops-03_TEBS_Nv-05_USA_NV | *M.* sp. cf. *placodops*-03 | *M_mendax* | Nv-05 | May 12 2017 | TH Eriksson & B Siqueiros |
| M_sp_cf_mexicanus-01_RRS_69-298_USA_AZ | *M.* sp. cf. *mexicanus*-01 | *M_mexicanus* | RRS_69-298 | Aug 13 1969 | RR Snelling |
| M_sp_cf_mexicanus-02_MLB_1122_USA_AZ | *M.* sp. cf. *mexicanus*-02 | *M_mexicanus* | MLB_1122 | Apr 17 2014 | ML Borowiec |
| M_sp_cf_mexicanus-02_MLB_421_USA_NV | *M.* sp. cf. *mexicanus*-02 | *M_mexicanus* | MLB_421 | Jul 2 2012 | ML Borowiec |
| M_sp_cf_mexicanus-02_RAJ_3061_MEX_BC | *M.* sp. cf. *mexicanus*-02 | *M_mexicanus* | RAJ_3061 | Mar 17 2003 | RA Johnson |
| M_sp_cf_mexicanus-02_RAJ_6012_USA_CA | *M.* sp. cf. *mexicanus*-02 | *M_mexicanus* | RAJ_6012 | Apr 27 2018 | RA Johnson |
| M_sp_cf_mexicanus-02_RAJ_6028_USA_CA | *M.* sp. cf. *mexicanus*-02 | *M_mexicanus* | RAJ_6028 | May 2 2018 | RA Johnson |
| M_sp_cf_mexicanus-02_SPC_6256_USA_CA | *M.* sp. cf. *mexicanus*-02 | *M_mexicanus* | SPC_6256 | Apr 11 2001 | SP Cover |
| | *M.* sp. cf. *mimicus-flaviceps* | *M_mimicus* | RAJ_1207 | Oct 11 1992 | RA Johnson |
| M_sp_cf_mimicus-flaviceps-01_RAJ_3076_MEX_BC | *M.* sp. cf. *mimicus-flaviceps*-01 | *M_mimicus* | RAJ_3076 | Mar 18 2003 | RA Johnson |
| M_sp_cf_mimicus-flaviceps-01_RRS_77-22_USA_CA | *M.* sp. cf. *mimicus-flaviceps*-01 | *M_mimicus* | RRS_77-22 | Apr 18 1977 | RR Snelling |
| M_sp_cf_mimicus-flaviceps-02_RAJ_2665_MEX_BCS | *M.* sp. cf. *mimicus-flaviceps*-02 | *M_mimicus* | RAJ_2665 | Mar 13 2002 | RA Johnson |
| M_sp_cf_mimicus-flaviceps-02_RAJ_3030_MEX_BCS | *M.* sp. cf. *mimicus-flaviceps*-02 | *M_mimicus* | RAJ_3030 | Mar 15 2003 | RA Johnson |
| M_sp_cf_mimicus-flaviceps-03_JT_Fre-01_USA_AZ | *M.* sp. cf. *mimicus-flaviceps*-03 | *M_mimicus* | Fre-01 | Jun 6 2015 | JE Taylor |
| M_sp_cf_mimicus-flaviceps-03_RAJ_5784_USA_AZ | *M.* sp. cf. *mimicus-flaviceps*-03 | *M_mimicus* | RAJ_5784 | Nov 13 2016 | RA Johnson |
| M_navajo_GCWJNW_NEV892_USA_NV | *M. navajo* | *M_navajo* | NEV_892 | May 17 1970 | GC & JN Wheeler |
| M_navajo_RAJ_2914_USA_AZ | *M. navajo* | *M_navajo* | RAJ_2914 | Aug 12 2002 | RA Johnson |
| M_navajo_RAJ_2986_MEX_BC | *M. navajo* | *M_navajo* | RAJ_2986 | Mar 10 2003 | RA Johnson |
| M_navajo_RAJ_3101_MEX_BC | *M. navajo* | *M_navajo* | RAJ_3101 | Mar 20 2003 | RA Johnson |
| M_navajo_SPC_7476_USA_AZ | *M. navajo* | *M_navajo* | SPC_7476 | Oct 17 2006 | SP Cover |
| M_nequazcatl_EMF_no#_MEX_SO | *M. nequazcatl* | *M_nequazcatl* | NA | Feb 5 1972 | EM Fisher |
| M_nequazcatl_PSW_13493_MEX_SO | *M. nequazcatl* | *M_nequazcatl* | PSW_13493 | Dec 20 1997 | PS Ward |
| M_nequazcatl_RAJ_962_MEX_SO | *M. nequazcatl* | *M_nequazcatl* | RAJ_962 | Nov 29 1996 | RA Johnson |

| | | | | | |
|---|---|---|---|---|---|
| M_nequazcatl_RAJ_SON95-26_MEX_SO | *M. nequazcatl* | *M_nequazcatl* | RAJ_SON_95-26 | May 29 1994 | RA Johnson |
| M_perimeces_EMFb_no#_MEX_BC | *M. perimeces* | *M_perimeces* | NA | Mar 29 1970 | EM Fisher |
| M_sp_cf_mendax-01_AW_6127_USA_TX | *M.* sp. cf. *mendax*-01 | *M_placodops* | AW_6127 | Mar 11 2018 | A Wild |
| M_sp_cf_mendax-04_PSW_13407-6_USA_NM | *M.* sp. cf. *mendax*-04 | *M_placodops* | PSW_13407-6 | Aug 14 1997 | PS Ward |
| M_sp_cf_mendax-04_WPM_16702_USA_NM | *M.* sp. cf. *mendax*-04 | *M_placodops* | WM_16702 | Aug 1 1994 | WP MacKay |
| M_sp_cf_mendax-04_WPM_19359_USA_CO | *M.* sp. cf. *mendax*-04 | *M_placodops* | WM_19359 | Jul 13 2000 | WP MacKay |
| M_sp_cf_placodops-01_ELVSDP_no#_USA_TX | *M.* sp. cf. *placodops*-01 | *M_placodops* | NA | Mar 28 1987 | EL Vargo & SD Porter |
| M_sp_cf_placodops-01_PSW_15596_USA_TX | *M.* sp. cf. *placodops*-01 | *M_placodops* | PSW_15596 | Apr 14 2006 | PS Ward |
| M_sp_cf_placodops-01_RWP_no#_USA_TX | *M.* sp. cf. *placodops*-01 | *M_placodops* | NA | Oct 12-13 2004 | RW Patrock |
| M_sp_cf_placodops-02_GCS_no#_USA_AZ | *M.* sp. cf. *placodops*-02 | *M_placodops* | NA | Jul 29 1986 | GC Snelling |
| M_sp_cf_placodops-02_KRW_no#_MEX_SO | *M.* sp. cf. *placodops*-02 | *M_placodops* | NA | Mar 14 1998 | KR Walker |
| M_sp_cf_placodops-02_PSW_16232_USA_AZ | *M.* sp. cf. *placodops*-02 | *M_placodops* | PSW_16232 | Apr 4 2009 | PS Ward |
| M_sp_cf_placodops-02_TE_Sou-01_USA_AZ | *M.* sp. cf. *placodops*-02 | *M_placodops* | Sou-01 | Sep 12 2015 | TH Eriksson |
| M_sp_cf_placodops-03_PSW_14981-03_USA_AZ | *M.* sp. cf. *placodops*-03 | *M_placodops* | PSW_14981-03 | Sep 6 2003 | PS Ward |
| M_sp_cf_placodops-03_PSW_15153_USA_CA | *M.* sp. cf. *placodops*-03 | *M_placodops* | PSW_15153 | Mar 27 2004 | PS Ward |
| M_sp_cf_placodops-03_RAJ_2978_USA_AZ | *M.* sp. cf. *placodops*-03 | *M_placodops* | RAJ_2978 | Mar 8 2003 | RA Johnson |
| M_pyramicus_GCWJNW_NEV1304_USA_NV | *M. pyramicus* | *M_pyramicus* | NEV_1304 | Jul 14 1970 | GC & JN Wheeler |
| M_pyramicus_PSW_16114_USA_CA | *M. pyramicus* | *M_pyramicus* | PSW_16114 | Jun 7 2008 | PS Ward |
| M_romainei_MAJO_71_USA_UT | *M. romainei* | *M_romainei* | O_71 | June 9 2013 | MA Jansen & Obrien |
| M_romainei_MLB_1543_USA_AZ | *M. romainei* | *M_romainei* | MLB_1543 | Mar 21 2018 | ML Borowiec |
| M_romainei_PSW_15174_USA_CA | *M. romainei* | *M_romainei* | PSW_15174 | Mar 28 2004 | PS Ward |
| M_romainei_RAJ_6039_USA_CA | *M. romainei* | *M_romainei* | RAJ_6039 | May 3 2018 | RA Johnson |
| M_romainei_RRS_73-47_USA_TX | *M. romainei* | *M_romainei* | RRS_73-47 | Apr 23 1973 | RR Snelling |
| M_romainei_SPC_6239_USA_AZ | *M. romainei* | *M_romainei* | SPC_6239 | Apr 9 2001 | SP Cover |
| M_sp_cf_kennedyi-romainei_RAJ_6009_USA_CA | *M.* sp. cf. *kennedyi-romainei* | *M_romainei* | RAJ_6009 | Apr 27 2018 | RA Johnson |
| M_semirufus_CG_no#_USA_CA | *M. semirufus* | *M_semirufus* | NA | Apr 8 1972 | C Gross |
| M_semirufus_PSW_14281_MEX_BC | *M. semirufus* | *M_semirufus* | PSW_14281 | Mar 29 2001 | PS Ward |
| M_semirufus_PSW_14810_USA_CA | *M. semirufus* | *M_semirufus* | PSW_14810 | Mar 18 2003 | PS Ward |
| M_semirufus_RAJ_2984_MEX_BC | *M. semirufus* | *M_semirufus* | RAJ_2984 | Mar 10 2003 | RA Johnson |
| M_semirufus_RAJ_BC1310_MEX_BC | *M. semirufus* | *M_semirufus* | RAJ_BC_1310 | Mar 11 1998 | RA Johnson |
| M_flaviceps_RAJ_2958_USA_AZ | *M. flaviceps* | *M_sp._(not_identified)* | RAJ_2958 | Jan 24 2003 | RA Johnson |
| M_kennedyi_RAJ_2234_USA_AZ | *M. kennedyi* | *M_sp._(not_identified)* | RAJ_2234 | Mar 21 2001 | RA Johnson |
| M_navajo_RAJ_3139-2_USA_AZ | *M. navajo* | *M_sp._(not_identified)* | RAJ_3139-2 | Aug 11 2003 | RA Johnson |
| M_romainei_RAJ_2349_USA_AZ | *M. romainei* | *M_sp._(not_identified)* | RAJ_2349 | Apr 8 2001 | RA Johnson |
| M_yuma_RAJ_2243_USA_AZ | *M. yuma* | *M_sp._(not_identified)* | RAJ_2243 | Mar 21 2001 | RA Johnson |
| M_sp_SON-1_RAJ_5781_MEX_SO | *M.* sp. SON-1 | *M_sp._(undescribed,_R._Johnson,_pers._obs.)* | RAJ_5781 | Oct 30 2016 | RA Johnson |
| M_sp_cf_colei_RAJ_2655_MEX_BC | *M.* sp. cf. *colei* | *M_sp._BCA-1_(undescribed,_Johnson_and_Ward,_2002)* | RAJ_2655 | Mar 10 2002 | RA Johnson |
| M_sp_cf_mimicus-flaviceps-01_RAJ_2990_MEX_BC | *M.* sp. cf. *mimicus-flaviceps*-01 | *M_sp._cf._flaviceps_(undescribed,_Johnson_and_Ward,_2002)* | RAJ_2990 | Mar 11 2003 | RA Johnson |
| M_sp_cf_mimicus-flaviceps-01_RAJ_3029_MEX_BC | *M.* sp. cf. *mimicus-flaviceps*-01 | *M_sp._cf._flaviceps_(undescribed,_Johnson_and_Ward,_2002)* | RAJ_3029 | Mar 15 2003 | RA Johnson |
| M_sp_cf_kennedyi_RAJ_2810_USA_AZ | *M.* sp. cf. *kennedyi* | *M_sp._cf._kennedyi_(undescribed,_R._Johnson,_pers._obs.)* | RAJ_2810 | May 29 2002 | RA Johnson |
| M_sp_cf_colei_RAJ_2723_MEX_BC | *M.* sp. cf. *mendax*-05 | *M_sp._cf._mendax_(undescribed,_Johnson_and_Ward,_2002)* | RAJ_2723 | Mar 19 2002 | RA Johnson |
| M_sp_cf_navajo_RAJ_3077_MEX_BC | *M.* sp. cf. *navajo* | *M_sp._cf._navajo_(undescribed,_R._Johnson,_pers._obs.)* | RAJ_3077 | Mar 18 2003 | RA Johnson |
| M_tenuinodis_PSW_15178_USA_CA | *M. tenuinodis* | *M_tenuinodis* | PSW_15178 | Mar 29 2004 | PS Ward |
| M_tenuinodis_RRS_no#_USA_CA | *M. tenuinodis* | *M_tenuinodis* | NA | Feb 5 1967 | RR Snelling |
| M_testaceus_MLB_1258_USA_CA | *M. testaceus* | *M_testaceus* | MLB_1258 | Apr 4 2015 | ML Borowiec |
| M_testaceus_MLB_397_USA_NV | *M. testaceus* | *M_testaceus* | MLB_397 | Jun 30 2012 | ML Borowiec |
| M_testaceus_MLB_446_USA_CA | *M. testaceus* | *M_testaceus* | MLB_446 | Jun 10 2012 | ML Borowiec |
| M_testaceus_RAJ_2259_MEX_BC | *M. testaceus* | *M_testaceus* | RAJ_2259 | Mar 23 2001 | RA Johnson |
| M_testaceus_RAJ_6018_USA_CA | *M. testaceus* | *M_testaceus* | RAJ_6018 | Apr 27 2018 | RA Johnson |
| M_testaceus_RRSCDG_77-20_USA_CA | *M. testaceus* | *M_testaceus* | RRS_77-20 | Apr 15 1977 | RR Snelling & CD George |
| M_wheeleri_MLB_1256_USA_CA | *M. wheeleri* | *M_wheeleri* | MLB_1256 | Mar 23 2015 | ML Borowiec |
| M_wheeleri_PSW_14302_USA_CA | *M. wheeleri* | *M_wheeleri* | PSW_14302 | Apr 1 2001 | PS Ward |
| M_wheeleri_RAJ_3094_MEX_BC | *M. wheeleri* | *M_wheeleri* | RAJ_3094 | Mar 20 2003 | RA Johnson |
| M_wheeleri_RAJ_6010_USA_CA | *M. wheeleri* | *M_wheeleri* | RAJ_6010 | Apr 27 2018 | RA Johnson |
| M_wheeleri_RAJ_6013_USA_CA | *M. wheeleri* | *M_wheeleri* | RAJ_6013 | Apr 27 2018 | RA Johnson |

| | | | | | |
|---|---|---|---|---|---|
| M_wheeleri_RAJ_6021_USA_CA | *M. wheeleri* | *M_wheeleri* | RAJ_6021 | Apr 30 2018 | RA Johnson |
| M_wheeleri_RRS_no#_USA_CA | *M. wheeleri* | *M_wheeleri* | NA | Aug 15 1965 | RR Snelling |
| M_tenuinodis_SPC_4826_USA_AZ | *M. tenuinodis* | *M_yuma* | SPC_4826 | Apr 4 1997 | SP Cover |
| M_yuma_PSW_15185_USA_CA | *M. yuma* | *M_yuma* | PSW_15185 | Mar 29 2004 | PS Ward |
| M_yuma_RAJ_3084_MEX_BC | *M. yuma* | *M_yuma* | RAJ_3084 | Mar 20 2003 | RA Johnson |
| M_yuma_RAJ_5783_MEX_SO | *M. yuma* | *M_yuma* | RAJ_5783 | Oct 30 2016 | RA Johnson |
| M_yuma_RRS_94-11a_USA_CA | *M. yuma* | *M_yuma* | RRS_94-11a | Apr 8 1994 | RR Sneling |
| Myrmelachista_joycei_JTL_8492_CR | *Myrmelachista joycei* | *Myrmelachista_ joycei* | JTL_8492 | Dec 27 2013 | JT Longino |
| Nylanderia_terricola_JTL_14763_USA_TX | *Nylanderia terricola* | *Nylanderia_ terricola* | JTL_14763 | Sep 01 1983 | JT Longino |
| Paratrechina_longicornis_LRD_270806-14_USA_FL | *Paratrechina longicornis* | *Paratrechina_ longicornis* | LRD_270806-14 | Aug 27 2006 | LR Davis |
| Polyergus_mexicanus_MLB_389-01_USA_CA | *Polyergus mexicanus* | *Polyergus_ mexicanus* | MLB_389.1 | June 29 2012 | ML Borowiec |

**Tab. A5 Sample and data accession information.** Colors indicate cluster-assignment based on the phylogenomic and population genetic analyses. The genus name *Myrmecocystus* was shortened to M.. Furthermore, the words "University, History, Biodiversity, Repository, unpublished" were also shortened to ensure readability.

| Sample-ID used in van Elst et al. 2021 and this study | Voucher institution | Accession | NCBI BioSample | Note | Reference |
|---|---|---|---|---|---|
| Formica_moki_MLB_1000_USA_CA | Arizona State Uni. Social Insect Biodiv. Rep. | ASU-SIBR00001001 | NA | Outgroup | Borowiec et al., unpubl. |
| Gnamptogenys_simulans_Wa-D-04-2-38_NI | John T. Longino Collection | CASENT0633201 | NA | Outgroup | Branstetter et al., 2017 |
| Lasius_alienus_RAJ_5575_MEX_SO | Arizona State Uni. Social Insect Biodiv. Rep. | ASU-SIBR00001002 | SAMN15830225 | Outgroup | van Elst et al. 2021 |
| Lasius_arizonicus_RAJ_5576_MEX_SO | Arizona State Uni. Social Insect Biodiv. Rep. | ASU-SIBR00001003 | SAMN15830226 | Outgroup | van Elst et al. 2021 |
| Lasius_colei_JT_Lc-01_USA_AZ | Arizona State Uni. Social Insect Biodiv. Rep. | ASU-SIBR00001004 | SAMN15830227 | Outgroup | van Elst et al. 2021 |
| Lasius_fuliginosus_MLB_2009 | Arizona State Uni. Social Insect Biodiv. Rep. | ASU-SIBR00001005 | SAMN15830228 | Outgroup | van Elst et al. 2021 |
| Lasius_sitiens_JTL_8439_USA_UT | John T. Longino Collection | CASENT0635911 | NA | Outgroup | Branstetter et al., 2017 |
| Lasius_umbratus_MLB_145_PO_LS | Arizona State Uni. Social Insect Biodiv. Rep. | ASU-SIBR00001006 | SAMN15830229 | Outgroup | van Elst et al. 2021 |
| M_arenarius_PSW_14630_USA_NV | Arizona State Uni. Social Insect Biodiv. Rep. | ASU-SIBR00001164 | SAMN15830230 | | van Elst et al. 2021 |
| M_arenarius_RCBRLB_no#_USA_NV | Nat. His. Museum of Los Angeles County | LACMENT346514 | SAMN15830231 | Paratype | van Elst et al. 2021 |
| M_christineae_RAJ_6029_USA_CA | Arizona State Uni. Social Insect Biodiv. Rep. | ASU-SIBR00001007 | SAMN15830232 | | van Elst et al. 2021 |
| M_christineae_RAJ_6035_USA_NV | Arizona State Uni. Social Insect Biodiv. Rep. | ASU-SIBR00001009 | SAMN15830233 | | van Elst et al. 2021 |
| M_christineae_RRS_77-10,1_USA_CA | Nat. His. Museum of Los Angeles County | LACMENT346525 | SAMN15830234 | Paratype | van Elst et al. 2021 |
| M_colei_RRS_69-118_USA_CA | Nat. His. Museum of Los Angeles County | LACMENT346515 | SAMN15830235 | | van Elst et al. 2021 |
| M_colei_TVE_52_USA_CA | Arizona State Uni. Social Insect Biodiv. Rep. | ASU-SIBR00001010 | SAMN15830236 | | van Elst et al. 2021 |
| M_creightoni_GCWJNW_NEV862_USA_NV | Nat. His. Museum of Los Angeles County | LACMENT346516 | SAMN15830237 | | van Elst et al. 2021 |
| M_creightoni_PSW_17805_USA_CA | Uni. of California, Davis, Collection | CASENT0842006 | SAMN15830238 | | van Elst et al. 2021 |
| M_creightoni_RAJ_961_USA_CA | Arizona State Uni. Social Insect Biodiv. Rep. | ASU-SIBR00001011 | SAMN15830239 | | van Elst et al. 2021 |
| M_depilis_CAK_1_USA_NM | Nat. His. Museum of Los Angeles County | LACMENT346510 | SAMN15830240 | | van Elst et al. 2021 |
| M_depilis_PSW_15480_USA_NM | Arizona State Uni. Social Insect Biodiv. Rep. | ASU-SIBR00001165 | SAMN15830241 | | van Elst et al. 2021 |
| M_depilis_SPC_6242_USA_NM | Museum of Comparative Zoology | MCZ654623 | SAMN15830242 | | van Elst et al. 2021 |
| M_ewarti_RRS_no#_USA_CA | Nat. His. Museum of Los Angeles County | LACMENT346526 | SAMN15830243 | Paratype | van Elst et al. 2021 |
| M_flaviceps_PSW_15180_USA_CA | Arizona State Uni. Social Insect Biodiv. Rep. | ASU-SIBR00001166 | SAMN15830244 | | van Elst et al. 2021 |
| M_flaviceps_PSW_15424_MEX_BC | Arizona State Uni. Social Insect Biodiv. Rep. | ASU-SIBR00001167 | SAMN15830245 | | van Elst et al. 2021 |
| M_flaviceps_RAJ_2727_MEX_BC | Arizona State Uni. Social Insect Biodiv. Rep. | ASU-SIBR00001013 | SAMN15830246 | | van Elst et al. 2021 |
| M_flaviceps_RAJ_2983_MEX_BC | Arizona State Uni. Social Insect Biodiv. Rep. | ASU-SIBR00001015 | SAMN15830248 | | van Elst et al. 2021 |
| M_flaviceps_RRS_94-116_USA_CA | Nat. His. Museum of Los Angeles County | LACMENT346500 | SAMN15830249 | | van Elst et al. 2021 |
| M_flaviceps_TVE_49_USA_CA | Arizona State Uni. Social Insect Biodiv. Rep. | ASU-SIBR00001016 | SAMN15830250 | | van Elst et al. 2021 |
| M_romainei_PSW_16759_USA_NV | Arizona State Uni. Social Insect Biodiv. Rep. | ASU-SIBR00001172 | SAMN15830282 | | van Elst et al. 2021 |
| M_sp_cf_mimicus-flaviceps-01_ PSW_10552_MEX_BC | Arizona State Uni. Social Insect Biodiv. Rep. | ASU-SIBR00001181 | SAMN15830393 | | van Elst et al. 2021 |
| M_sp_cf_mimicus-flaviceps-01_ RAJ_6025_USA_CA | Arizona State Uni. Social Insect Biodiv. Rep. | ASU-SIBR00001129 | SAMN15830397 | | van Elst et al. 2021 |
| M_sp_cf_mimicus-flaviceps-03_ MLB_1544_USA_AZ | Arizona State Uni. Social Insect Biodiv. Rep. | ASU-SIBR00001133 | SAMN15830402 | | van Elst et al. 2021 |
| M_sp_cf_mimicus-flaviceps-03_ SPC_4846_USA_AZ | Museum of Comparative Zoology | MCZ661690 | SAMN15830404 | | van Elst et al. 2021 |
| M_hammettensis_GCWJNW_ NEV1338_USA_NV | Nat. His. Museum of Los Angeles County | LACMENT346517 | SAMN15830251 | | van Elst et al. 2021 |
| M_intonsus_RRS_69-71_MEX_BCS | Nat. His. Museum of Los Angeles County | LACMENT346505 | SAMN15830252 | Paratype | van Elst et al. 2021 |
| M_kathjuli_JPDKED_no#_USA_CA | Nat. His. Museum of Los Angeles County | LACMENT346501 | SAMN15830253 | Paratype | van Elst et al. 2021 |
| M_kathjuli_PSW_14795_USA_CA | Arizona State Uni. Social Insect Biodiv. Rep. | ASU-SIBR00001168 | SAMN15830254 | | van Elst et al. 2021 |
| M_kathjuli_RAJ_6007_USA_CA | Arizona State Uni. Social Insect Biodiv. Rep. | ASU-SIBR00001017 | SAMN15830255 | | van Elst et al. 2021 |

| | | | | | |
|---|---|---|---|---|---|
| M_kathjuli_RAJ_6022_USA_CA | Arizona State Uni. Social Insect Biodiv. Rep. | ASU-SIBR00001018 | SAMN15830256 | | van Elst et al. 2021 |
| M_kennedyi_GCSJH_98-065_USA_CA | Nat. His. Museum of Los Angeles County | LACMENT346502 | SAMN15830257 | | van Elst et al. 2021 |
| M_kennedyi_RAJ_1249_MEX_BC | Arizona State Uni. Social Insect Biodiv. Rep. | ASU-SIBR00001020 | SAMN15830258 | | van Elst et al. 2021 |
| M_kennedyi_RAJ_2725_MEX_BC | Arizona State Uni. Social Insect Biodiv. Rep. | ASU-SIBR00001022 | SAMN15830260 | | van Elst et al. 2021 |
| M_kennedyi_RAJ_5778_MEX_SO | Arizona State Uni. Social Insect Biodiv. Rep. | ASU-SIBR00001023 | SAMN15830261 | | van Elst et al. 2021 |
| M_kennedyi_RAJ_910-1_USA_AZ | Arizona State Uni. Social Insect Biodiv. Rep. | ASU-SIBR00001024 | SAMN15830262 | | van Elst et al. 2021 |
| M_sp_cf_kennedyi-romainei_ MAJO_15_USA_NV | Arizona State Uni. Social Insect Biodiv. Rep. | ASU-SIBR00001019 | SAMN15830296 | | van Elst et al. 2021 |
| M_koso_RRS_67-274_USA_CA | Nat. His. Museum of Los Angeles County | LACMENT346512 | SAMN15830263 | Paratype | van Elst et al. 2021 |
| M_sp_cf_kennedyi-romainei_ PSW_15850_USA_CA | Arizona State Uni. Social Insect Biodiv. Rep. | ASU-SIBR00001175 | SAMN15830297 | | van Elst et al. 2021 |
| M_lugubris_GHA_no#_USA_CA | Nat. His. Museum of Los Angeles County | LACMENT346518 | SAMN15830264 | | van Elst et al. 2021 |
| M_melanoticus_LBC_no#_MEX_PU | Nat. His. Museum of Los Angeles County | LACMENT346522 | SAMN15830265 | Homotype | van Elst et al. 2021 |
| M_sp_cf_melliger_DMO_no#_MEX_CO | Uni. of California, Davis, Collection | CASENT280506 | SAMN15830299 | | van Elst et al. 2021 |
| M_sp_cf_melliger_JCA_2121_USA_TX | Arizona State Uni. Social Insect Biodiv. Rep. | ASU-SIBR00001163 | SAMN15830300 | | van Elst et al. 2021 |
| M_sp_cf_melliger_RRS_no#_MEX_PU | Nat. His. Museum of Los Angeles County | LACMENT346506 | SAMN15830305 | | van Elst et al. 2021 |
| M_sp_cf_melliger_WPEM_17561_USA_TX | Uni. of Texas El Paso Biodiv. Collections | UTEP:Ento:25421 | SAMN15830306 | | van Elst et al. 2021 |
| M_sp_cf_melliger_WPM_5148_MEX_CH | Uni. of Texas El Paso Biodiv. Collections | UTEP:Ento:25424 | SAMN15830307 | | van Elst et al. 2021 |
| M_sp_cf_mendax-01_JDH_no#_USA_TX | Uni. of Texas Biodiv. Collections: Entomology | UTIC204184 | SAMN15830311 | | van Elst et al. 2021 |
| M_placodops_SS_no#_USA_CA | Arizona State Uni. Social Insect Biodiv. Rep. | ASU-SIBR00001176 | SAMN15830313 | | van Elst et al. 2021 |
| M_semirufus_TEBS_Ca-01_USA_CA | Arizona State Uni. Social Insect Biodiv. Rep. | ASU-SIBR00001042 | SAMN15830292 | | van Elst et al. 2021 |
| M_semirufus_TEBS_Ca-02_USA_CA | Arizona State Uni. Social Insect Biodiv. Rep. | ASU-SIBR00001043 | SAMN15830293 | | van Elst et al. 2021 |
| M_sp_cf_melliger_JT_Bue-01_MEX_SO | Arizona State Uni. Social Insect Biodiv. Rep. | ASU-SIBR00001053 | SAMN15830301 | | van Elst et al. 2021 |
| M_sp_cf_melliger_JT_Bue-09_MEX_SO | Arizona State Uni. Social Insect Biodiv. Rep. | ASU-SIBR00001054 | SAMN15830302 | | van Elst et al. 2021 |
| M_sp_cf_melliger_JT_Hop-11_USA_AZ | Arizona State Uni. Social Insect Biodiv. Rep. | ASU-SIBR00001055 | SAMN15830303 | | van Elst et al. 2021 |
| M_sp_cf_melliger_RAJ_5588_MEX_SO | Arizona State Uni. Social Insect Biodiv. Rep. | ASU-SIBR00001056 | SAMN15830304 | | van Elst et al. 2021 |
| M_sp_cf_mendax-01_FMH_1305_USA_TX | Uni. of Texas Biodiv. Collections: Entomology | UTIC204216 | SAMN15830309 | | van Elst et al. 2021 |
| M_sp_cf_mendax-01_FMH_no#_USA_TX | Uni. of Texas El Paso Biodiv. Collections | UTEP:Ento:25422 | SAMN15830310 | | van Elst et al. 2021 |
| M_sp_cf_mendax-01_WPM_13384_USA_TX | Uni. of Texas El Paso Biodiv. Collections | UTEP:Ento:25423 | SAMN15830312 | | van Elst et al. 2021 |
| M_sp_cf_mendax-03_JT_Fre-02_USA_AZ | Arizona State Uni. Social Insect Biodiv. Rep. | ASU-SIBR00001063 | SAMN15830314 | | van Elst et al. 2021 |
| M_sp_cf_mendax-03_JT_Hop-03_USA_AZ | Arizona State Uni. Social Insect Biodiv. Rep. | ASU-SIBR00001064 | SAMN15830315 | | van Elst et al. 2021 |
| M_sp_cf_mendax-03_JT_Mol-02_USA_AZ | Arizona State Uni. Social Insect Biodiv. Rep. | ASU-SIBR00001068 | SAMN15830316 | | van Elst et al. 2021 |
| M_sp_cf_mendax-03_JT_Mol-03_USA_AZ | Arizona State Uni. Social Insect Biodiv. Rep. | ASU-SIBR00001069 | SAMN15830317 | | van Elst et al. 2021 |
| M_sp_cf_mendax-03_JT_Ord-03_USA_AZ | Arizona State Uni. Social Insect Biodiv. Rep. | ASU-SIBR00001070 | SAMN15830318 | | van Elst et al. 2021 |
| M_sp_cf_mendax-03_JT_Ord-05_USA_AZ | Arizona State Uni. Social Insect Biodiv. Rep. | ASU-SIBR00001071 | SAMN15830319 | | van Elst et al. 2021 |
| M_sp_cf_mendax-03_JT_Ord-08_USA_AZ | Arizona State Uni. Social Insect Biodiv. Rep. | ASU-SIBR00001072 | SAMN15830320 | | van Elst et al. 2021 |
| M_sp_cf_mendax-03_JT_Syc-03_USA_AZ | Arizona State Uni. Social Insect Biodiv. Rep. | ASU-SIBR00001091 | SAMN15830321 | | van Elst et al. 2021 |
| M_sp_cf_mendax-03_KRW_no#_MEX_SO | Arizona State Uni. Social Insect Biodiv. Rep. | ASU-SIBR00001177 | SAMN15830322 | | van Elst et al. 2021 |
| M_sp_cf_mendax-03_MH_Sup-03_USA_AZ | Arizona State Uni. Social Insect Biodiv. Rep. | ASU-SIBR00001090 | SAMN15830323 | | van Elst et al. 2021 |
| M_sp_cf_mendax-03_PSW_14509_USA_AZ | Arizona State Uni. Social Insect Biodiv. Rep. | ASU-SIBR00001178 | SAMN15830324 | | van Elst et al. 2021 |
| M_sp_cf_mendax-03_TE_Don-01_USA_AZ | Arizona State Uni. Social Insect Biodiv. Rep. | ASU-SIBR00001061 | SAMN15830325 | | van Elst et al. 2021 |
| M_sp_cf_mendax-03_TE_Dri-01_USA_AZ | Arizona State Uni. Social Insect Biodiv. Rep. | ASU-SIBR00001062 | SAMN15830326 | | van Elst et al. 2021 |
| M_sp_cf_mendax-03_TE_Pat-01_USA_AZ | Arizona State Uni. Social Insect Biodiv. Rep. | ASU-SIBR00001073 | SAMN15830327 | | van Elst et al. 2021 |
| M_sp_cf_mendax-03_TE_Pat-08_USA_AZ | Arizona State Uni. Social Insect Biodiv. Rep. | ASU-SIBR00001074 | SAMN15830328 | | van Elst et al. 2021 |
| M_sp_cf_mendax-03_TE_Pat-09_USA_AZ | Arizona State Uni. Social Insect Biodiv. Rep. | ASU-SIBR00001075 | SAMN15830329 | | van Elst et al. 2021 |
| M_sp_cf_mendax-03_TE_Pcr-01_USA_AZ | Arizona State Uni. Social Insect Biodiv. Rep. | ASU-SIBR00001076 | SAMN15830330 | | van Elst et al. 2021 |
| M_sp_cf_mendax-03_TE_Pcr-10_USA_AZ | Arizona State Uni. Social Insect Biodiv. Rep. | ASU-SIBR00001077 | SAMN15830331 | | van Elst et al. 2021 |
| M_sp_cf_mendax-03_TE_Pin-04_USA_AZ | Arizona State Uni. Social Insect Biodiv. Rep. | ASU-SIBR00001078 | SAMN15830332 | | van Elst et al. 2021 |
| M_sp_cf_mendax-03_TE_Pin-07_USA_AZ | Arizona State Uni. Social Insect Biodiv. Rep. | ASU-SIBR00001079 | SAMN15830333 | | van Elst et al. 2021 |
| M_sp_cf_mendax-03_TE_Pin-14_USA_AZ | Arizona State Uni. Social Insect Biodiv. Rep. | ASU-SIBR00001080 | SAMN15830334 | | van Elst et al. 2021 |
| M_sp_cf_mendax-03_TE_Sch-01_USA_AZ | Arizona State Uni. Social Insect Biodiv. Rep. | ASU-SIBR00001081 | SAMN15830335 | | van Elst et al. 2021 |
| M_sp_cf_mendax-03_TE_Sch-07_USA_AZ | Arizona State Uni. Social Insect Biodiv. Rep. | ASU-SIBR00001082 | SAMN15830336 | | van Elst et al. 2021 |
| M_sp_cf_mendax-03_TE_Sch-11_USA_AZ | Arizona State Uni. Social Insect Biodiv. Rep. | ASU-SIBR00001083 | SAMN15830337 | | van Elst et al. 2021 |
| M_sp_cf_mendax-03_TE_Sie-05_USA_AZ | Arizona State Uni. Social Insect Biodiv. Rep. | ASU-SIBR00001084 | SAMN15830338 | | van Elst et al. 2021 |
| M_sp_cf_mendax-03_TE_Sie-126_USA_AZ | Arizona State Uni. Social Insect Biodiv. Rep. | ASU-SIBR00001085 | SAMN15830339 | | van Elst et al. 2021 |
| M_sp_cf_mendax-03_TE_Sie-143_USA_AZ | Arizona State Uni. Social Insect Biodiv. Rep. | ASU-SIBR00001086 | SAMN15830340 | | van Elst et al. 2021 |
| M_sp_cf_mendax-03_TE_Sie-151_USA_AZ | Arizona State Uni. Social Insect Biodiv. Rep. | ASU-SIBR00001087 | SAMN15830341 | | van Elst et al. 2021 |
| M_sp_cf_mendax-03_TE_Sie-20_USA_AZ | Arizona State Uni. Social Insect Biodiv. Rep. | ASU-SIBR00001088 | SAMN15830342 | | van Elst et al. 2021 |
| M_sp_cf_mendax-03_TE_Sie-72_USA_AZ | Arizona State Uni. Social Insect Biodiv. Rep. | ASU-SIBR00001089 | SAMN15830343 | | van Elst et al. 2021 |
| M_sp_cf_mendax-03_TECO_Ced-01_USA_AZ | Arizona State Uni. Social Insect Biodiv. Rep. | ASU-SIBR00001059 | SAMN15830344 | | van Elst et al. 2021 |
| M_sp_cf_mendax-03_TECO_Ced-02_USA_AZ | Arizona State Uni. Social Insect Biodiv. Rep. | ASU-SIBR00001060 | SAMN15830345 | | van Elst et al. 2021 |
| M_sp_cf_mendax-04_JGLW_Chi-02_USA_AZ | Arizona State Uni. Social Insect Biodiv. Rep. | ASU-SIBR00001096 | SAMN15830346 | | van Elst et al. 2021 |
| M_sp_cf_mendax-04_JGLW_Chi-06_USA_AZ | Arizona State Uni. Social Insect Biodiv. Rep. | ASU-SIBR00001097 | SAMN15830347 | | van Elst et al. 2021 |

| | | | | | |
|---|---|---|---|---|---|
| M_sp_cf_mendax-04_JGLW_Chi-08_USA_AZ | Arizona State Uni. Social Insect Biodiv. Rep. | ASU-SIBR00001098 | SAMN15830348 | | van Elst et al. 2021 |
| M_sp_cf_mendax-04_JT_Bue-02_MEX_SO | Arizona State Uni. Social Insect Biodiv. Rep. | ASU-SIBR00001092 | SAMN15830349 | | van Elst et al. 2021 |
| M_sp_cf_mendax-04_JT_Bue-04_MEX_SO | Arizona State Uni. Social Insect Biodiv. Rep. | ASU-SIBR00001093 | SAMN15830350 | | van Elst et al. 2021 |
| M_sp_cf_mendax-04_JT_Bue-11_MEX_SO | Arizona State Uni. Social Insect Biodiv. Rep. | ASU-SIBR00001094 | SAMN15830351 | | van Elst et al. 2021 |
| M_sp_cf_mendax-04_JT_Car-01_USA_AZ | Arizona State Uni. Social Insect Biodiv. Rep. | ASU-SIBR00001095 | SAMN15830352 | | van Elst et al. 2021 |
| M_sp_cf_mendax-04_JT_Coc-02_USA_AZ | Arizona State Uni. Social Insect Biodiv. Rep. | ASU-SIBR00001101 | SAMN15830353 | | van Elst et al. 2021 |
| M_sp_cf_mendax-04_JT_Coc-03_USA_AZ | Arizona State Uni. Social Insect Biodiv. Rep. | ASU-SIBR00001102 | SAMN15830354 | | van Elst et al. 2021 |
| M_sp_cf_mendax-04_JT_Coc-10_USA_AZ | Arizona State Uni. Social Insect Biodiv. Rep. | ASU-SIBR00001103 | SAMN15830355 | | van Elst et al. 2021 |
| M_sp_cf_mendax-04_JT_Fre-04_USA_AZ | Arizona State Uni. Social Insect Biodiv. Rep. | ASU-SIBR00001104 | SAMN15830356 | | van Elst et al. 2021 |
| M_sp_cf_mendax-04_JT_Fre-05_USA_AZ | Arizona State Uni. Social Insect Biodiv. Rep. | ASU-SIBR00001105 | SAMN15830357 | | van Elst et al. 2021 |
| M_sp_cf_mendax-04_JT_Kit-01_USA_AZ | Arizona State Uni. Social Insect Biodiv. Rep. | ASU-SIBR00001106 | SAMN15830358 | | van Elst et al. 2021 |
| M_sp_cf_mendax-04_JT_Mil-01_USA_AZ | Arizona State Uni. Social Insect Biodiv. Rep. | ASU-SIBR00001107 | SAMN15830359 | | van Elst et al. 2021 |
| M_sp_cf_mendax-04_JT_Ram-01_USA_AZ | Arizona State Uni. Social Insect Biodiv. Rep. | ASU-SIBR00001120 | SAMN15830360 | | van Elst et al. 2021 |
| M_sp_cf_mendax-04_PSW_16919-01_MEX_CH | Arizona State Uni. Social Insect Biodiv. Rep. | ASU-SIBR00001180 | SAMN15830362 | | van Elst et al. 2021 |
| M_sp_cf_mendax-04_RAJ_2536_MEX_CH | Arizona State Uni. Social Insect Biodiv. Rep. | ASU-SIBR00001117 | SAMN15830363 | | van Elst et al. 2021 |
| M_sp_cf_mendax-04_RAJ_2800_MEX_CH | Arizona State Uni. Social Insect Biodiv. Rep. | ASU-SIBR00001118 | SAMN15830364 | | van Elst et al. 2021 |
| M_sp_cf_mendax-04_RAJ_4930_USA_AZ | Arizona State Uni. Social Insect Biodiv. Rep. | ASU-SIBR00001119 | SAMN15830365 | | van Elst et al. 2021 |
| M_sp_cf_mendax-04_TE_Chi-50_USA_AZ | Arizona State Uni. Social Insect Biodiv. Rep. | ASU-SIBR00001099 | SAMN15830366 | | van Elst et al. 2021 |
| M_sp_cf_mendax-04_TEBS_Co-01_USA_CO | Arizona State Uni. Social Insect Biodiv. Rep. | ASU-SIBR00001100 | SAMN15830367 | | van Elst et al. 2021 |
| M_sp_cf_mendax-04_TEBS_Nm-01_USA_NM | Arizona State Uni. Social Insect Biodiv. Rep. | ASU-SIBR00001111 | SAMN15830368 | | van Elst et al. 2021 |
| M_sp_cf_mendax-04_TEBS_Nm-02_USA_NM | Arizona State Uni. Social Insect Biodiv. Rep. | ASU-SIBR00001112 | SAMN15830369 | | van Elst et al. 2021 |
| M_sp_cf_mendax-04_TEBS_Nm-03_USA_NM | Arizona State Uni. Social Insect Biodiv. Rep. | ASU-SIBR00001113 | SAMN15830370 | | van Elst et al. 2021 |
| M_sp_cf_mendax-04_TEBS_Nm-04_USA_NM | Arizona State Uni. Social Insect Biodiv. Rep. | ASU-SIBR00001114 | SAMN15830371 | | van Elst et al. 2021 |
| M_sp_cf_mendax-04_TEBS_Nm-05_USA_NM | Arizona State Uni. Social Insect Biodiv. Rep. | ASU-SIBR00001115 | SAMN15830372 | | van Elst et al. 2021 |
| M_sp_cf_mendax-04_TEBS_Nm-08_USA_NM | Arizona State Uni. Social Insect Biodiv. Rep. | ASU-SIBR00001116 | SAMN15830373 | | van Elst et al. 2021 |
| M_sp_cf_mendax-04_TEJGJT_Mor-01_USA_AZ | Arizona State Uni. Social Insect Biodiv. Rep. | ASU-SIBR00001108 | SAMN15830374 | | van Elst et al. 2021 |
| M_sp_cf_mendax-04_TEJGJT_Mor-06_USA_AZ | Arizona State Uni. Social Insect Biodiv. Rep. | ASU-SIBR00001109 | SAMN15830375 | | van Elst et al. 2021 |
| M_sp_cf_mendax-04_TEJGJT_Mor-08_USA_AZ | Arizona State Uni. Social Insect Biodiv. Rep. | ASU-SIBR00001110 | SAMN15830376 | | van Elst et al. 2021 |
| M_sp_cf_mendax-04_TEJGJT_Sky-03_USA_AZ | Arizona State Uni. Social Insect Biodiv. Rep. | ASU-SIBR00001121 | SAMN15830377 | | van Elst et al. 2021 |
| M_sp_cf_mendax-04_TEJGJT_Sky-06_USA_AZ | Arizona State Uni. Social Insect Biodiv. Rep. | ASU-SIBR00001122 | SAMN15830378 | | van Elst et al. 2021 |
| M_sp_cf_mendax-04_TEJGJT_Sky-13_USA_AZ | Arizona State Uni. Social Insect Biodiv. Rep. | ASU-SIBR00001123 | SAMN15830379 | | van Elst et al. 2021 |
| M_sp_cf_mendax-04_WPM_16472_USA_NM | Uni. of Texas El Paso Biodiv. Collections | UTEP:Ento:25425 | SAMN15830380 | | van Elst et al. 2021 |
| M_sp_cf_mendax-04_WPM_16704_USA_NM | Uni. of Texas El Paso Biodiv. Collections | UTEP:Ento:25427 | SAMN15830382 | | van Elst et al. 2021 |
| M_sp_cf_mendax-04_WPM_no#_USA_NM | Uni. of Texas El Paso Biodiv. Collections | UTEP:Ento:25429 | SAMN15830384 | | van Elst et al. 2021 |
| M_sp_cf_placodops-02_RRS_77-12_USA_CA | Nat. His. Museum of Los Angeles County | LACMENT346507 | SAMN15830412 | | van Elst et al. 2021 |
| M_sp_cf_placodops-02_TEBS_Nv-02_USA_NV | Arizona State Uni. Social Insect Biodiv. Rep. | ASU-SIBR00001136 | SAMN15830414 | | van Elst et al. 2021 |
| M_sp_cf_placodops-02_TEBS_Nv-03_USA_NV | Arizona State Uni. Social Insect Biodiv. Rep. | ASU-SIBR00001137 | SAMN15830415 | | van Elst et al. 2021 |
| M_sp_cf_placodops-03_TEBS_Nv-01_USA_NV | Arizona State Uni. Social Insect Biodiv. Rep. | ASU-SIBR00001139 | SAMN15830419 | | van Elst et al. 2021 |
| M_sp_cf_placodops-03_TEBS_Nv-04_USA_NV | Arizona State Uni. Social Insect Biodiv. Rep. | ASU-SIBR00001140 | SAMN15830420 | | van Elst et al. 2021 |
| M_sp_cf_placodops-03_TEBS_Nv-05_USA_NV | Arizona State Uni. Social Insect Biodiv. Rep. | ASU-SIBR00001141 | SAMN15830421 | | van Elst et al. 2021 |
| M_sp_cf_mexicanus-01_RRS_69-298_USA_AZ | Nat. His. Museum of Los Angeles County | LACMENT346523 | SAMN15830386 | | van Elst et al. 2021 |
| M_sp_cf_mexicanus-02_MLB_1122_USA_AZ | Arizona State Uni. Social Insect Biodiv. Rep. | ASU-SIBR00001025 | SAMN15830387 | | van Elst et al. 2021 |
| M_sp_cf_mexicanus-02_MLB_421_USA_NV | Arizona State Uni. Social Insect Biodiv. Rep. | ASU-SIBR00001026 | SAMN15830388 | | van Elst et al. 2021 |
| M_sp_cf_mexicanus-02_RAJ_3061_MEX_BC | Arizona State Uni. Social Insect Biodiv. Rep. | ASU-SIBR00001027 | SAMN15830389 | | van Elst et al. 2021 |
| M_sp_cf_mexicanus-02_RAJ_6012_USA_CA | Arizona State Uni. Social Insect Biodiv. Rep. | ASU-SIBR00001028 | SAMN15830390 | | van Elst et al. 2021 |
| M_sp_cf_mexicanus-02_RAJ_6028_USA_CA | Arizona State Uni. Social Insect Biodiv. Rep. | ASU-SIBR00001029 | SAMN15830391 | | van Elst et al. 2021 |
| M_sp_cf_mexicanus-02_SPC_6256_USA_CA | Museum of Comparative Zoology | MCZ654633 | SAMN15830392 | | van Elst et al. 2021 |
| | Arizona State Uni. Social Insect Biodiv. Rep. | ASU-SIBR00001125 | NA | | van Elst et al. 2021 |
| M_sp_cf_mimicus-flaviceps-01_RAJ_3076_MEX_BC | Arizona State Uni. Social Insect Biodiv. Rep. | ASU-SIBR00001128 | SAMN15830396 | | van Elst et al. 2021 |
| M_sp_cf_mimicus-flaviceps-01_RRS_77-22_USA_CA | Nat. His. Museum of Los Angeles County | LACMENT346511 | SAMN15830398 | | van Elst et al. 2021 |
| M_sp_cf_mimicus-flaviceps-02_RAJ_2665_MEX_BCS | Arizona State Uni. Social Insect Biodiv. Rep. | ASU-SIBR00001130 | SAMN15830399 | | van Elst et al. 2021 |
| M_sp_cf_mimicus-flaviceps-02_RAJ_3030_MEX_BCS | Arizona State Uni. Social Insect Biodiv. Rep. | ASU-SIBR00001131 | SAMN15830400 | | van Elst et al. 2021 |
| M_sp_cf_mimicus-flaviceps-03_JT_Fre-01_USA_AZ | Arizona State Uni. Social Insect Biodiv. Rep. | ASU-SIBR00001132 | SAMN15830401 | | van Elst et al. 2021 |
| M_sp_cf_mimicus-flaviceps-03_RAJ_5784_USA_AZ | Arizona State Uni. Social Insect Biodiv. Rep. | ASU-SIBR00001134 | SAMN15830403 | | van Elst et al. 2021 |
| M_navajo_GCWJNW_NEV892_USA_NV | Nat. His. Museum of Los Angeles County | LACMENT346524 | SAMN15830266 | | van Elst et al. 2021 |
| M_navajo_RAJ_2914_USA_AZ | Arizona State Uni. Social Insect Biodiv. Rep. | ASU-SIBR00001030 | SAMN15830267 | | van Elst et al. 2021 |
| M_navajo_RAJ_2986_MEX_BC | Arizona State Uni. Social Insect Biodiv. Rep. | ASU-SIBR00001031 | SAMN15830268 | | van Elst et al. 2021 |
| M_navajo_RAJ_3101_MEX_BC | Arizona State Uni. Social Insect Biodiv. Rep. | ASU-SIBR00001032 | SAMN15830269 | | van Elst et al. 2021 |
| M_navajo_SPC_7476_USA_AZ | Museum of Comparative Zoology | MCZ655243 | SAMN15830271 | | van Elst et al. 2021 |
| M_nequazcatl_EMF_no#_MEX_SO | Nat. His. Museum of Los Angeles County | LACMENT346503 | SAMN15830272 | Paratype | van Elst et al. 2021 |

| | | | | | |
|---|---|---|---|---|---|
| M_nequazcatl_PSW_13493_MEX_SO | Arizona State Uni. Social Insect Biodiv. Rep. | ASU-SIBR00001169 | SAMN15830273 | | van Elst et al. 2021 |
| M_nequazcatl_RAJ_962_MEX_SO | Arizona State Uni. Social Insect Biodiv. Rep. | ASU-SIBR00001034 | SAMN15830274 | | van Elst et al. 2021 |
| M_nequazcatl_RAJ_SON95-26_MEX_SO | Arizona State Uni. Social Insect Biodiv. Rep. | ASU-SIBR00001035 | SAMN15830275 | | van Elst et al. 2021 |
| M_perimeces_EMFb_no#_MEX_BC | Nat. His. Museum of Los Angeles County | LACMENT346519 | SAMN15830276 | Paratype | van Elst et al. 2021 |
| M_sp_cf_mendax-01_AW_6127_USA_TX | Arizona State Uni. Social Insect Biodiv. Rep. | ASU-SIBR00001058 | SAMN15830308 | | van Elst et al. 2021 |
| M_sp_cf_mendax-04_PSW_13407-6_USA_NM | Arizona State Uni. Social Insect Biodiv. Rep. | ASU-SIBR00001179 | SAMN15830361 | | van Elst et al. 2021 |
| M_sp_cf_mendax-04_WPM_16702_USA_NM | Uni. of Texas El Paso Biodiv. Collections | UTEP:Ento:25426 | SAMN15830381 | | van Elst et al. 2021 |
| M_sp_cf_mendax-04_WPM_19359_USA_CO | Uni. of Texas El Paso Biodiv. Collections | UTEP:Ento:25428 | SAMN15830383 | | van Elst et al. 2021 |
| M_sp_cf_placodops-01_ELVSDP_no#_USA_TX | Arizona State Uni. Social Insect Biodiv. Rep. | ASU-SIBR00001162 | SAMN15830406 | | van Elst et al. 2021 |
| M_sp_cf_placodops-01_PSW_15596_USA_TX | Arizona State Uni. Social Insect Biodiv. Rep. | ASU-SIBR00001182 | SAMN15830407 | | van Elst et al. 2021 |
| M_sp_cf_placodops-01_RWP_no#_USA_TX | Uni. of Texas Biodiv. Collections: Entomology | UTIC204391 | SAMN15830408 | | van Elst et al. 2021 |
| M_sp_cf_placodops-02_GCS_no#_USA_AZ | Nat. His. Museum of Los Angeles County | LACMENT346508 | SAMN15830409 | | van Elst et al. 2021 |
| M_sp_cf_placodops-02_KRW_no#_MEX_SO | Arizona State Uni. Social Insect Biodiv. Rep. | ASU-SIBR00001183 | SAMN15830410 | | van Elst et al. 2021 |
| M_sp_cf_placodops-02_PSW_16232_USA_AZ | Arizona State Uni. Social Insect Biodiv. Rep. | ASU-SIBR00001184 | SAMN15830411 | | van Elst et al. 2021 |
| M_sp_cf_placodops-02_TE_Sou-01_USA_AZ | Arizona State Uni. Social Insect Biodiv. Rep. | ASU-SIBR00001138 | SAMN15830413 | | van Elst et al. 2021 |
| M_sp_cf_placodops-03_PSW_14981-03_USA_AZ | Arizona State Uni. Social Insect Biodiv. Rep. | ASU-SIBR00001185 | SAMN15830416 | | van Elst et al. 2021 |
| M_sp_cf_placodops-03_PSW_15153_USA_CA | Arizona State Uni. Social Insect Biodiv. Rep. | ASU-SIBR00001186 | SAMN15830417 | | van Elst et al. 2021 |
| M_sp_cf_placodops-03_RAJ_2978_USA_AZ | Arizona State Uni. Social Insect Biodiv. Rep. | ASU-SIBR00001142 | SAMN15830418 | | van Elst et al. 2021 |
| M_pyramicus_GCWJNW_NEV1304_USA_NV | Nat. His. Museum of Los Angeles County | LACMENT346527 | SAMN15830277 | | van Elst et al. 2021 |
| M_pyramicus_PSW_16114_USA_CA | Arizona State Uni. Social Insect Biodiv. Rep. | ASU-SIBR00001170 | SAMN15830278 | | van Elst et al. 2021 |
| M_romainei_MAJO_71_USA_UT | Arizona State Uni. Social Insect Biodiv. Rep. | ASU-SIBR00001038 | SAMN15830279 | | van Elst et al. 2021 |
| M_romainei_MLB_1543_USA_AZ | Arizona State Uni. Social Insect Biodiv. Rep. | ASU-SIBR00001012 | SAMN15830280 | | van Elst et al. 2021 |
| M_romainei_PSW_15174_USA_CA | Arizona State Uni. Social Insect Biodiv. Rep. | ASU-SIBR00001171 | SAMN15830281 | | van Elst et al. 2021 |
| M_romainei_RAJ_6039_USA_CA | Arizona State Uni. Social Insect Biodiv. Rep. | ASU-SIBR00001041 | SAMN15830284 | | van Elst et al. 2021 |
| M_romainei_RRS_73-47_USA_TX | Nat. His. Museum of Los Angeles County | LACMENT346513 | SAMN15830285 | | van Elst et al. 2021 |
| M_romainei_SPC_6239_USA_AZ | Museum of Comparative Zoology | MCZ654622 | SAMN15830286 | | van Elst et al. 2021 |
| M_sp_cf_kennedyi-romainei_RAJ_6009_USA_CA | Arizona State Uni. Social Insect Biodiv. Rep. | ASU-SIBR00001040 | SAMN15830298 | | van Elst et al. 2021 |
| M_semirufus_CG_no#_USA_CA | Nat. His. Museum of Los Angeles County | LACMENT346509 | SAMN15830287 | | van Elst et al. 2021 |
| M_semirufus_PSW_14281_MEX_BC | Arizona State Uni. Social Insect Biodiv. Rep. | ASU-SIBR00001173 | SAMN15830288 | | van Elst et al. 2021 |
| M_semirufus_PSW_14810_USA_CA | Arizona State Uni. Social Insect Biodiv. Rep. | ASU-SIBR00001174 | SAMN15830289 | | van Elst et al. 2021 |
| M_semirufus_RAJ_2984_MEX_BC | Arizona State Uni. Social Insect Biodiv. Rep. | ASU-SIBR00001044 | SAMN15830290 | | van Elst et al. 2021 |
| M_semirufus_RAJ_BC1310_MEX_BC | Arizona State Uni. Social Insect Biodiv. Rep. | ASU-SIBR00001045 | SAMN15830291 | | van Elst et al. 2021 |
| M_flaviceps_RAJ_2958_USA_AZ | Arizona State Uni. Social Insect Biodiv. Rep. | ASU-SIBR00001014 | SAMN15830247 | | van Elst et al. 2021 |
| M_kennedyi_RAJ_2234_USA_AZ | Arizona State Uni. Social Insect Biodiv. Rep. | ASU-SIBR00001021 | SAMN15830259 | | van Elst et al. 2021 |
| M_navajo_RAJ_3139-2_USA_AZ | Arizona State Uni. Social Insect Biodiv. Rep. | ASU-SIBR00001033 | SAMN15830270 | | van Elst et al. 2021 |
| M_romainei_RAJ_2349_USA_AZ | Arizona State Uni. Social Insect Biodiv. Rep. | ASU-SIBR00001039 | SAMN15830283 | | van Elst et al. 2021 |
| M_yuma_RAJ_2243_USA_AZ | Arizona State Uni. Social Insect Biodiv. Rep. | ASU-SIBR00001155 | SAMN15830440 | | van Elst et al. 2021 |
| M_sp_SON-1_RAJ_5781_MEX_SO | Arizona State Uni. Social Insect Biodiv. Rep. | ASU-SIBR00001143 | SAMN15830422 | | van Elst et al. 2021 |
| M_sp_cf_colei_RAJ_2655_MEX_BC | Arizona State Uni. Social Insect Biodiv. Rep. | ASU-SIBR00001051 | SAMN15830294 | | van Elst et al. 2021 |
| M_sp_cf_mimicus-flaviceps-01_RAJ_2990_MEX_BC | Arizona State Uni. Social Insect Biodiv. Rep. | ASU-SIBR00001126 | SAMN15830394 | | van Elst et al. 2021 |
| M_sp_cf_mimicus-flaviceps-01_RAJ_3029_MEX_BC | Arizona State Uni. Social Insect Biodiv. Rep. | ASU-SIBR00001127 | SAMN15830395 | | van Elst et al. 2021 |
| M_sp_cf_kennedyi_RAJ_2810_USA_AZ | Arizona State Uni. Social Insect Biodiv. Rep. | ASU-SIBR00001052 | SAMN15830295 | | van Elst et al. 2021 |
| M_sp_cf_colei_RAJ_2723_MEX_BC | Arizona State Uni. Social Insect Biodiv. Rep. | ASU-SIBR00001057 | SAMN15830385 | | van Elst et al. 2021 |
| M_sp_cf_navajo_RAJ_3077_MEX_BC | Arizona State Uni. Social Insect Biodiv. Rep. | ASU-SIBR00001135 | SAMN15830405 | | van Elst et al. 2021 |
| M_tenuinodis_PSW_15178_USA_CA | Arizona State Uni. Social Insect Biodiv. Rep. | ASU-SIBR00001187 | SAMN15830423 | | van Elst et al. 2021 |
| M_tenuinodis_RRS_no#_USA_CA | Nat. His. Museum of Los Angeles County | LACMENT346520 | SAMN15830424 | Paratype | van Elst et al. 2021 |
| M_testaceus_MLB_1258_USA_CA | Arizona State Uni. Social Insect Biodiv. Rep. | ASU-SIBR00001144 | SAMN15830426 | | van Elst et al. 2021 |
| M_testaceus_MLB_397_USA_NV | Arizona State Uni. Social Insect Biodiv. Rep. | ASU-SIBR00001145 | SAMN15830427 | | van Elst et al. 2021 |
| M_testaceus_MLB_446_USA_CA | Arizona State Uni. Social Insect Biodiv. Rep. | ASU-SIBR00001146 | SAMN15830428 | | van Elst et al. 2021 |
| M_testaceus_RAJ_2259_MEX_BC | Arizona State Uni. Social Insect Biodiv. Rep. | ASU-SIBR00001147 | SAMN15830429 | | van Elst et al. 2021 |
| M_testaceus_RAJ_6018_USA_CA | Arizona State Uni. Social Insect Biodiv. Rep. | ASU-SIBR00001149 | SAMN15830430 | | van Elst et al. 2021 |
| M_testaceus_RRSCDG_77-20_USA_CA | Nat. His. Museum of Los Angeles County | LACMENT346528 | SAMN15830431 | | van Elst et al. 2021 |
| M_wheeleri_MLB_1256_USA_CA | Arizona State Uni. Social Insect Biodiv. Rep. | ASU-SIBR00001150 | SAMN15830432 | | van Elst et al. 2021 |
| M_wheeleri_PSW_14302_USA_CA | Arizona State Uni. Social Insect Biodiv. Rep. | ASU-SIBR00001188 | SAMN15830433 | | van Elst et al. 2021 |
| M_wheeleri_RAJ_3094_MEX_BC | Arizona State Uni. Social Insect Biodiv. Rep. | ASU-SIBR00001151 | SAMN15830434 | | van Elst et al. 2021 |
| M_wheeleri_RAJ_6010_USA_CA | Arizona State Uni. Social Insect Biodiv. Rep. | ASU-SIBR00001152 | SAMN15830435 | | van Elst et al. 2021 |
| M_wheeleri_RAJ_6013_USA_CA | Arizona State Uni. Social Insect Biodiv. Rep. | ASU-SIBR00001153 | SAMN15830436 | | van Elst et al. 2021 |
| M_wheeleri_RAJ_6021_USA_CA | Arizona State Uni. Social Insect Biodiv. Rep. | ASU-SIBR00001154 | SAMN15830437 | | van Elst et al. 2021 |
| M_wheeleri_RRS_no#_USA_CA | Nat. His. Museum of Los Angeles County | LACMENT346504 | SAMN15830438 | Paratype | van Elst et al. 2021 |
| M_tenuinodis_SPC_4826_USA_AZ | Museum of Comparative Zoology | MCZ661684 | SAMN15830425 | | van Elst et al. 2021 |
| M_yuma_PSW_15185_USA_CA | Arizona State Uni. Social Insect Biodiv. Rep. | ASU-SIBR00001189 | SAMN15830439 | | van Elst et al. 2021 |

| | | | | | |
|---|---|---|---|---|---|
| M_yuma_RAJ_3084_MEX_BC | Arizona State Uni. Social Insect Biodiv. Rep. | ASU-SIBR00001156 | SAMN15830441 | | van Elst et al. 2021 |
| M_yuma_RAJ_5783_MEX_SO | Arizona State Uni. Social Insect Biodiv. Rep. | ASU-SIBR00001158 | SAMN15830442 | | van Elst et al. 2021 |
| M_yuma_RRS_94-11a_USA_CA | Nat. His. Museum of Los Angeles County | LACMENT346521 | SAMN15830443 | | van Elst et al. 2021 |
| Myrmelachista_joycei_JTL_8492_CR | John T. Longino Collection | CASENT0635767 | NA | Outgroup | Branstetter et al., 2017 |
| Nylanderia_terricola_JTL_14763_USA_TX | Arizona State Uni. Social Insect Biodiv. Rep. | ASU-SIBR00001160 | NA | Outgroup | Messer et al., unpubl. |
| Paratrechina_longicornis_LRD_270806-14_USA_FL | NA | NA | NA | Outgroup | Messer et al., unpubl. |
| Polyergus_mexicanus_MLB_389-01_USA_CA | Arizona State Uni. Social Insect Biodiv. Rep. | ASU-SIBR00001161 | NA | Outgroup | Borowiec et al., unpubl. |
